# Effectiveness and equity of vaccination strategies against Rift Valley fever in a heterogeneous landscape

**DOI:** 10.1101/2024.07.18.604096

**Authors:** Warren S. D. Tennant, Eric Cardinale, Youssouf Moutroifi, Simon E. F. Spencer, Onzade Charafouddine, Mike J. Tildesley, Raphaëlle Métras

## Abstract

Spatio-temporal variations in environment and socio-agricultural factors create heterogeneity in livestock disease transmission risk, raising challenges in identifying populations most at risk and how this risk changes over time. Consequently, prioritising control strategies, such as vaccination, to achieve optimal or equitable outcomes across regions impedes the design of an effective vaccination strategy. We developed a metapopulation model for Rift Valley fever transmission in livestock across the Comoros archipelago which incorporates livestock vaccination in addition to heterogeneity in viral transmission rates and animal movements. We used the model to evaluate three vaccine allocation strategies–proportional allocation, optimal allocation for maximising total infections averted across the archipelago, and optimal allocation for more equitable outcomes across islands—under different vaccination coverage levels and animal identification scenarios. We report that (i) both archipelago-wide and island-specific strategy effectiveness were impacted by vaccination rate, allocation strategy, and animal identification approach, (ii) optimally allocating vaccines improved strategy effectiveness compared with proportional allocation but resulted in inequitable outcomes between islands, and (iii) tagging animals post-vaccination boosted overall strategy effectiveness for all vaccination rates.

## Introduction

Designing effective vaccination programs against livestock disease is challenged by spatio-temporal heterogeneities in the environmental and socio-agricultural factors that influence disease emergence, spread and persistence. For many infectious livestock diseases, such as foot and mouth disease, Rift Valley fever (RVF) and *Peste des Petit Ruminants*, variation in disease transmission across space and time may be attributed to spatially clustered distributions of livestock [1, 2], trade of livestock between affected regions [3, 4], and differences in livestock species composition and susceptibility to disease [5, 6]. For vector-borne zoonotic diseases, such as RVF, Bluetongue and Lumpy Skin disease, additional factors like the variation in the abundance and competence of different vector species to transmit the virus [7, 8], seasonal fluctuations in vector populations [9, 10], and the presence of wildlife reservoirs that can act as disease hosts [11, 12], further complicate and limit our understanding of disease transmission. As a result it is not immediately clear which populations are most at risk of infection and how this changes across time, raising difficulties in how to prioritise control strategies, such as vaccination, to achieve the most effective or equitable outcomes across or between regions.

Mathematical models are increasingly used for designing effective vaccination programs against livestock diseases. These modelling approaches typically incorporate information on host demography, disease transmission mechanisms and properties of the vaccine. They may also be fitted to epidemiological data, allowing an analysis into how different vaccine strategies would perform in past or future forecasted outbreaks (for example, see [13–15]). However, previous modelling approaches for vector-borne livestock diseases, such as RVF, have focused on evaluating the impact of vaccination in theoretical settings [16–18] or spatially homogeneous regions [14, 19, 20], preventing a formal quantitative assessment of how vaccines should be allocated across a spatially heterogeneous region to achieve optimal or equitable epidemiological outcomes.

In this paper, we utilised the Comoros archipelago, a network of four islands in the southwestern Indian Ocean, as a case study to assess optimal vaccine allocation strategies against RVF while considering spatio-temporal heterogeneity in transmission and animal movement between regions. We extended a mathematical model that simulates livestock transmission dynamics of RVF in the Comoros archipelago [21] to incorporate a description of vaccine allocation across islands. The model was used to evaluate the effectiveness of three vaccine allocation strategies possible to implement on the field: (i) proportional allocation based on island livestock population sizes, (ii) optimal allocation to maximise total infections averted across the archipelago, and (iii) optimal allocation to achieve more equitable epidemiological outcomes between islands. We further explored the impacts of tagging livestock post-vaccination to track their vaccine history and investigated the effects of targeting different age groups, frequency, and timings of vaccine administration.

## Results

The purpose of this study was to evaluate the effectiveness of different vaccine strategies against Rift Valley fever virus (RVFV). A spatially explicit, time-dependent mathematical model was developed to describe transmission of RVF in livestock across islands in the Comoros archipelago, incorporating a description of vaccination against infection Figure 1. The model was used to investigate the effects of vaccination in the Comoros archipelago on epidemiological outcomes in livestock by estimating optimal strategies for allocation of vaccines between islands, across feasible vaccination rates, and livestock tagging strategies.

**Figure 1:**
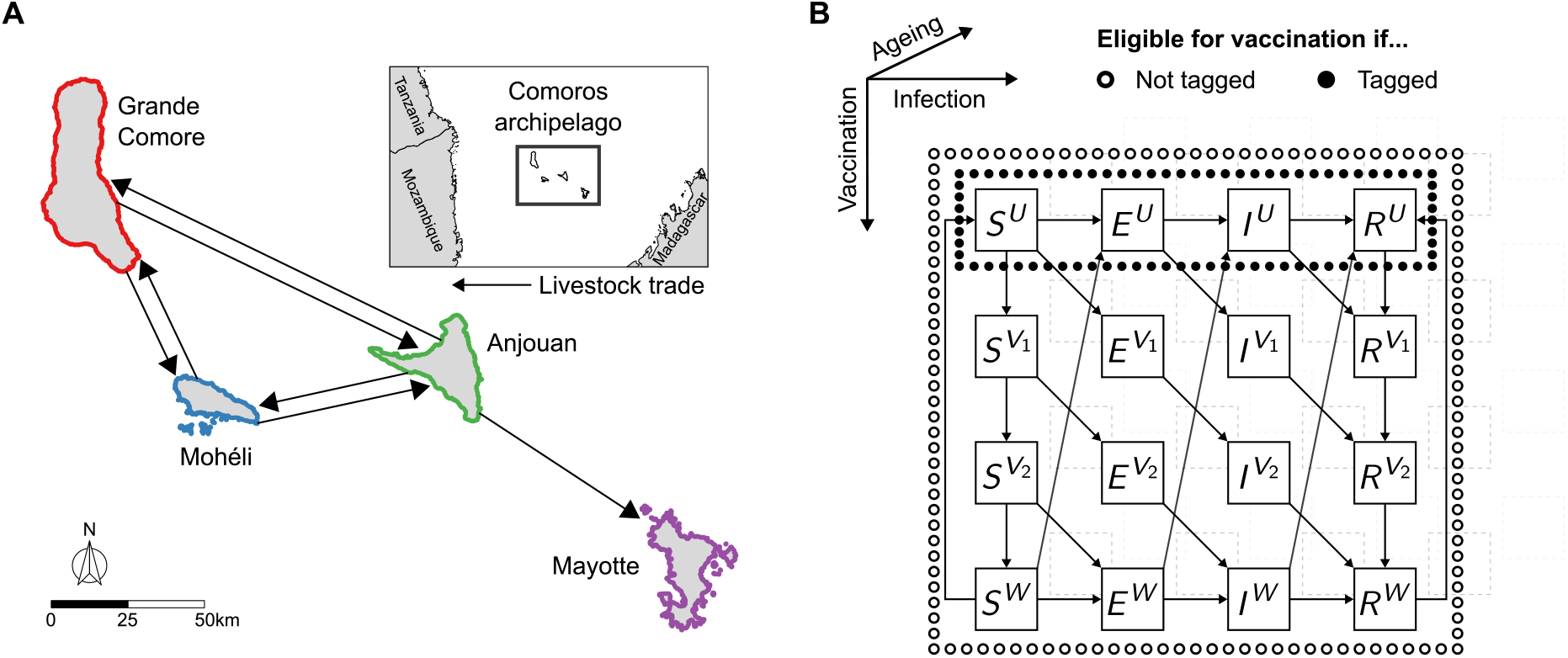
Schematic outlining the inter- and intra-island processes of livestock in the mathematical model. To evaluate the effect of vaccine control measures against Rift Valley fever virus (RVFV), we developed a mathematical model to describe RVFV infection in livestock (cattle, sheep and goats). **(A)** In the model, animals moved between the four islands in the Comoros archipelago—Grande Comore (red), Mohéli (blue), Anjouan (green) and Mayotte (purple)—as governed by the livestock trade network as shown. **(B)** The livestock population of each island was divided into compartments defined by infection status, vaccination status, and age. The diagram shows the compartments (squares) and the direction of transfer of livestock between compartments (arrows). The compartments eligible for vaccination depended on whether or not livestock were tagged (black circles) or not (white circles). The notation in each compartment define the infection status (horizontally arranged) of the animals: susceptible to the virus (*S*), infected but not yet infectious (*E*), infectious (*I*) or recovered with life-long immunity to reinfection (*R*). The superscripts denote the vaccination status (vertically arranged) of animals: unvaccinated (*U*), vaccinated with developing protection (*V*_1_ and *V*_2_), or vaccinated and partially protected from infection (*W*). The transitions between compartments of different age groups (diagonally arranged) are not drawn for illustrative purposes.

### Estimating vaccine allocation across the archipelago

Given a number of vaccines to administer annually throughout the archipelago, vaccines were allocated to each island in three different ways: (i) proportional to the livestock population size of each island, (ii) optimally to maximise infections averted in livestock across the archipelago, and (iii) optimally to maximise the infections averted on the worst-performing island—defined as the island with the lowest percentage of infections averted in livestock. Whilst the second allocation approach aimed for an optimal archipelago-wide outcome, the third allocation ensured that all islands would perform at least as well as the worst-performing island, therefore accounting for equity between island-specific outcomes. The last two allocations were estimated using an optimisation algorithm (see Methods section), which incorporated uncertainty in island-specific RVFV transmission potential and livestock movement between islands. The estimation of each vaccine allocation are shown in Figure 2, and detailed results are presented below.

**Figure 2:**
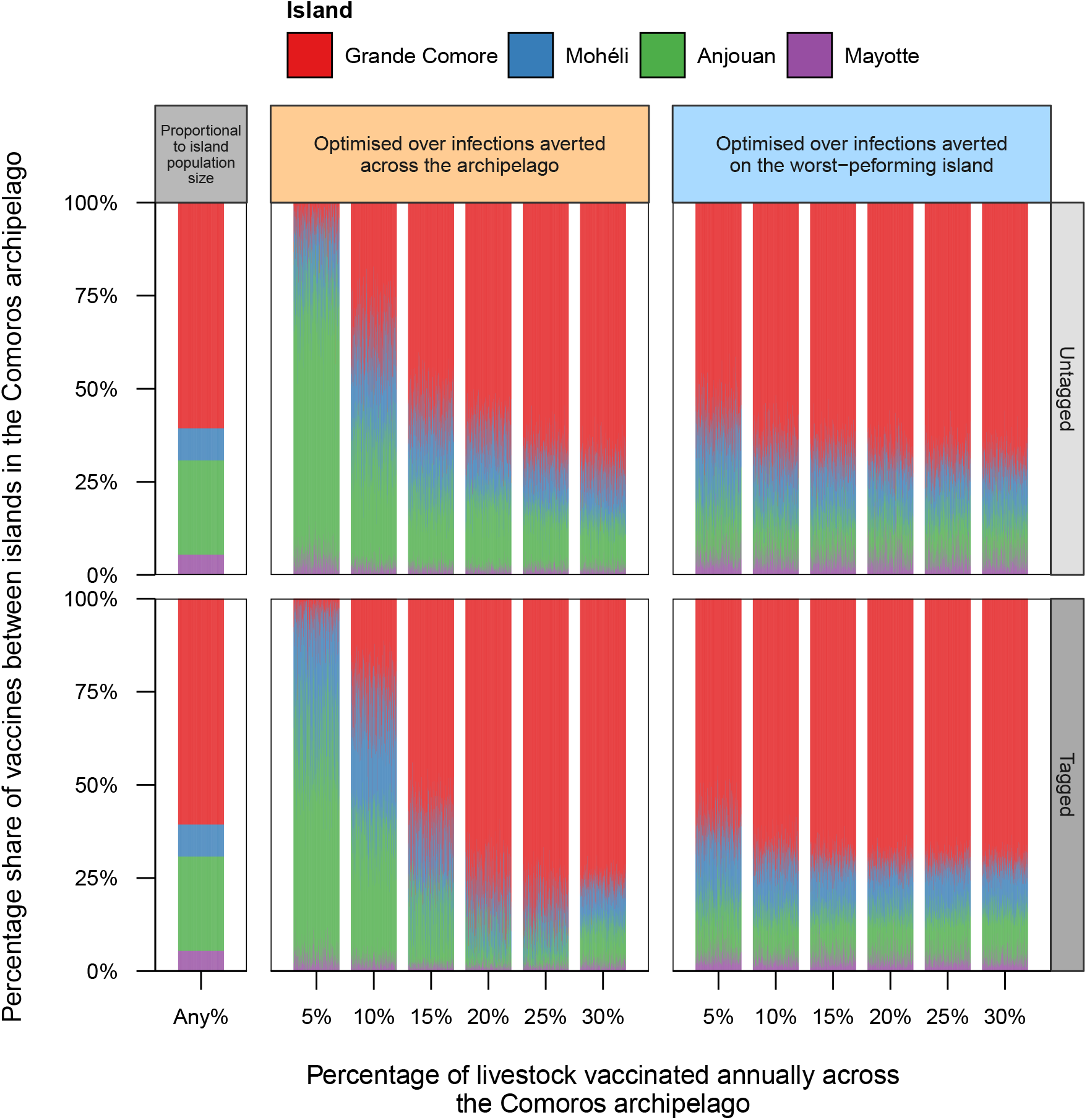
Vaccine allocation strategies across islands in the Comoros archipelago. Vaccines were allocated to each of the four islands in the archipelago—Grande Comore (red), Mohéli (blue), Anjouan (green) and Mayotte (purple)—either proportionally to the livestock population size of each island (left-most panel), or optimally to maximise the percentage of infections averted across the archipelago (middle panel), or the percentage of infections averted on the island with the worst performance (rightmost panel). Optimally allocating vaccines based on the percentage of infections averted across the archipelago was dependent on the vaccination rate, exhibiting a shift from allocating the majority of vaccines to Anjouan for lower vaccination rates, to Grande Comore for higher vaccination rates. The vertical bars show the percentage of vaccines assigned to each island for different vaccination rates and tagging strategies. Refer to Supplementary Table 1 for detailed numerical results.

### Proportional to livestock population size

By allocating vaccines proportional to the population size of each island, 60.7%, 8.6%, 25.3% and 5.4% of vaccines were allocated to Grande Comore, Mohéli, Anjouan and Mayotte respectively on an annual basis (Figure 2; grey, left-most panels). This vaccine allocation was independent of any vaccination rate or livestock tagging strategy.

### Optimal outcomes across the archipelago

When optimising in terms of percentage of infections averted across the archipelago, the allocation of vaccines was largely dependent on the overall vaccination rate (Figure 2; orange, middle panels), and resulted in the majority of vaccines assigned to Anjouan and Grande Comore for low and high vaccination rates respectively. These allocations were robust to both tagging strategies. For example, under the untagged scenario, vaccinating 5% of livestock across the archipelago annually favoured Anjouan, receiving a median of 71.7% (95% credible interval (CrI) = [55.4, 87.2]) of vaccines, followed by Mohéli with 16.0% (95% CrI = [2.7, 33.9]), Grande Comore with 6.4% (95% CrI = [0.2, 20.9]), and then Mayotte with 4.0% (95% CrI = [0.3, 11.1]) of vaccines. In contrast, a 30% annual vaccine coverage resulted in assigning a median of 71.9% (95% CrI = [62.6, 82.4]) of vaccines to Grande Comore, 13.8% (95% CrI = [4.5, 20.6]) to Mohéli, 13.02% (95% CrI = [7.3, 18.3]) to Anjouan and 1.52% (95% CrI = [0.1, 5.2]) to Mayotte. Similar vaccine allocations were found when animals were tagged post-vaccination. Vaccinating 5% of livestock annually in the archipelago, islands receiving vaccines from the most to the least percentage of vaccines was Anjouan (with a median of 62.8%), Mohéli (28.3%), Grande Comore (4.1%) and Mayotte (3.1%). Refer to Supplementary Table 1 for the median and 95% credible intervals of the percentage of vaccines assigned to each island for all combinations of tested vaccine allocations, tagging strategies and vaccination rates.

### Optimal outcomes on the worst-performing island

In contrast to optimising in terms of infections averted across the entire archipelago, optimising vaccine allocation in terms of the percentage of infections averted on the island with the worst-performance was robust to both vaccination rate and tagging strategy (Figure 2; blue, right-most panels). With an annual archipelago-wide vaccination rate of 5%, the median (and 95% CrI) of the percentage of vaccines assigned to Grande Comore, Mohéli, Anjouan and Mayotte was 59.8% [47.1, 73.1], 18.7% [10.8, 33.3], 14.4% [6.4, 27.3], and 4.4% [1.1, 14.9], respectively, for untagged livestock; and 62.6% [49.7, 72.7], 18.1% [11.4, 30.3], 14.1% [7.1, 24.0], respectively, assuming livestock were tagged. These corresponded to a greater percentage of livestock on Grande Comore and Mohéli being vaccinated than on Anjouan and Mayotte across different vaccination rates on average. For example, vaccinating 30% of livestock in the archipelago annually and assuming animals were untagged, the median percentage of livestock vaccinated on Grande Comore and Mohéli annually was 35.1% and 43.2% respectively, whereas only 12.2% and 22.5% of livestock were vaccinated on Anjouan and Mayotte. Refer to Supplementary Figure 1 and Supplementary Table 2 for the percentage of livestock vaccinated annually on each island for all combinations of tested vaccine allocations, tagging strategies and vaccination rates.

### Assessing the effectiveness of vaccination across the archipelago

Using the three vaccine allocations as described above, the model was simulated for 35 years and compared with the model without vaccination to establish the effectiveness of a long-term continuous vaccine campaign. Figure 3 shows the percentage of infections averted over the archipelago under the different vaccination strategies. Refer to Supplementary Figures 2 to 5 and Supplementary Tables 3 and 4 for the model predicted number of infections across the archipelago with and without vaccination.

**Figure 3:**
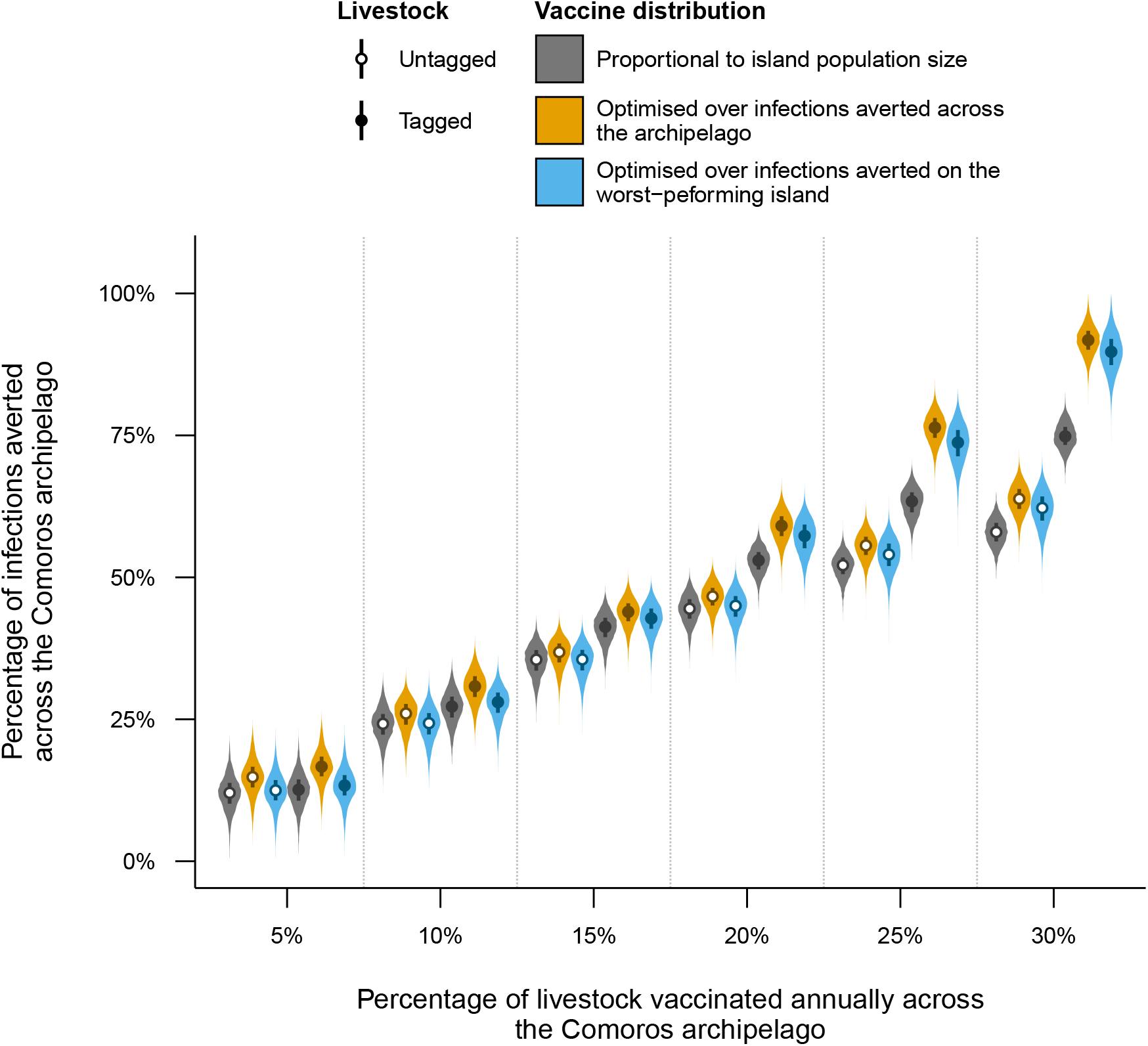
Effectiveness of different vaccine strategies against Rift Valley fever virus (RVFV) across the Comoros archipelago. For a range of vaccination rates, allocating vaccines to the four islands in the Comoros archipelago optimally by percentage of infections averted across the archipelago (orange) and by percentage of infections averted on the worst-performing island (blue; where the worst-performing island was defined as the island with the lowest percentage of infections averted) outperformed allocating vaccines proportionally to the population size of each island (grey). Tagging animals upon vaccination (black circles) also outperformed strategies where animals were not tagged (white circles), where the difference was amplified for increased vaccination rates. The violins show the percentage of infections averted across the Comoros archipelago for different annual vaccination rates, allocation methods and tagging strategies. The points and boxplots show the median and inter-quartile range for each scenario respectively. All metrics shown were based on 25,000 model simulations.

Vaccinating a greater percentage of livestock annually resulted in a greater percentage of infections averted across the archipelago for all vaccine strategies. Allocating vaccines proportional to the population size of each island and not tagging livestock post-vaccination resulted in a median of 12.0% (95% prediction interval (PI) = [5.9, 17.6]), 24.2% (95% PrI = [18.0, 29.5]), 35.5% (95% PrI = [29.4, 40.0]), 44.5% (95% PrI = [38.9, 48.9]), 52.1% (95% PrI = [47.2, 56.0]), 58.0% (95% PrI = [53.2, 62.2]) of infections averted when vaccinating 5%, 10%, 15%, 20%, 25% and 30% of livestock annually across the archipelago, respectively.

### Proportional versus optimal vaccine allocation

Optimally allocating vaccines between islands (Figure 3; orange and blue violins) outperformed allocating vaccines proportionally to population size (Figure 3; grey violins) for all vaccination rates and tagging strategies, with the advantage of optimal vaccine allocations increasing as the vaccination rate increased. For example, at 5% vaccination and with untagged livestock, the optimal outcome allocation averted a median of 14.8% (95% PrI = [8.5, 20.8]) infections compared to the 12.0% of infections with proportional allocation. This difference widened up to a vaccination rate of 30%, with optimal and proportional vaccine allocations averting 63.8% (95% PrI = [58.5, 68.5]) and 58.0% of infections respectively. The percentage of infections averted across the archipelago with the archipelago-wide optimal vaccine allocation (Figure 3; orange violins) consistently outperformed the equitable outcome vaccine allocation (Figure 3; blue violins) in terms of infections averted across the archipelago.

### Tagging livestock post-vaccination

Livestock tagging post-vaccination increased the percentage of infections averted across the archipelago when compared with strategies where livestock were not tagged, for all vaccine allocations and vaccination rates, but became more pronounced as vaccination rates increased. For instance, at the largest vaccination rate (30%) tagging animals averted 44% more infections than not tagging animals, specifically averting a median of 62.2% (95% PrI = [55.4, 67.6]) and 89.7% (95% PrI = [82.71, 96.1]) of infections across the archipelago in the untagged and tagged scenarios, respectively. Model predictions of infections averted across the archipelago for all vaccination rates, vaccine allocations and tagging strategies can be found in Supplementary Figures 6 to 8 and Supplementary Table 4.

### Vaccine administration efficiency

In addition to increasing infections averted, tagging livestock post-vaccination improved the efficiency of vaccine administration, defined as the proportion of vaccines administrated to susceptible livestock only (i.e. not to livestock with vaccine-induced nor natural protection against RVFV), as tagging ensured a greater percentage of vaccines were administered to animals that did not have any prior protection against RVFV infection. For example, the median efficiency across the archipelago was 66.9% (95% PrI = [65.5, 68.1]) and 95.8 (95% PrI = [94.2, 97.3]) under 30% annual vaccine coverage for the untagged and tagged scenarios, respectively. However, in all strategies the proportion of livestock with natural immunity to the virus would increase following an outbreak, resulting in a drop in vaccine administration efficiency on each island (Supplementary Figures 9 to 11). Refer to Supplementary Table 5 the median and 95% prediction interval of the (mean) administration efficiency on each island for each vaccine strategy.

### Impact of vaccinating the young, at the start of the year and vaccination frequency

Using the estimated optimal vaccine allocations for each vaccination rate and tagging scenario, the percentage of infections averted across the Comoros archipelago was simulated for scenarios which vaccinated only the youngest two age groups, vaccinated only in the first month of the epidemiological year (July) and only vaccinated every two or three years. Vaccinating only the two youngest age groups and/or only in the first month of the year was similar to vaccinating all age groups and throughout the year (Supplementary Figure 12). However, vaccinating less frequently resulted in a lower percentage of infections averted across the Comoros archipelago (Supplementary Figure 13). This result was consistent between vaccination rates; when comparing vaccinating 10% of livestock every year versus 30% of untagged livestock every three years, for example, which averted 26% (95% PrI = [20, 30]) and 22% (95% PrI = [16, 27]) of infections, respectively. Refer to Supplementary Figures 12 and 13 for detailed numerical results on each scenario.

### Evaluating the equity of island-specific strategy effectiveness

To assess equity of outcomes between islands due to vaccination, the percentage of infections averted on each island was measured for all vaccination rates, vaccine allocations and tagging strategies as described above. Figure 4 shows the median and 95% prediction interval of infections averted on each island in the Comoros archipelago with both optimal vaccine allocations when vaccinating 5% and 30% of the total archipelago livestock population annually. Refer to Supplementary Figures 14 to 16 and Supplementary Tables 6 and 7 for the model predicted number of infections and percentage of infections averted per island for all vaccination rates, vaccine allocations and tagging strategies.

**Figure 4:**
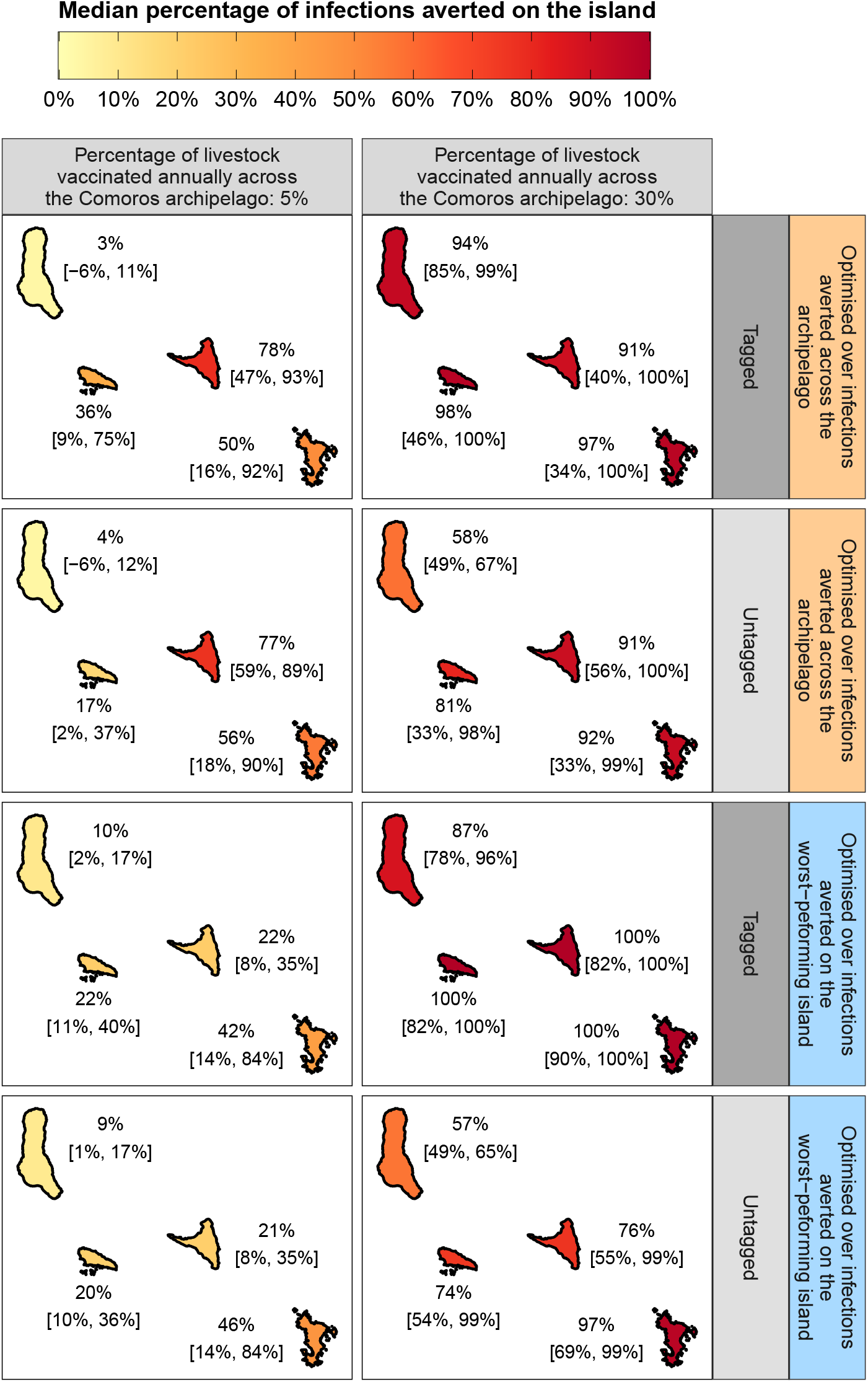
Equity in vaccine strategy effectiveness between islands in the Comoros archipelago. Optimally allocating vaccines in terms of total infections averted across the archipelago (orange panels) resulted in an imbalance of vaccine strategy effectiveness at the island-level at low vaccination rates. To ameliorate this, vaccines were also optimally allocated in terms of infections averted on the worst performing island (blue panels; where the worst-performing island was defined as the island with the lowest percentage of infections averted), resulting in a less pronounced imbalance of strategy effectiveness between islands at low vaccination rates. Shown is the median and 95% prediction interval of the percentage of infections averted on each island when vaccinating 5% and 30% of livestock across the archipelago annually under each optimal vaccine allocation and tagging strategy. Summary statistics were generated using 25,000 model simulations. See Supplementary Tables 6 and 7 for detailed numerical results.

### Imbalance in strategy effectiveness between islands at low vaccination rates

Vaccines allocated according to maximising the number of infections averted across the archipelago lead to a discrepancy in the percentage of infections averted between islands, particularly at lower vaccination rates: vaccinating 5% of untagged livestock annually across the archipelago averted 4% (95% PrI = [−6, 12]) of infections on Grande Comore, 17% (95% PrI = [2, 37]) on Mohéli, 77% (95% PrI = [59, 89]) on Anjouan and 56% (95% PrI = [18, 90]) on Mayotte. For this strategy, it was noted that Grande Comore had the potential to avert a negative number of infections. That is, vaccination resulted in a greater number of model predicted infections than without vaccination. This result is due to vaccination delaying seasonal outbreaks of RVFV to subsequent years where virus transmission was higher because of more suitable environmental conditions, resulting in larger outbreaks (see Supplementary Figure 6, for example).

### Allocating vaccines to achieve greater equity in epidemiological outcomes

Optimally allocating vaccines in terms of infections averted on the worst-performing island partially addressed the imbalance in strategy effectiveness at low vaccination rates. For example, for the same scenario (5% vaccination rate and untagged livestock), 9% (95% PrI = [1, 17]), 20% (95% PrI = [10, 36]), 21% (95% PrI = [8, 35]) and 46% (95% PrI = [14, 84]) of infections were averted on Grande Comore, Mohéli, Anjouan and Mayotte, respectively. The imbalance in infections averted on each island was also ameliorated in this manner for vaccine strategies where livestock were tagged post-vaccination. Detailed differences in the percentage of infections averted on each island are provided in Supplementary Figures 17 to 19 and Supplementary Table 8.

## Discussion

Spatio-temporal heterogeneities in environmental and socio-agricultural factors present a significant challenge to developing effective vaccination strategies against infectious livestock diseases. These challenges are further amplified for vector-borne zoonotic diseases, such as RVF, where uncertainties about the role of vectors and wildlife reservoirs on disease transmission add an extra layer of complexity. Addi-tionally, geopolitical factors can introduce pressure to balance disease control outcomes between regions while achieving maximum effectiveness globally. Given these challenges, quantifying the effectiveness of a range of vaccine allocations in a heterogeneous landscape is crucial. Therefore our study employed a mathematical modeling approach to investigate three vaccine allocation strategies for mitigating RVFV transmission in the Comoros archipelago, a network of four islands in the southwestern Indian Ocean. Our model explicitly considers the spatial and temporal variations and uncertainty in disease transmission potential within each island, as well as the movement of livestock between islands.

Our findings suggest that (i) allocation of vaccines proportional to livestock population sizes did not lead to the most effective overall control strategy against RVFV, (ii) optimising the allocation of vaccines to achieve maximum strategy effectiveness resulted in an imbalance of strategy effectiveness at the island-level, and (iii) tracking the vaccination status of livestock substantially increased strategy effectiveness. Finally, we provided evidence that administering vaccines shortly after an outbreak was highly inefficient.

Previous modeling studies of RVF have primarily focused on mass vaccination campaigns in theoretical settings or spatially homogeneous regions, with outcomes measured over short timeframes [20, 19, 16, 14, 18]. Here, we evaluated the effectiveness of vaccine strategies against RVF in the context of a preventive, continuous, long-term campaign. We measured strategy effectiveness in terms of model-predicted livestock infections averted. With this approach, equitably allocating vaccines between the four islands in the Comoros archipelago was less effective than allocating vaccines to achieve optimal strategy effectiveness. This was because the equitable allocation does not take into account the different transmission potential of the virus on each island. This optimal allocation was sensitive to the number of vaccines administered annually. With low vaccination rates, allocating vaccines towards Anjouan, the island with the second-lowest transmission potential but second-largest livestock population, was more effective than allocating to Grande Comore, the island with the largest transmission potential and livestock population size. This finding aligns with previous work demonstrating that focusing vaccine efforts towards a single population may be the most optimal approach with limited vaccine supplies [22]. However, such an approach may raise ethical concerns, as it could lead to significantly beneficial outcomes for one region over another.

In our study, optimally allocating vaccines did result in some islands experiencing a significantly lower percentage of infections averted compared to others. This was the case for all vaccination rates. To address this, we developed an approach for maintaining high overall strategy effectiveness while achieving equitable outcomes across all islands. We optimised vaccine allocation to ensure that all islands performed at least as well as the island with the worst average performance. This approach not only improved the balance in strategy effectiveness between islands but also maintained an effective outcome globally (across the islands), outperforming equitable resource allocation on average. However, it is important to note that even with this approach, the equitable-outcome allocation did not achieve purely equal outcomes across all islands. This is likely due to the spatio-temporal heterogeneity in disease transmission and host susceptibility, which can lead to a unique set of possible qualitative outcomes for each island: frequent outbreaks (endemic), sporadic outbreaks (epidemic), or an absence of outbreaks (elimination). Therefore, it is unlikely that any vaccine allocation strategy can achieve perfectly equal epidemiological outcomes (in finite time) between regions with heterogeneous epidemiological drivers.

In contrast to the optimal outcome vaccine allocation, the equitable outcome allocation was less sensitive to changes in livestock vaccination rate. This suggests that an effective and equitable vaccine strategy covering regions with spatio-temporal heterogeneity in disease transmission may be devised prior to knowing how many vaccines are available, an ongoing challenge in resource limited settings [23–25]. However, even if vaccines were able to be mobilised during an epidemic effectively, our results further highlight that it may be inefficient to vaccinate shortly after an outbreak if reactive measures are the priority, and that vaccinating with very limited supply may result in worse epidemiological outcomes compared to if no vaccines were administered. Our proposal for a continuous prophylactic vaccine campaign in a resource-limited setting may motivate a reliable and sufficiently large, albeit limited, supply of vaccines ready for deployment. In addition to availability, there are other significant barriers preventing equitable access to livestock vaccines, including acceptability, affordability and accessibility [26, 25]. Some vaccines may also have existing standards on when they are administered (e.g. prior to the rainy season). Our findings suggest that strategy effectiveness may be robust to changes in the timing of vaccine administration provided it is annual. There may be some strategies to improve access and acceptance of vaccines through the involvement of donors and international organisations [27], however we acknowledge that there are many social barriers to effective vaccine deployment.

Our work highlighted that tagging livestock in order to track their vaccine history may be beneficial for improving overall strategy effectiveness. Tagging livestock post-vaccination limits vaccine wastage— that is, the model highlights that focusing vaccine efforts on livestock who do not have any vaccine-induced protection against infection is beneficial not only from the perspective of the number of infections averted in livestock, but may thus reduce the cost, or demand on supply, of vaccines to achieve the same epidemiological outcomes. The benefits of animal identification on the reduction of livestock disease burden has been previously highlighted, e.g. for *Peste des Petits Ruminants* [13], and the tagging strategy presented here emphasises that tracking which and when livestock have been vaccinated can focus vaccine efforts towards unprotected animals. However, the assumption here is that it is (perfectly) known when vaccine-induced immunity in vaccinated livestock has waned. In reality this may only be known through serological testing, alongside a DIVA-compliant vaccine [28], to delineate whether an animal has any natural or vaccine-induced protection. A full cost-benefit analysis would be needed to determine whether it is more economically efficient for all livestock to be sampled and serologically tested for protection against RVFV prior to administration of the vaccine, or otherwise. Tagging livestock, as well as increasing overall strategy effectiveness, could help to further build the picture of the livestock trade network, which may help to address control targets for other livestock diseases, such as FMD.

While our study sheds light on optimal vaccine allocation for RVF control in the Comoros archipelago, some limitations in the modelling approach are important to consider. Firstly, we explicitly assumed that future trends in RVFV transmission potential over space and time (as governed by the Normalised Difference Vegetation Index) was identical to that of the 2004–2021 period. This implies that climate effects on mosquito demography and the resulting changes in transmission rate are similar to the past when projecting forward vaccine strategies until 2050. This may not be the case given that mosquito demography is sensitive to rainfall, temperature and humidity [29–31]. However, climate forecasting was beyond the scope of our study, and instead this motivates further development of our approach to consider adaptive surveillance and control measures, whereby the optimal control strategy may be updated over time, taking into account possible changes in recent environmental predictions as well as socio-agricultural patterns (e.g. temporary movement bans). On the spatial scale, we have assumed that livestock mix homogeneously within each island. Consequently, our results imply an equitable allocation of vaccines within each island itself. This may not only be impractical, but a better approach would be to target areas of each island that are at higher risk of RVFV transmission. Although there exists an abundance of mathematical models which describe livestock disease transmission at fine spatial scales, extensive within-island serological information at a finer spatial scale would be required to parameterise such a model in the manner we have done in our study, and consequently evaluate the effectiveness of within-island vaccine allocation strategies. Finally, we defined strategy effectiveness purely in terms of the model-predicted percentage of infections averted. In reality, a policy-maker’s evaluation of a vaccine strategy may be multi-faceted, considering not only epidemiological benefits, but also social and economical ones. As a result, the definition of an effective vaccine strategy may have multiple objectives, leading to additional trade-offs between benefits and costs, where there may not always be a clear objective optimal approach [16, 32].

In conclusion, our study employed a mathematical modelling approach to investigate optimal vaccine allocation strategies for mitigating Rift Valley Fever virus transmission using the Comoros archipelago as a case study. We explicitly considered the spatio-temporal heterogeneities in environmental and socio-ecological factors, demonstrating that equitable allocation across all islands may not be the most effective approach. Striving to achieve optimal outcomes over a spatially heterogeneous region may result in inequity between regions, highlighting the inherent tension between equity and effectiveness in resource allocation. However, we also identified an approach that can address some of the imbalances between competing objectives. Finally, tracking the vaccination status of livestock through tagging, thereby prioritising vaccination of susceptible animals can significantly improve epidemiological outcomes. Our study highlights the importance of considering spatio-temporal heterogeneities and trade-offs between equity and effectiveness when designing vaccine allocation strategies for livestock diseases like Rift Valley fever.

## Methods

### Mathematical model

To assess the effectiveness of different vaccination strategies, we built on a previous metapopulation model describing the spread of Rift Valley fever virus (RVFV) infection in livestock across the Comoros archipelago [21] by including vaccination against RVFV infection.

### Demography and infection

We modelled the livestock population (cattle, sheep and goats) as in the previous model [21]. To summarise, we used a metapopulation framework of *n* patches (for the Comoros archipelago, *n* = 4), where each patch in the metapopulation is herein referred to as an island. In the model, livestock move between each island via a livestock trade network (Figure 1A), and within each island, the livestock population at each time *t* is subdivided into *A* age groups. Each age group is further divided by their infection status, namely susceptible (with compartment denoted *S*), exposed (*E*), infectious (*I*) or recovered (*R*). The proportion that animals moved from S to E was modelled as a exponential function of Normalised Difference Vegetation Index (NDVI). Refer to Supplementary Methods 1 for a detailed description of the demographic and infection processes and Supplementary Table 9 for all notation used in the mathematical model.

### Vaccination

Vaccination was incorporated into the model as follows, and a schematic summarising the model is shown in Figure 1.

The vaccination status of animals is either unprotected (with compartment denoted with a superscript of *U*), one week post-vaccination (*V*_1_), two weeks post-vaccination (*V*_2_) or well-protected (*W*). With time, island and age group denoted using subscripts *t, i* and *a* respectively, the number of individuals at time *t*, island *i*, age group *a*, infection status 𝒳 and vaccination status 𝒱 is denoted by 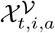. For example, the number of susceptible individuals that are not protected with a vaccine at time *t*, on island *i* and of age group *a* is denoted by 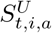.

The number of vaccine doses to administer across the metapopulation per week, *ψ*, is distributed to each island *i* as a proportion *ρ*_*i*_. Livestock are vaccinated irrespective of their infection history, and only animals in age groups up to and including age group *A*^vac^ were vaccinated. The rate at which livestock are vaccinated depended on whether livestock are assumed to be tagged or not. If all livestock are tagged, then the vaccination history of an animal is known, and so the vaccine was only administered to animals that were unprotected (through vaccination), *U*. If all livestock are not tagged, then vaccines are administered to all livestock irrespective of their vaccination status. It is assumed that re-vaccination of livestock which are developing vaccine protection, *V*_1_ or *V*_2_, or vaccine-protected, *W*, does not alter their current vaccination status. The weekly proportion of an animal on island *i* of age group *a* being vaccinated at time *t* is

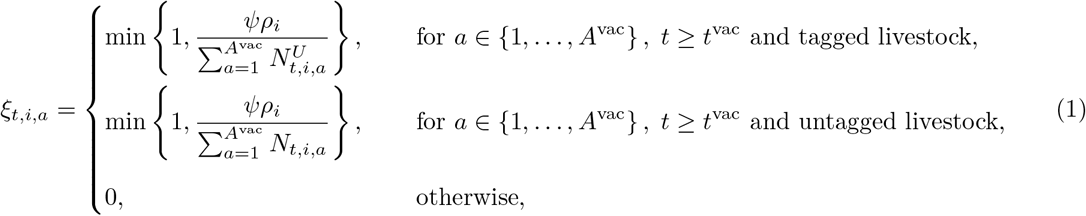

where *A*^vac^ denotes the maximum age group that vaccines were administered to, and 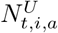 denotes the total unvaccinated population of age group *a* at time *t* on island *i*.

Livestock that are vaccinated took two weeks to develop vaccine-induced protection. These compartments are denoted by *V*_1_ and *V*_2_. During this time, livestock are still infected at the same force of infection as unprotected livestock. Livestock are partially protected from infection two weeks after vaccination. These livestock are infected at a proportion (1 − *p*^eff^) of the rate that unprotected livestock are infected, where *p*^eff^ denotes the efficacy of the vaccine.

In the model, vaccine-induced protection waned with weekly proportion *ω*. Livestock whose immunity wanes are assumed to be unprotected and eligible for re-vaccination. The weekly probability that vaccine-induced immunity waned is given by

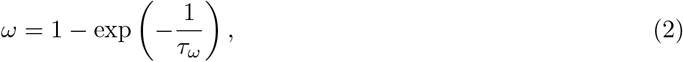

where *τ*_*ω*_ denotes the mean number of weeks a vaccinated and protected individual had vaccine-induced immunity to the virus.

### Parameterisation

Using the Comoros archipelago as a case study to evaluate the impact of different vaccine strategies against RVFV, the number of islands in the metapopulation was set to four (*n* = 4). The demographic and infection processes were parameterised as in the previous model [21] and can be found in Supplementary Methods 1.

Vaccine efficacy, *p*_eff_, and duration of protection, *τ*_*ω*_, within the model was primarily parameterised from field studies on vaccines in cattle, sheep and goats [33–37]. Njenga et al. [36] found 97% and 91% immunogenicity in sheep and goats respectively, and 67% immunogenicity in cattle at 14 days post-vaccination. For the purposes of this study to compare the effectiveness of a range of vaccination strategies against RVFV, and as the model does not delineate between cattle, sheep nor goats, the efficacy of the vaccine in the model was set at 90%. Trials typically follow livestock for up to two years [34–37] post-vaccination, exhibiting lasting protection for this duration. The duration of vaccine-induced protection was thus set at 2 years in the model.

The vaccine was introduced after the end of the model fitting period (*t*_*V*_ = 528), and all age groups were eligible for vaccination (*A*^*V*^ = 10)., and the primary results of the studied vaccinating all 10 age groups (*A*^*V*^ = 10). Six different vaccination rates, *ψ*, were simulated using the model, corresponding to vaccinating 5%, 10%, 15%, 20%, 25% and 30% of the livestock population across the Comoros archipelago annually. Three different approaches for allocating the vaccines to islands in the archipelago, *ρ*_*i*_, were used—(i) proportional to the population size of each island, (ii) optimally to maximise infections averted across the archipelago, and (iii) optimally to maximise the infections averted on the island with the worst overall performance. The latter two optimal approaches of allocating vaccines is described below.

### Optimisation of vaccine allocation

One goal of the study was to determine the optimal way to distribute vaccines across the archipelago. Given a number of vaccines to administer in the metapopulation per unit time, *ψ* as described above, vaccines were allocated to each island per unit time according to the vaccine distribution *ρ*. Here, we denote the complete set of possible vaccine allocations by *P*, where *P* defines an (*n* − 1) simplex:

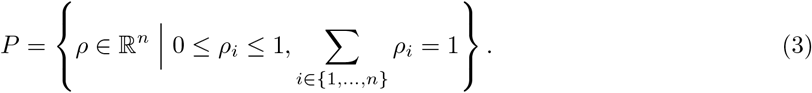

We sought to optimise this distribution of vaccines by maximising some objective function *f* that depended on *ρ*. We denote the optimal vaccine distribution with *ρ*^*^.

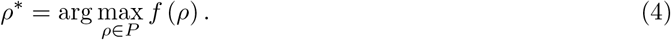

The objective function was defined in terms of the effectiveness of a given vaccine strategy, *g* as defined below, and included any uncertainty in model parameters, denoted by *θ* (refer to Supplementary Table 10 for parameter values and ranges). Integrating over the joint distribution of model parameters *π* (*θ*) yields the following objective function:

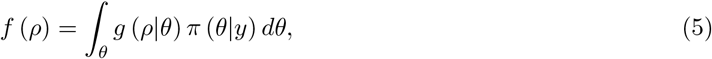

where *g* (*ρ*|*θ*) denotes the effectiveness of a given vaccine distribution *ρ* and model parameters *θ*, and *π* (*θ*|*y*) denotes the posterior distribution of the fitted model without vaccination as in Tennant et al. [21].

Direct analysis of the above objective function is intractable, therefore we used Monte Carlo integration to approximate it. The following equation gives an unbiased estimate of the objective function for any given vaccine distribution *ρ*:

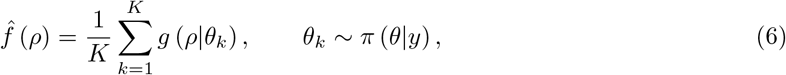

where *θ*_*k*_ denotes a single sample from the joint distribution of model parameters *π* (*θ*), *K* is the total number of random samples of model parameters, and *g* (*ρ*|*θ*_*k*_) represents the effectiveness of a vaccine distribution *ρ* given a single sample of model parameters *θ*_*k*_.

### Vaccine strategy effectiveness

We used two different definitions of vaccine strategy effectiveness functions, *g*_1_ and *g*_2_, to optimise the vaccine distribution across the metapopulation:

1. The first effectiveness function, *g*_1_, considered a collective approach where effectiveness was defined as the proportion of infections averted across the entire metapopulation with vaccination compared to without vaccination from the time the vaccine was first administered at *t*_v_ up to some time *T*.

2. The second effectiveness function, denoted as *g*_2_, considered an equitable approach where effectiveness was defined as the proportion of infections averted on the worst-performing island only; that is, the island with the least number of infections averted.

The equations describing the different objective functions are defined below, where the absence of any vaccination against RVFV is denoted by |_*ψ*=0_.

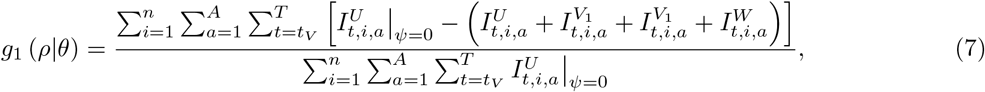

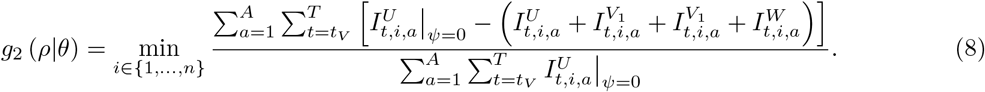

### Optimisation algorithm

To calculate the optimal distribution of vaccines across the metapopulation, we employed a sequential Monte Carlo (SMC) optimisation algorithm [38]. SMC algorithms are well-suited for this task because they allow us to explore potential multimodal objective functions, as may be the case here with a spacetime dependent model, which can be a challenge for gradient-based optimization methods. The optimisation algorithm worked by gradually concentrating a set of vaccine distributions, herein referred to as ‘particles’, into regions of parameter space associated with a larger evaluations of the objective function. The algorithm that we used consisted of three main components:

1. a sampling part, in which particles with a higher objective function evaluation were favoured more highly than others,
2. a density-tempering part, which allowed particles to move from the initially proposed distribution of particles towards an empirical distribution of the objective function, and
3. a rejuvenation part, which meant that sampled particles would be jittered using a Metropolis-Hastings kernel with a (truncated) multivariate Gaussian proposal distribution.

The particle which maximised the objective function determined the optimal particle. Refer to the Supplementary Methods 2 for a detailed description of each of the above components and pseudo-code for the optimisation algorithm.

### Vaccine strategies evaluated

This study aimed to assess the effectiveness of a range of long-term vaccination campaigns on disease burden in livestock. The effectiveness of a single vaccine strategy was calculated by first simulating the model described above forward in time using Supplementary Equations 5 to 41 where future disease transmission rates were computed using recycled Normalised Difference Vegetation Index (NDVI) from July 2004 until the end of June 2021. With this approach, the model was simulated with and without vaccination up to an equivalent time horizon of June 2050 (*T* = 2207). For each simulation, the following metrics were calculated: (i) the percentage of infections averted across the archipelago (see Equation 7), (ii) the percentage of infections averted on each island in the archipelago, and (iii) the vaccine efficiency which was defined as the percentage of vaccines that were administered to susceptible and unprotected livestock (the *S*^*U*^ compartment in the model).

Table 1 shows the list of vaccine strategies that were assessed in this work. The model was used to calculate the effectiveness of six vaccination strategies differing in their assumption on whether or not livestock were identifiable via tagging and vaccine allocation across the Comoros archipelago.

**Table 1:** Vaccine strategies evaluated using the mathematical model. The mathematical model describing infection with and vaccination against RVFV in livestock across the archipelago was used to assess the effectiveness of six vaccination strategies. These vaccine strategies differed in their assumption on whether or not livestock were identifiable via tagging, and how the vaccines were distributed between the four islands in the Comoros archipelago.

| Livestock identifiability | Vaccine distribution |
| --- | --- |
| Untagged | Proportional to the livestock population size of each island |
| Tagged | Proportional to the livestock population size of each island |
| Untagged | Optimised for the percentage of infections averted across the Comoros archipelago |
| Tagged | Optimised for the percentage of infections averted across the Comoros archipelago |
| Untagged | Optimised for the percentage of infections averted on the worst-performing island |
| Tagged | Optimised for the percentage of infections averted on the worst-performing island |

Three different vaccine allocations were used: (i) distributing vaccines proportional to the livestock population size of each island, (ii) optimally to maximise infections averted across the archipelago (see Equation 7), and (iii) optimally to maximise the infections averted on the island with the worst overall performance (see Equation 8). The optimisation algorithm was executed 500 times to calculate the latter two optimal allocations. Each vaccination strategy was evaluated with six different vaccination rates— 5%, 10%, 15%, 20%, 25% and 30%—defining the percentage of livestock vaccinated across the archipelago annually.

### Sensitivity analysis

Finally, for each vaccination rate, we used the optimal allocation of vaccines (in terms of maximised infections averted across the archipelago) and used the model to further simulate the impact of (i) only vaccinating the two youngest age groups, (ii) only vaccinating in the first week of each epidemiological year, and (iii) only vaccinating every two or three years.

### Computational implementation

The metapopulation model and optimisation algorithm were coded and executed in C++20 with the GNU Scientific Library (version 2.7) [39]. The outputs of these algorithms were analysed and visualised in R (version 4.3.2) [40] and the tidyverse library (version 2.0.0) [41].

## Data availability

The datasets used to simulate the mathematical model forward in time and execute the optimisation algorithm, alongside a full description of the data, are fully accessible through the GitHub repository: wtennant/rvf_vaccine [42].

## Code availability

All code, including the metapopulation model, optimisation algorithm, and data analysis scripts, and a walk through of how to replicate our analysis for one vaccine strategy presented in our study are publicly available through the GitHub repository: wtennant/rvf vaccine [42].

## Acknowledgements

This study was conducted under the project entitled ‘US-UK Collab: Adaptive surveillance and control for the elimination of endemic disease’. W.S.D.T. and M.J.T. were supported by the Biotechnology and Biological Sciences Research Council [grant number BB/T004312/1].

## Author contributions

W.S.D.T., E.C., M.J.T. and R.M. conceptualised the study design. W.S.D.T., M.J.T., S.E.F.S. and R.M. designed the mathematical model. W.S.D.T. implemented the code for the mathematical model and optimisation algorithm. W.S.D.T., E.C., Y.M., O.C. and R.M. defined the vaccine strategy scenarios to evaluate. W.S.D.T. and R.M. formally parameterised the mathematical model. W.S.D.T. performed the formal analysis of model and optimisation results. All authors interpreted the results. W.S.D.T. and R.M. wrote the first draft of the manuscript. All authors critically reviewed and edited the manuscript. All authors approved the final draft of the manuscript.

## Competing interests

The authors declare no competing interests.

## Notes

https://www.github.com/wtennant/rvf_vaccine

### Competing Interest Statement

The authors have declared no competing interest.

## References

[1] Nkamwesiga, J., Korennoy, F., Lumu, P., Nsamba, P., Mwiine, F. N., Roesel, K., Wieland, B., Perez, A., Kiara, H., and Muhanguzi, D. (2022). Spatio-temporal cluster analysis and transmission drivers for Peste des Petits Ruminants in Uganda. Transboundary and emerging diseases, 69(5):e1642–e1658.

[2] Tildesley, M. J., House, T. A., Bruhn, M. C., Curry, R. J., O’Neil, M., Allpress, J. L., Smith, G., and Keeling, M. J. (2010). Impact of spatial clustering on disease transmission and optimal control. Proceedings of the National Academy of Sciences, 107(3):1041–1046.

[3] Sumner, T., Orton, R. J., Green, D. M., Kao, R. R., and Gubbins, S. (2017). Quantifying the roles of host movement and vector dispersal in the transmission of vector-borne diseases of livestock. PLoS computational biology, 13(4):e1005470.

[4] Ekwem, D., Morrison, T. A., Reeve, R., Enright, J., Buza, J., Shirima, G., Mwajombe, J. K., Lembo, T., and Hopcraft, J. G. C. (2021). Livestock movement informs the risk of disease spread in traditional production systems in East Africa. Scientific Reports, 11(1):16375.

[5] Miller, R. S., Sweeney, S. J., Slootmaker, C., Grear, D. A., Di Salvo, P. A., Kiser, D., and Shwiff, S. A. (2017). Cross-species transmission potential between wild pigs, livestock, poultry, wildlife, and humans: implications for disease risk management in North America. Scientific Reports, 7(1):7821.

[6] Ostfeld, R. S. and Keesing, F. (2012). Effects of host diversity on infectious disease. Annual review of ecology, evolution, and systematics, 43:157–182.

[7] Turell, M. J., Linthicum, K. J., Patrican, L. A., Davies, F. G., Kairo, A., and Bailey, C. L. (2008). Vector competence of selected African mosquito (Diptera: Culicidae) species for Rift Valley fever virus. Journal of medical entomology, 45(1):102–108.

[8] Jean Jose Nepomichene, T.N., Elissa, N., Cardinale, E., and Boyer, S. (2015). Species diversity, abundance, and host preferences of mosquitoes (Diptera: Culicidae) in two different ecotypes of Madagascar with recent RVFV transmission. Journal of Medical Entomology, 52(5):962–969.

[9] Collins, Á.B., Mee, J. F., Doherty, M. L., Barrett, D. J., and England, M. E. (2018). Culicoides species composition and abundance on Irish cattle farms: implications for arboviral disease transmission. Parasites & vectors, 11:1–13.

[10] Charron, M. V., Kluiters, G., Langlais, M., Seegers, H., Baylis, M., and Ezanno, P. (2013). Seasonal and spatial heterogeneities in host and vector abundances impact the spatiotemporal spread of bluetongue. Veterinary Research, 44:1–12.

[11] Ruiz-Fons, F., Sánchez-Matamoros, A., Gortázar, C., and Sánchez-Vizcaíno, J. M. (2014). The role of wildlife in bluetongue virus maintenance in Europe: lessons learned after the natural infection in Spain. Virus Research, 182:50–58.

[12] Rostal, M. K., Liang, J. E., Zimmermann, D., Bengis, R., Paweska, J., and Karesh, W. B. (2017). Rift Valley fever: does wildlife play a role? Ilar Journal, 58(3):359–370.

[13] ElArbi, A. S., Kane, Y., Metras, R., Hammami, P., Ciss, M., Beye, A., Lancelot, R., Diallo, A., and Apolloni, A. (2019). PPR control in a Sahelian setting: what vaccination strategy for Mauritania? Frontiers in Veterinary Science, 6:242.

[14] Métras, R., Edmunds, W. J., Youssouffi, C., Dommergues, L., Fournié, G., Camacho, A., Funk, S., Cardinale, E., Le Godais, G., Combo, S., et al. (2020). Estimation of Rift Valley fever virus spillover to humans during the Mayotte 2018–2019 epidemic. Proceedings of the National Academy of Sciences, 117(39):24567–24574.

[15] Fournié, G., Waret-Szkuta, A., Camacho, A., Yigezu, L. M., Pfeiffer, D. U., and Roger, F. (2018). A dynamic model of transmission and elimination of peste des petits ruminants in Ethiopia. Proceedings of the National Academy of Sciences, 115(33):8454–8459.

[16] Adongo, D., Fister, K. R., Gaff, H., and Hartley, D. (2013). Optimal control applied to Rift Valley fever. Natural Resource Modeling, 26(3):385–402.

[17] Butt, A. I. K., Aftab, H., Imran, M., and Ismaeel, T. (2023). Mathematical study of lumpy skin disease with optimal control analysis through vaccination. Alexandria Engineering Journal, 72: 247–259.

[18] Falowo, O. D., Olaniyi, S., and Oladipo, A. T. (2023). Optimal control assessment of Rift Valley fever model with vaccination and environmental sanitation in the presence of treatment delay. Modeling Earth Systems and Environment, 9(1):457–471.

[19] Gachohi, J. M., Njenga, M. K., Kitala, P., and Bett, B. (2016). Modelling vaccination strategies against Rift Valley fever in livestock in Kenya. PLoS Neglected Tropical Diseases, 10(12):e0005049.

[20] Cecilia, H., Drouin, A., Métras, R., Balenghien, T., Durand, B., Chevalier, V., and Ezanno, P. (2022). Mechanistic models of Rift Valley fever virus transmission: A systematic review. PLoS Neglected Tropical Diseases, 16(11):e0010339.

[21] Tennant, W. S., Cardinale, E., Cêtre-Sossah, C., Moutroifi, Y., Le Godais, G., Colombi, D., Spencer, S. E., Tildesley, M. J., Keeling, M. J., Charafouddine, O., et al. (2021). Modelling the persistence and control of Rift Valley fever virus in a spatially heterogeneous landscape. Nature communications, 12(1):5593.

[22] Keeling, M. J. and Shattock, A. (2012). Optimal but unequitable prophylactic distribution of vaccine. Epidemics, 4(2):78–85.

[23] Alders, R., Bagnol, B., Young, M., Ahlers, C., Brum, E., Rushton, J., et al. (2007). Challenges and constraints to vaccination in developing countries. Developments in biologicals, 130(8):73–82.

[24] Lewis, C. E. and Roth, J. A. (2021). Challenges in having vaccines available to control transboundary diseases of livestock. Current Issues in Molecular Biology, 42(1):1–40.

[25] Nuvey, F. S., Fink, G., Hattendorf, J., Mensah, G. I., Addo, K. K., Bonfoh, B., and Zinsstag, J. (2023). Access to vaccination services for priority ruminant livestock diseases in Ghana: Barriers and determinants of service utilization by farmers. Preventive Veterinary Medicine, 215:105919.

[26] Acosta, D., Ludgate, N., McKune, S. L., and Russo, S. (2022). Who has access to livestock vaccines? Using the social-ecological model and intersectionality frameworks to identify the social barriers to peste des petits ruminants vaccines in Karamoja, Uganda. Frontiers in Veterinary Science, 9:831752.

[27] Donadeu, M., Nwankpa, N., Abela-Ridder, B., and Dungu, B. (2019). Strategies to increase adoption of animal vaccines by smallholder farmers with focus on neglected diseases and marginalized populations. PLoS Neglected Tropical Diseases, 13(2):e0006989.

[28] Van Oirschot, J. (1999). Diva vaccines that reduce virus transmission. Journal of biotechnology, 73 (2-3):195–205.

[29] Turell, M. (1989). Effect of environmental temperature on the vector competence of Aedes fowleri for Rift Valley fever virus. Research in Virology, 140:147–154.

[30] Brubaker, J. F. and Turell, M. J. (1998). Effect of environmental temperature on the susceptibility of Culex pipiens (Diptera: Culicidae) to Rift Valley fever virus. Journal of medical entomology, 35 (6):918–921.

[31] Sang, R., Lutomiah, J., Said, M., Makio, A., Koka, H., Koskei, E., Nyunja, A., Owaka, S., Matoke-Muhia, D., Bukachi, S., et al. (2017). Effects of irrigation and rainfall on the population dynamics of rift valley fever and other arbovirus mosquito vectors in the epidemic-prone Tana River County, Kenya. Journal of medical entomology, 54(2):460–470.

[32] Bolzoni, L., Bonacini, E., Della Marca, R., and Groppi, M. (2019). Optimal control of epidemic size and duration with limited resources. Mathematical biosciences, 315:108232.

[33] Kitandwe, P. K., McKay, P. F., Kaleebu, P., and Shattock, R. J. (2022). An overview of Rift Valley fever vaccine development strategies. Vaccines, 10(11):1794.

[34] Dungu, B., Louw, I., Lubisi, A., Hunter, P., von Teichman, B. F., and Bouloy, M. (2010). Evaluation of the efficacy and safety of the Rift Valley Fever Clone 13 vaccine in sheep. Vaccine, 28(29):4581– 4587.

[35] von Teichman, B., Engelbrecht, A., Zulu, G., Dungu, B., Pardini, A., and Bouloy, M. (2011). Safety and efficacy of Rift Valley fever Smithburn and Clone 13 vaccines in calves. Vaccine, 29(34): 5771–5777.

[36] Njenga, M. K., Njagi, L., Thumbi, S. M., Kahariri, S., Githinji, J., Omondi, E., Baden, A., Murithi, M., Paweska, J., Ithondeka, P. M., et al. (2015). Randomized controlled field trial to assess the immunogenicity and safety of Rift Valley Fever Clone 13 vaccine in livestock. PLoS Neglected Tropical Diseases, 9(3):e0003550.

[37] Miller, M. M., Bennett, K. E., Drolet, B. S., Lindsay, R., Mecham, J. O., Reeves, W. K., Weingartl, H. M., and Wilson, W. C. (2015). Evaluation of the efficacy, potential for vector transmission, and duration of immunity of MP-12, an attenuated Rift Valley fever virus vaccine candidate, in sheep. Clinical and Vaccine Immunology, 22(8):930–937.

[38] Duan, J., Li, S., and Xu, Y. (2023). Sequential Monte Carlo optimization and statistical inference. Wiley Interdisciplinary Reviews: Computational Statistics, 15(3):e1598.

[39] Galassi, M. et al. (2018). GNU Scientific Library Reference Manual. https://www.gnu.org/software/gsl/.

[40] R Core Team. (2023). R: A language and environment for statistical computing. R Foundation for Statistical Computing, Vienna, Austria. https://www.R-project.org/.

[41] Wickham, H., Averick, M., Bryan, J., Chang, W., McGowan, L. D., François, R., Grolemund, G., Hayes, A., Henry, L., Hester, J., et al. (2019). Welcome to the tidyverse. Journal of Open Source Software, 4(43):1686. doi: 10.21105/joss.01686.

[42] Tennant, W. S., Cardinale, E., Spencer, S. E., Tildesley, M. J., and Métras, R. (2024). Supporting data and code for the paper entitled Effectiveness and equity of vaccination strategies against Rift Valley fever in a heterogeneous landscape. GitHub repository, https://github.com/wtennant/rvf_vaccine.

[43] Gilbert, M., Nicolas, G., Cinardi, G., Van Boeckel, T. P., Vanwambeke, S. O., Wint, G., and Robinson, T. P. (2018). Global distribution data for cattle, buffaloes, horses, sheep, goats, pigs, chickens and ducks in 2010. Scientific data, 5(1):1–11.

[44] Janelle, J., Issoufi, A., Grimaldine, A., and Tillard, E. (2013). Référentiel technico-économique des élevages d’ovins et de caprins à Mayotte. Centre de coopération internationale en recherche agronomique pour le développement.

[45] Tillard, E., Moussa, T., Balberini, L., Aubriot, D., and Berre, D. (2013). Référentiel technico-économique des élevages de bovins à mayotte. Centre de coopération internationale en recherche agronomique pour le développement.

[46] Chopin, N. (2002). A sequential particle filter method for static models. Biometrika, 89(3):539–552.

