## Supplementary Information for "Effectiveness and equity of vaccination strategies against Rift Valley fever in a heterogeneous landscape"

### Supplementary Methods

#### Supplementary Methods 1: Mathematical model

Below, we describe the demographic and infection processes of the mathematical model as in Tennant et al. [1]. This model was extended to include a description of vaccination (see the main text).

##### Demographic process

A weekly proportion of livestock, $\delta_a$, move due to ageing from age group $a$ to age group $a+1$, and livestock die at a weekly age-dependent proportion denoted by $\mu_a$. The initial proportion of livestock in age group $a$ is denoted by $p_a$. Livestock in age group $a$ move from island $i$ to island $j$ with weekly proportion $m_{i,j,a}$, where $m_{i,i,a}$ denotes the weekly proportion of livestock of age group $a$ remaining on island $i$. Only livestock up to and including age group $A^{\text{move}}$ are moved between islands in the metapopulation. Each week, livestock are born into the first age group at a rate of $\nu_i$ and assumed to be susceptible to infection. The total livestock population on each island are assumed to be constant over time, therefore

$$\nu_i = \sum_{a=1}^{A} \left[ \mu_a N_{t,i,a} + (1-\mu_a) \sum_{j=1}^{n} (m_{i,j,a} N_{t,i,a} - m_{j,i,a} N_{t,j,a}) \right], \qquad (1)$$

where $N_{t,i,a}$ denotes the total livestock population at time $t$ on island $i$ in age group $a$.

##### Infection process

Susceptible livestock ($S$) are infected at a time- and island-dependent weekly proportion denoted by $\lambda_{t,i}$ such that

$$\lambda_{t,i} = 1 - \exp\left( \beta_{t,i} \frac{\sum_{a=1}^{A} I_{t,i,a}}{\sum_{a=1}^{A} N_{t,i,a}} \right), \qquad (2)$$

where $\beta_{t,i}$ denotes the weekly transmission rate of RVFV. As mosquitoes are not explicitly modelled, Normalised Difference Vegetation Index (NDVI) is used as a proxy to capture the effects of seasonal oscillations in mosquito demography on Rift Valley fever virus transmission rates. We assumed that the transmission rate scaled exponentially with island-specific increases NDVI, denoted by $\alpha$, where the (natural logarithm of the) minimum transmission rate per island is denoted by $\gamma_i$ [1]. Therefore,

$$\beta_{t,i} = \exp\left[ \alpha \left( \text{NDVI}_{t,i} - \min_{s,j} \text{NDVI}_{s,j} \right) + \gamma_i \right], \qquad (3)$$

where $\text{NDVI}_{t,i}$ denotes the Normalised Difference Vegetation Index at time $t$ on island $i$.

Upon infection, livestock move into the exposed compartment ($E$) where they remain for one week before transitioning to the infectious compartment ($I$). After one week in the infectious compartment, livestock transition into the recovered compartment ($R$) where they are assumed to have life-long immunity to reinfection. Infectious livestock of age group $a$ are introduced into island $i$ from outside the metapopulation system at a time-dependent rate $I^{\text{ext}}_{t,i,a}$. External introduction of livestock begins at time $t^{\text{ext}}_{(\text{start})}$ and lasts for $t^{\text{ext}}_{(\text{duration})}$ weeks. Infectious livestock are externally introduced in this way every $t^{\text{ext}}_{(\text{freq})}$ weeks. Only infectious livestock up to age $A^{\text{ext}}$ are imported into the metapopulation, and are assumed to be distributed in age according to the initial age distribution of livestock, $p_a$. Therefore,

$$I^{\text{ext}}_{t,i,a} = \begin{cases} \iota^{\text{ext}}_i \dfrac{p_a}{\sum_{b=1}^{A^{\text{ext}}} p_b}, & \text{for } a \in \left\{1, \ldots, A^{\text{ext}}\right\} \text{ and } t - t^{\text{ext}}_{(\text{start})} \leq t^{\text{ext}}_{(\text{duration})} \left( \bmod\, t^{\text{ext}}_{(\text{freq})} \right), \\ 0, & \text{otherwise,} \end{cases} \tag{4}$$

where $\iota^{\text{ext}}_i$ denotes the expected number of infectious livestock introduced into island $i$ each week.

#### Model equations

With the demographic and infection processes as described above, and the model structure and vaccination process described in the main text, the complete set of model equations is defined as follows.

Given the number of livestock in each compartment at time $t$ on island $i$ and rates of transition between compartments as defined above, the number of livestock in each compartment at time $t+1$ was defined as follows: for animals in the first age group ($a = 1$),

$$S^U_{t+1,i,1} = \nu_{t,i} + \sum_{j=1}^{n} m_{j,i,1} (1-\delta_1)(1-\mu_1)(1-\lambda_{t,j}) \left[ (1-\xi_{t,j,1}) S^U_{t,j,1} + \omega S^W_{t,j,1} \right] \tag{5}$$

$$E^U_{t+1,i,1} = \sum_{j=1}^{n} m_{j,i,1} (1-\delta_1)(1-\mu_1) \lambda_{t,j} \left[ (1-\xi_{t,j,1}) S^U_{t,j,1} + \omega S^W_{t,j,1} \right] \tag{6}$$

$$I^U_{t+1,i,1} = I^{\text{ext}}_{i,1} + \sum_{j=1}^{n} m_{j,i,1} (1-\delta_1)(1-\mu_1) \left[ (1-\xi_{t,j,1}) E^U_{t,j,1} + \omega E^W_{t,j,1} \right] \tag{7}$$

$$R^U_{t+1,i,1} = \sum_{j=1}^{n} m_{j,i,1} (1-\delta_1)(1-\mu_1) \left[ (1-\xi_{t,j,1}) \left( I^U_{t,j,1} + R^U_{t,j,1} \right) + \omega \left( I^W_{t,j,1} + R^W_{t,j,1} \right) \right] \tag{8}$$

$$S^{V_1}_{t+1,i,1} = \sum_{j=1}^{n} m_{j,i,1} (1-\delta_1)(1-\mu_1)(1-\lambda_{t,j}) \xi_{t,j,1} S^U_{t,j,1} \tag{9}$$

$$E^{V_1}_{t+1,i,1} = \sum_{j=1}^{n} m_{j,i,1} (1-\delta_1)(1-\mu_1) \lambda_{t,j} \xi_{t,j,1} S^U_{t,j,1} \tag{10}$$

$$I^{V_1}_{t+1,i,1} = \sum_{j=1}^{n} m_{j,i,1} (1-\delta_1)(1-\mu_1) \xi_{t,j,1} E^U_{t,j,1} \tag{11}$$

$$R^{V_1}_{t+1,i,1} = \sum_{j=1}^{n} m_{j,i,1} (1-\delta_1)(1-\mu_1) \xi_{t,j,1} \left( I^U_{t,j,1} + R^U_{t,j,1} \right) \tag{12}$$

$$S^{V_2}_{t+1,i,1} = \sum_{j=1}^{n} m_{j,i,1} \left(1-\delta_1\right)\left(1-\mu_1\right)\left(1-\lambda_{t,j}\right) S^{V_1}_{t,j,1} \tag{13}$$

$$E^{V_2}_{t+1,i,1} = \sum_{j=1}^{n} m_{j,i,1} \left(1-\delta_1\right)\left(1-\mu_1\right)\lambda_{t,j} S^{V_1}_{t,j,1} \tag{14}$$

$$I^{V_2}_{t+1,i,1} = \sum_{j=1}^{n} m_{j,i,1} \left(1-\delta_1\right)\left(1-\mu_1\right) E^{V_1}_{t,j,1} \tag{15}$$

$$R^{V_2}_{t+1,i,1} = \sum_{j=1}^{n} m_{j,i,1} \left(1-\delta_1\right)\left(1-\mu_1\right)\left(I^{V_1}_{t,j,1} + R^{V_1}_{t,j,1}\right) \tag{16}$$

$$S^{W}_{t+1,i,1} = \sum_{j=1}^{n} m_{j,i,1} \left(1-\delta_1\right)\left(1-\mu_1\right)\left[\left(1-\lambda_{t,j}\right) S^{V_2}_{t,j,1} + \left(1-p^{\text{eff}}\lambda_{t,j}\right)\left(1-\omega\right) S^{W}_{t,j,1}\right] \tag{17}$$

$$E^{W}_{t+1,i,1} = \sum_{j=1}^{n} m_{j,i,1} \left(1-\delta_1\right)\left(1-\mu_1\right)\lambda_{t,j}\left[S^{V_2}_{t,j,1} + p^{\text{eff}}\left(1-\omega\right) S^{W}_{t,j,1}\right] \tag{18}$$

$$I^{W}_{t+1,i,1} = \sum_{j=1}^{n} m_{j,i,1} \left(1-\delta_1\right)\left(1-\mu_1\right)\left(E^{V_2}_{t,j,1} + \left(1-\omega\right) E^{W}_{t,j,1}\right) \tag{19}$$

$$R^{W}_{t+1,i,1} = \sum_{j=1}^{n} m_{j,i,1} \left(1-\delta_1\right)\left(1-\mu_1\right)\left[I^{V_2}_{t,j,1} + R^{V_2}_{t,j,1} + \left(1-\omega\right)\left(I^{W}_{t,j,1} + R^{W}_{t,j,1}\right)\right], \tag{20}$$

and for livestock in the remaining age groups $a \in \{2, \ldots, A\}$,

$$\begin{aligned} S^{U}_{t+1,i,a} = &\sum_{j=1}^{n} m_{j,i,a} \left(1-\delta_a\right)\left(1-\mu_a\right)\left(1-\lambda_{t,j}\right)\left[\left(1-\xi_{t,j,a}\right) S^{U}_{t,j,a} + \omega S^{W}_{t,j,a}\right] \\ &+ \sum_{j=1}^{n} m_{j,i,a-1}\, \delta_{a-1}\left(1-\mu_{a-1}\right)\left(1-\lambda_{t,j}\right)\left[\left(1-\xi_{t,j,a-1}\right) S^{U}_{t,j,a-1} + \omega S^{W}_{t,j,a-1}\right], \end{aligned} \tag{21}$$

$$\begin{aligned} E^{U}_{t+1,i,a} = &\sum_{j=1}^{n} m_{j,i,a} \left(1-\delta_a\right)\left(1-\mu_a\right)\lambda_{t,j}\left[\left(1-\xi_{t,j,a}\right) S^{U}_{t,j,a} + \omega S^{W}_{t,j,a}\right] \\ &+ \sum_{j=1}^{n} m_{j,i,a-1}\, \delta_{a-1}\left(1-\mu_{a-1}\right)\lambda_{t,j}\left[\left(1-\xi_{t,j,a-1}\right) S^{U}_{t,j,a-1} + \omega S^{W}_{t,j,a-1}\right], \end{aligned} \tag{22}$$

$$\begin{aligned} I^{U}_{t+1,i,a} = &\, I^{\text{ext}}_{t,i,a} + \sum_{j=1}^{n} m_{j,i,a} \left(1-\delta_a\right)\left(1-\mu_a\right)\left[\left(1-\xi_{t,j,a}\right) E^{U}_{t,j,a} + \omega E^{W}_{t,j,a}\right] \\ &+ \sum_{j=1}^{n} m_{j,i,a-1}\, \delta_{a-1}\left(1-\mu_{a-1}\right)\left[\left(1-\xi_{t,j,a-1}\right) E^{U}_{t,j,a-1} + \omega E^{W}_{t,j,a-1}\right], \end{aligned} \tag{23}$$

$$\begin{aligned} R^{U}_{t+1,i,a} = &\sum_{j=1}^{n} m_{j,i,a} \left(1-\delta_a\right)\left(1-\mu_a\right)\left[\left(1-\xi_{t,j,a}\right)\left(I^{U}_{t,j,a} + R^{U}_{t,j,a}\right) + \omega\left(I^{W}_{t,j,a} + R^{W}_{t,j,a}\right)\right] \\ &+ \sum_{j=1}^{n} m_{j,i,a-1}\, \delta_{a-1}\left(1-\mu_{a-1}\right)\left[\left(1-\xi_{t,j,a-1}\right)\left(I^{U}_{t,j,a-1} + R^{U}_{t,j,a-1}\right)\right. \\ &\qquad\qquad\qquad\qquad \left. + \,\omega\left(I^{W}_{t,j,a-1} + R^{W}_{t,j,a-1}\right)\right], \end{aligned} \tag{24}$$

$$\begin{aligned} S^{V_1}_{t+1,i,a} = &\sum_{j=1}^{n} m_{j,i,a} \left(1-\delta_a\right)\left(1-\mu_a\right)\left(1-\lambda_{t,j}\right)\xi_{t,j,a} S^{U}_{t,j,a} \\ &+ \sum_{j=1}^{n} m_{j,i,a-1}\, \delta_{a-1}\left(1-\mu_{a-1}\right)\left(1-\lambda_{t,j}\right)\xi_{t,j,a-1} S^{U}_{t,j,a-1}, \end{aligned} \tag{25}$$

$$E^{V_1}_{t+1,i,a} = \sum_{j=1}^{n} m_{j,i,a}\left(1-\delta_a\right)\left(1-\mu_a\right)\lambda_{t,j}\xi_{t,j,a}S^{U}_{t,j,a} + \sum_{j=1}^{n} m_{j,i,a-1}\,\delta_{a-1}\left(1-\mu_{a-1}\right)\lambda_{t,j}\xi_{t,j,a-1}S^{U}_{t,j,a-1}, \tag{26}$$

$$I^{V_1}_{t+1,i,a} = \sum_{j=1}^{n} m_{j,i,a}\left(1-\delta_a\right)\left(1-\mu_a\right)\xi_{t,j,a}E^{U}_{t,j,a} + \sum_{j=1}^{n} m_{j,i,a-1}\,\delta_{a-1}\left(1-\mu_{a-1}\right)\xi_{t,j,a-1}E^{U}_{t,j,a-1}, \tag{27}$$

$$R^{V_1}_{t+1,i,a} = \sum_{j=1}^{n} m_{j,i,a}\left(1-\delta_a\right)\left(1-\mu_a\right)\xi_{t,j,a}\left(I^{U}_{t,j,a}+R^{U}_{t,j,a}\right) + \sum_{j=1}^{n} m_{j,i,a-1}\,\delta_{a-1}\left(1-\mu_{a-1}\right)\xi_{t,j,a-1}\left(I^{U}_{t,j,a-1}+R^{U}_{t,j,a-1}\right), \tag{28}$$

$$S^{V_2}_{t+1,i,a} = \sum_{j=1}^{n} m_{j,i,a}\left(1-\delta_a\right)\left(1-\mu_a\right)\left(1-\lambda_{t,j}\right)S^{V_1}_{t,j,a} + \sum_{j=1}^{n} m_{j,i,a-1}\,\delta_{a-1}\left(1-\mu_{a-1}\right)\left(1-\lambda_{t,j}\right)S^{V_1}_{t,j,a-1}, \tag{29}$$

$$E^{V_2}_{t+1,i,a} = \sum_{j=1}^{n} m_{j,i,a}\left(1-\delta_a\right)\left(1-\mu_a\right)\lambda_{t,j}S^{V_1}_{t,j,a} + \sum_{j=1}^{n} m_{j,i,a-1}\,\delta_{a-1}\left(1-\mu_{a-1}\right)\lambda_{t,j}S^{V_1}_{t,j,a-1}, \tag{30}$$

$$I^{V_2}_{t+1,i,a} = \sum_{j=1}^{n} m_{j,i,a}\left(1-\delta_a\right)\left(1-\mu_a\right)E^{V_1}_{t,j,a} + \sum_{j=1}^{n} m_{j,i,a-1}\,\delta_{a-1}\left(1-\mu_{a-1}\right)E^{V_1}_{t,j,a-1}, \tag{31}$$

$$R^{V_2}_{t+1,i,a} = \sum_{j=1}^{n} m_{j,i,a}\left(1-\delta_a\right)\left(1-\mu_a\right)\left(I^{V_1}_{t,j,a}+R^{V_1}_{t,j,a}\right) + \sum_{j=1}^{n} m_{j,i,a-1}\,\delta_{a-1}\left(1-\mu_{a-1}\right)\left(I^{V_1}_{t,j,a-1}+R^{V_1}_{t,j,a-1}\right), \tag{32}$$

$$S^{W}_{t+1,i,a} = \sum_{j=1}^{n} m_{j,i,a}\left(1-\delta_a\right)\left(1-\mu_a\right)\left[\left(1-\lambda_{t,j}\right)S^{V_2}_{t,j,a} + \left(1-p_{\text{eff}}\lambda_{t,j}\right)\left(1-\omega\right)S^{W}_{t,j,a}\right] + \sum_{j=1}^{n} m_{j,i,a-1}\,\delta_{a-1}\left(1-\mu_{a-1}\right)\left[\left(1-\lambda_{t,j}\right)S^{V_2}_{t,j,a-1} + \left(1-p_{\text{eff}}\lambda_{t,j}\right)\left(1-\omega\right)S^{W}_{t,j,a-1}\right], \tag{33}$$

$$E^{W}_{t+1,i,a} = \sum_{j=1}^{n} m_{j,i,a}\left(1-\delta_a\right)\left(1-\mu_a\right)\lambda_{t,j}\left[S^{V_2}_{t,j,a} + p_{\text{eff}}\left(1-\omega\right)S^{W}_{t,j,a}\right] + \sum_{j=1}^{n} m_{j,i,a-1}\,\delta_{a-1}\left(1-\mu_{a-1}\right)\lambda_{t,j}\left[S^{V_2}_{t,j,a-1} + p_{\text{eff}}\left(1-\omega\right)S^{W}_{t,j,a-1}\right], \tag{34}$$

$$I^{W}_{t+1,i,a} = \sum_{j=1}^{n} m_{j,i,a}\left(1-\delta_a\right)\left(1-\mu_a\right)\left(E^{V_2}_{t,j,a} + \left(1-\omega\right)E^{W}_{t,j,a}\right) + \sum_{j=1}^{n} m_{j,i,a-1}\,\delta_{a-1}\left(1-\mu_{a-1}\right)\left(E^{V_2}_{t,j,a-1} + \left(1-\omega\right)E^{W}_{t,j,a-1}\right), \tag{35}$$

$$R^{W}_{t+1,i,a} = \sum_{j=1}^{n} m_{j,i,a} (1-\delta_a)(1-\mu_a) \left[ I^{V_2}_{t,j,a} + R^{V_2}_{t,j,a} + (1-\omega)\left(I^{W}_{t,j,a} + R^{W}_{t,j,a}\right)\right]$$
$$+ \sum_{j=1}^{n} m_{j,i,a-1}\, \delta_{a-1} (1-\mu_{a-1}) \left[ I^{V_2}_{t,j,a-1} + R^{V_2}_{t,j,a-1} + (1-\omega)\left(I^{W}_{t,j,a-1} + R^{W}_{t,j,a-1}\right)\right]. \quad (36)$$

At time $t = 0$, the livestock population on each island $i$ was assumed to be entirely unprotected from any vaccine, and so for $a \in \{1, \ldots, A\}$ and $i \in \{1, \ldots, N\}$,

$$S^{U}_{0,i,a} = p_a \left[(1-\epsilon_i) N_i - E_{0,i} - I_{0,i}\right], \quad (37)$$

$$E^{U}_{0,i,a} = p_a E_{0,i}, \quad (38)$$

$$I^{U}_{0,i,a} = p_a I_{0,i}, \quad (39)$$

$$R^{U}_{0,i,a} = \epsilon_i p_a N_i, \quad (40)$$

$$\mathcal{X}^{\mathcal{V}}_{0,i,a} = 0, \qquad \text{for } \mathcal{X} \in \{S, E, I, R\} \text{ and } \mathcal{V} \in \{V_1, V_2, W\}, \quad (41)$$

where $N_i$ denoted the total livestock population on island $i$, $E_{0,i}$ denoted the number of exposed animals at time $t = 0$, $I_{0,i}$ denoted the number of exposed animals at time $t = 0$ and $\epsilon_i$ was the initial proportion of the population on each island $i$ that have recovered from the virus.

#### Demographic and infection process parameterisation

The demographic and infection processes were parameterised as in the previous model by Tennant et al. [1]. Below, we describe the parameterisation of these processes in detail.

Using the Comoros archipelago as a case study to evaluate the impact of different vaccine strategies against RVFV, the number of islands in the metapopulation was set to four ($n = 4$). The livestock population was subdivided into 10 age groups ($A = 10$), with each age group representing livestock of 0–1 years old, 1–2 years old, ..., and greater than 9 years old. Only livestock in the first two age groups could be externally introduced ($A^{\text{ext}} = 2$) or moved between islands in the metapopulation ($A^{\text{move}} = 2$). The initial time ($t = 0$) corresponded to July 2004.

The initial number of exposed and infectious individuals was assumed to be small with $E_{0,i} = I_{0,i} = 5$. The population size of each island in the Comoros archipelago, $N_i$, was calculated using the Gridded Livestock Map of the World [2], with initial age distribution, $p_a$, set to the age profiles calculated by Janelle et al. [3] and Tillard et al. [4]. Each time step in the model represented 1.08 weeks (an epidemiological week), and thus the weekly ageing probability $\delta_a$ for the first nine age groups was set to 1/48 with $\delta_{10} = 0$. The mortality rates of age groups 1–9 and 10 were set to $8.8 \times 10^{-3}$ and $6.2 \times 10^{-3}$ respectively [3, 4]. Infectious imports were introduced into the system every 10 years ($t^{\text{ext}}_{(\text{freq})} = 480$).

The model in the absence of vaccination ($\psi = 0$) was fitted in a Bayesian framework to a series of cross-sectional and longitudinal seroprevalence surveys conducted on each of the four islands in the Comoros archipelago from July 2004 until June 2015 ($t = 527$). Fitting the model to these data informed the remaining demographic and infection process parameters of the model: the probability of moving between islands per week, $m_{i,j}$, the island-specific disease transmission rate parameters, $\alpha$ and $\gamma_i$, external importation of livestock, $\iota_i^{\text{ext}}$, $t_{(\text{start})}^{\text{ext}}$ and $t_{(\text{duration})}^{\text{ext}}$, and initial proportion of the population immune to RVFV on each island, $\epsilon_i$. Supplementary Table 10 shows a list of posterior parameter estimates and Supplementary Figure 20 shows the posterior distribution of each parameter inferred through model fitting. Refer to Tennant et al. [1] for the full details and discussion of the model fitting procedure and results.

#### Supplementary Methods 2: Optimisation algorithm

In order to determine the optimal allocation of vaccines, a sequential Monte Carlo (SMC) scheme was employed. This algorithm consisted of three main parts:

1. a sampling part, in which particles with a higher objective function evaluation were favoured more highly than others,
2. a density-tempering part, which allowed particles to move from the initially proposed distribution of particles towards an empirical distribution of the objective function, and
3. a rejuvenation part, which meant that sampled particles would be jittered using a Metropolis-Hastings kernel with a (truncated) multivariate Gaussian proposal distribution.

Below we describe each of these three components. The complete optimisation algorithm and associated notation can be found in Supplementary Algorithm 1.

##### Sampling particles

The algorithm aimed to obtain a set of vaccine distributions that were representative of the objective function $f(\rho)$. However, the objective function may not be positive for all $\rho \in P$ and may not directly define a probability density function. Instead, we obtained samples from the following monotonic transformation of the objective function:

$$h(\rho) \propto \exp\left[m f(\rho)\right], \tag{42}$$

where $m \geq 1$ defines the peakiness of the transformed objective function.

**Input :** $J \geq 1$, // Number of SMC particles
$K \geq 1$, // Number of effectiveness samples
$I(\rho)$, // Initial distribution of SMC particles
$g(\rho|\theta)$, // Effectiveness function
$\pi(\theta)$, // Joint distribution of model parameters
$m \geq 1$, // Scalar for objective function
$d_1 \in [0,1]$, // First tempering weight of objective function
$q(.|\rho)$, // Proposal distribution
$c_{\max} > 0$. // Maximum cumulative acceptance rate
**Output:** $\rho^*$ such that $f(\rho^*) = \max_{\rho \in P} f(\rho) = \max_{\rho \in P} \int_\theta g(\rho|\theta)\, h(\theta)\, \mathrm{d}\theta$.

// Initialisation of particles and initial evaluation of objective function.
**for** $j = 1$ **to** $J$ **do**
    $w_j^{(1)} \leftarrow 1/J$;
    $\rho_j^{(1)} \sim I(\rho)$;
    **for** $k = 1$ **to** $K$ **do**
        $\theta_k \sim \pi(\theta)$;
        $g_{jk} \leftarrow g\left(\rho_j^{(1)}|\theta_k\right)$;
    **end**
    $\hat{f}_j \leftarrow \frac{1}{K}\sum_{k=1}^{K} g_{jk}$;
    $h_j^{(1)} \leftarrow \exp\left(m\hat{f}_j\right)^{d_1} I(\rho_j)^{(1-d_1)}$;
**end**
$s \leftarrow 1$; // Progress through tempering weights until the initial distribution vanishes.
**while** $d_s < 1$ **do**
    **Find** the largest $d_{s+1} \in [d_s, 1]$ such that $\left(\sum_{j=1}^{J} w_j^{(s+1)}\right)^2 / \sum_{j=1}^{J}\left(w_j^{(s+1)}\right)^2 \geq J/2$ with
        $w_j^{(s+1)} = w_j^{(s)} h_j^{(s+1)} / h_j^{(s)}$ and $h_j^{(s)} = \exp\left(m\hat{f}_j\right)^{d_s} I(\rho_j)^{(1-d_s)}$;
    **Sample** $\rho_j^{(s+1)}$ from $\rho_j^{(s)}$ with weights $w_j^{(s+1)}$;
    $c \leftarrow 0$; // Jitter particles using a Metropolis-Hastings algorithm.
    **while** $c < c_{\max}$ **do**
        **for** $j = 1$ **to** $J$ **do**
            $\rho_j' \sim q(.|\rho_j)$;
            **for** $k = 1$ **to** $K$ **do**
                $\theta_k \sim \pi(\theta)$;
                $g_{jk}' \leftarrow g\left(\rho_j'|\theta_k\right)$;
            **end**
            $\hat{f}_j{}' \leftarrow \frac{1}{K}\sum_{k=1}^{K} g_{jk}$;
            $h_j' \leftarrow \exp\left(m\hat{f}_j{}'\right)^{d_{s+1}} I\left(\rho_j'\right)^{(1-d_{s+1})}$;
            $\alpha \leftarrow \left[h_j' q\left(\rho_j'|\rho_j\right)\right] / \left[h_j^{(s+1)} q\left(\rho_j|\rho_j'\right)\right]$;
            $v \sim U(0,1)$;
            **if** $\alpha < \min\{1, v\}$ **then**
                $\rho_j^{(s+1)} \leftarrow \rho_j'$;
                $h_j^{(s+1)} \leftarrow h_j'$;
                $c \leftarrow c + 1/J$;
            **end**
        **end**
    **end**
    $s \leftarrow s + 1$;
**end**
$j^* \leftarrow \arg\max_{1 \leq j \leq J} h_j^{(s)}$; // Extract the particle with the largest evaluation.
$\rho^* \leftarrow \rho_{j^*}$;

**Supplementary Algorithm 1: Sequential Monte Carlo (SMC) optimisation algorithm.** Pseudo-code of the algorithm used to determine the optimal distribution of vaccines across all islands in the Comoros archipelago. Here, particles corresponded to a unique distribution of vaccines, and the objective function corresponded to the mean effectiveness of a given strategy. The algorithm included tempering from the initial distribution of particles towards the distribution of particles governed by the objective function.

The transformed function defined a probability density function (up to an unknown normalising constant) from which particles could be sampled. As the transformation was (positive) monotonic, the optimal vaccine distribution for $h(\rho)$ was the same as the objective function $f(\rho)$:

$$\rho_j \sim h(.), \qquad j \in \{1, ..., J\}, \tag{43}$$

$$\hat{\rho}^* = \arg\max_{j \in \{1,...,J\}} h(\rho_j) = \arg\max_{j \in \{1,...,J\}} f(\rho_k), \tag{44}$$

where $\rho_j$ denotes a single sample from the transformed objective function, $J$ is the total number of particles to be sampled, and $\hat{\rho}^*$ is an estimate of the optimal vaccine distribution.

In order to obtain samples from the transformed objective function, we first sampled particles from some known sampling distribution, denoted $I(\rho)$, and used importance sampling to filter the particles towards $h(\rho)$. To filter the initial set of particles in a single step may lead to a poorly representative sample of $h(\rho)$, so we introduced an intermediate set of density-tempered distributions, denoted $h_{d_s}(\rho)$, which allowed a more smooth transition from $I(\rho)$ to $h(\rho)$ across multiple steps.

##### Intermediate density-tempered distributions

At each step $s$ in the filtering process, we defined the intermediate density-tempered distributions as:

$$h_{d_s}(\rho) = h(\rho)^{d_s} I(\rho)^{(1-d_s)}, \tag{45}$$

where $d_s \in [0, 1]$ is the density-tempering parameter at step $s$, which monotonically increase with $s$. By setting $d_1 = 0$, the first intermediate density-tempered distribution, $h_{d_1}(\rho)$, is equivalent to the initial distribution $I(\rho)$.

Given a set of particles representing $h_{d_s}(\rho)$, the particles were filtered to the next intermediate distribution $h_{d_{s+1}}(\rho)$ by first calculating the following importance weights:

$$w_j^{(s+1)} = w_j^{(s)} \frac{h_{d_{s+1}}\left(\rho_j^{(s)}\right)}{h_{d_s}\left(\rho_j^{(s)}\right)}, \tag{46}$$

$$w_j^{(1)} := \frac{1}{J}, \tag{47}$$

where $w_j^{(s)}$ denotes the importance weight at step $s$, and $\rho_j^{(s)}$ is a sample from the intermediate density-tempered distribution $h_{d_s}(\rho)$. The importance weights were then used to sample the next set of particles $\rho_j^{(s+1)}$. Using this approach, the normalising constant of $h(\rho)$ did not need to be known.

To ensure a smooth transition between intermediate density-tempered distributions, the sequence of density-tempering parameters was chosen to be the largest $d_{s+1} \in [d_s, 1]$ such that the effective sample size (ESS) from one intermediate distribution to the next remained above $J/2$, where

$$\text{ESS} = \frac{\left(\sum_{j=1}^{J} w_j^{(s+1)}\right)^2}{\sum_{j=1}^{J} \left(w_j^{(s+1)}\right)^2}. \tag{48}$$

##### Particle rejuvenation

Repeatedly filtering particles from one intermediate distribution to the next alone would result in a lack of diversity among particles, leading to a poor representation of the final distribution $h(\rho)$. To ameliorate this, we used a Metropolis-Hastings step to jitter and rejuvenate the set of particles.

After each resampling step in the particle filter, we proposed a new particle $\rho_j'$ from a proposal distribution $q(.|\rho_j)$ for all $j \in \{1, ..., J\}$. The proposed particle $\rho_j'$ was then evaluated using the current intermediate distribution, and replaced the previous particle $\rho_j$ with probability

$$\alpha = \min\left\{1, \frac{h\left(\rho_j'\right)^{d_s} I\left(\rho_j'\right)^{1-d_s}}{h\left(\rho_j\right)^{d_s} I\left(\rho_j\right)^{1-d_s}} \frac{q\left(\rho_j|\rho_j'\right)}{q\left(\rho_j'|\rho_j\right)}\right\}, \tag{49}$$

where $h\left(\rho_j'\right)^{d_s} I\left(\rho_j'\right)^{1-d_s}$ denotes the intermediate distribution at step $s$. This procedure was repeated until a cumulative acceptance rate, $c_{\max}$, was reached.

In this work the proposal distribution $q(.|\rho_j)$ was selected to be a mixture distribution with equal weights between: (i) the initialisation distribution $I$, (ii) a multivariate Gaussian with mean $\rho_j$ and covariance equal to the identity matrix scaled by $\frac{0.1^2}{n}$, representing a small random walk around the current particle $\rho_j$, and (iii) a multivariate Gaussian with mean and covariance matrix calculated directly from the entire set of particles $\rho$. This proposal distribution allowed the parameter space to be continuously explored, whilst considering information from the current set of particles [5].

##### Optimal particle

Particles were filtered sequentially through each intermediate distribution and rejuvenated in turn up to step $S$ with $d_S = 1$. This yielded a set of particles representing the transformed objective function $h(\rho)$. An estimate for the optimal vaccine distribution was determined as the vaccine distribution $\rho$ which maximised $h(\rho)$ as in Supplementary Equation 45.

#### Supplementary References

[1] Tennant, W. S., Cardinale, E., Cêtre-Sossah, C., Moutroifi, Y., Le Godais, G., Colombi, D., Spencer, S. E., Tildesley, M. J., Keeling, M. J., Charafouddine, O., et al. (2021). Modelling the persistence

and control of Rift Valley fever virus in a spatially heterogeneous landscape. *Nature communications*, 12(1):5593.

[2] Gilbert, M., Nicolas, G., Cinardi, G., Van Boeckel, T. P., Vanwambeke, S. O., Wint, G., and Robinson, T. P. (2018). Global distribution data for cattle, buffaloes, horses, sheep, goats, pigs, chickens and ducks in 2010. *Scientific data*, 5(1):1–11.

[3] Janelle, J., Issoufi, A., Grimaldine, A., and Tillard, E. (2013). Référentiel technico-économique des élevages d'ovins et de caprins à Mayotte. *Centre de coopération internationale en recherche agronomique pour le développement.*

[4] Tillard, E., Moussa, T., Balberini, L., Aubriot, D., and Berre, D. (2013). Référentiel technico-économique des élevages de bovins à mayotte. *Centre de coopération internationale en recherche agronomique pour le développement.*

[5] Chopin, N. (2002). A sequential particle filter method for static models. *Biometrika*, 89(3):539–552.

#### Supplementary Tables

**Supplementary Table 1: Vaccine allocation to islands in the Comoros archipelago under a range of vaccine strategies**. Assuming animals were not tagged (U) or tagged (T) and given a total percentage of livestock to vaccinate annually across the archipelago, vaccines were either allocated to each of the four islands proportional to population size (P), optimised in terms of infections averted across the archipelago (1) or optimised in terms of infections averted on the worst-performing island (2). The table shows the median and 95% credible interval for the percentage of vaccines assigned to each island in the archipelago for each vaccine strategy and six different vaccination rates. The allocation of vaccines for the proportional to island population size case was independent of the percentage of livestock vaccinated annually across the archipelago. For the optimised strategies, each median and 95% credible interval was calculated from 500 executions of the optimisation algorithm.

| Strategy identifier | Percentage of livestock vaccinated annually across the archipelago | Island | Median percentage of vaccines assigned to the island [95% credible interval] |
|---|---|---|---|
| UP | Any% | Grande Comore | 60.65% [60.65%, 60.65%] |
| UP | Any% | Mohéli | 8.62% [8.62%, 8.62%] |
| UP | Any% | Anjouan | 25.31% [25.31%, 25.31%] |
| UP | Any% | Mayotte | 5.42% [5.42%, 5.42%] |
| TP | Any% | Grande Comore | 60.65% [60.65%, 60.65%] |
| TP | Any% | Mohéli | 8.62% [8.62%, 8.62%] |
| TP | Any% | Anjouan | 25.31% [25.31%, 25.31%] |
| TP | Any% | Mayotte | 5.42% [5.42%, 5.42%] |
| U1 | 5% | Grande Comore | 6.40% [0.24%, 20.90%] |
| U1 | 5% | Mohéli | 15.99% [2.70%, 33.91%] |
| U1 | 5% | Anjouan | 71.70% [55.43%, 87.16%] |
| U1 | 5% | Mayotte | 4.00% [0.25%, 11.09%] |
| U2 | 5% | Grande Comore | 59.78% [47.10%, 73.07%] |
| U2 | 5% | Mohéli | 18.66% [10.84%, 33.33%] |
| U2 | 5% | Anjouan | 14.40% [6.43%, 27.31%] |
| U2 | 5% | Mayotte | 4.39% [1.07%, 14.86%] |
| T1 | 5% | Grande Comore | 4.08% [0.15%, 17.37%] |
| T1 | 5% | Mohéli | 28.25% [8.20%, 51.86%] |
| T1 | 5% | Anjouan | 62.76% [38.17%, 81.34%] |
| T1 | 5% | Mayotte | 3.07% [0.13%, 9.93%] |
| T2 | 5% | Grande Comore | 62.63% [49.66%, 72.66%] |
| T2 | 5% | Mohéli | 18.09% [11.42%, 30.27%] |
| T2 | 5% | Anjouan | 14.11% [7.06%, 24.00%] |
| T2 | 5% | Mayotte | 3.73% [1.19%, 12.10%] |
| U1 | 10% | Grande Comore | 38.68% [21.69%, 54.20%] |
| U1 | 10% | Mohéli | 27.60% [13.42%, 35.89%] |
| U1 | 10% | Anjouan | 34.21% [18.09%, 44.37%] |
| U1 | 10% | Mayotte | 2.19% [0.14%, 5.76%] |
| U2 | 10% | Grande Comore | 67.53% [55.77%, 78.88%] |

**Supplementary Table 1 (continued): Vaccine allocation to islands in the Comoros archipelago under a range of vaccine strategies.** Refer to the initial table caption for the full description.

| Strategy identifier | Percentage of livestock vaccinated annually across the archipelago | Island | Median percentage of vaccines assigned to the island [95% credible interval] |
|---|---|---|---|
| U2 | 10% | Mohéli | 14.84% [8.53%, 27.21%] |
| U2 | 10% | Anjouan | 11.37% [5.64%, 22.04%] |
| U2 | 10% | Mayotte | 4.04% [0.91%, 13.61%] |
| T1 | 10% | Grande Comore | 25.47% [13.50%, 43.42%] |
| T1 | 10% | Mohéli | 34.28% [23.96%, 40.06%] |
| T1 | 10% | Anjouan | 37.51% [22.26%, 45.50%] |
| T1 | 10% | Mayotte | 2.25% [0.13%, 6.57%] |
| T2 | 10% | Grande Comore | 69.26% [60.67%, 77.69%] |
| T2 | 10% | Mohéli | 14.14% [9.19%, 22.58%] |
| T2 | 10% | Anjouan | 11.88% [7.24%, 19.56%] |
| T2 | 10% | Mayotte | 3.10% [0.88%, 9.28%] |
| U1 | 15% | Grande Comore | 57.55% [44.41%, 69.50%] |
| U1 | 15% | Mohéli | 20.73% [11.01%, 29.09%] |
| U1 | 15% | Anjouan | 20.16% [10.58%, 31.33%] |
| U1 | 15% | Mayotte | 1.85% [0.09%, 5.12%] |
| U2 | 15% | Grande Comore | 68.32% [59.39%, 78.12%] |
| U2 | 15% | Mohéli | 14.67% [8.97%, 24.80%] |
| U2 | 15% | Anjouan | 10.94% [6.34%, 20.17%] |
| U2 | 15% | Mayotte | 4.13% [0.76%, 12.79%] |
| T1 | 15% | Grande Comore | 61.07% [48.76%, 81.11%] |
| T1 | 15% | Mohéli | 18.56% [11.65%, 24.71%] |
| T1 | 15% | Anjouan | 18.63% [1.13%, 27.80%] |
| T1 | 15% | Mayotte | 1.53% [0.08%, 4.19%] |
| T2 | 15% | Grande Comore | 70.77% [64.17%, 78.01%] |
| T2 | 15% | Mohéli | 12.69% [9.12%, 19.84%] |
| T2 | 15% | Anjouan | 11.56% [8.30%, 17.45%] |
| T2 | 15% | Mayotte | 3.41% [0.85%, 10.55%] |
| U1 | 20% | Grande Comore | 62.14% [52.49%, 74.97%] |
| U1 | 20% | Mohéli | 16.50% [8.77%, 24.13%] |
| U1 | 20% | Anjouan | 19.73% [9.56%, 26.09%] |
| U1 | 20% | Mayotte | 1.56% [0.10%, 4.47%] |
| U2 | 20% | Grande Comore | 69.42% [59.95%, 78.43%] |
| U2 | 20% | Mohéli | 13.15% [8.52%, 22.52%] |
| U2 | 20% | Anjouan | 10.34% [6.70%, 20.40%] |

**Supplementary Table 1 (continued): Vaccine allocation to islands in the Comoros archipelago under a range of vaccine strategies.** Refer to the initial table caption for the full description.

| Strategy identifier | Percentage of livestock vaccinated annually across the archipelago | Island | Median percentage of vaccines assigned to the island [95% credible interval] |
|---|---|---|---|
| U2 | 20% | Mayotte | 4.38% [0.63%, 14.52%] |
| T1 | 20% | Grande Comore | 79.06% [64.80%, 93.03%] |
| T1 | 20% | Mohéli | 13.43% [1.84%, 18.40%] |
| T1 | 20% | Anjouan | 5.76% [0.48%, 18.24%] |
| T1 | 20% | Mayotte | 1.21% [0.04%, 3.87%] |
| T2 | 20% | Grande Comore | 71.63% [66.40%, 77.76%] |
| T2 | 20% | Mohéli | 11.41% [8.82%, 18.20%] |
| T2 | 20% | Anjouan | 11.89% [8.69%, 18.66%] |
| T2 | 20% | Mayotte | 3.10% [0.74%, 9.92%] |
| U1 | 25% | Grande Comore | 68.93% [58.79%, 78.63%] |
| U1 | 25% | Mohéli | 13.85% [6.36%, 21.85%] |
| U1 | 25% | Anjouan | 15.57% [9.15%, 21.35%] |
| U1 | 25% | Mayotte | 1.62% [0.15%, 4.78%] |
| U2 | 25% | Grande Comore | 70.26% [61.59%, 79.36%] |
| U2 | 25% | Mohéli | 11.70% [8.19%, 22.99%] |
| U2 | 25% | Anjouan | 11.27% [7.20%, 22.77%] |
| U2 | 25% | Mayotte | 3.99% [0.54%, 12.34%] |
| T1 | 25% | Grande Comore | 82.42% [70.77%, 93.71%] |
| T1 | 25% | Mohéli | 11.21% [1.29%, 15.58%] |
| T1 | 25% | Anjouan | 5.19% [0.21%, 14.83%] |
| T1 | 25% | Mayotte | 1.29% [0.10%, 4.22%] |
| T2 | 25% | Grande Comore | 71.46% [65.50%, 77.10%] |
| T2 | 25% | Mohéli | 12.13% [9.18%, 18.76%] |
| T2 | 25% | Anjouan | 11.73% [8.93%, 18.01%] |
| T2 | 25% | Mayotte | 3.02% [0.92%, 9.79%] |
| U1 | 30% | Grande Comore | 71.86% [62.60%, 82.42%] |
| U1 | 30% | Mohéli | 13.75% [4.51%, 20.57%] |
| U1 | 30% | Anjouan | 13.02% [7.30%, 18.28%] |
| U1 | 30% | Mayotte | 1.52% [0.09%, 5.24%] |
| U2 | 30% | Grande Comore | 70.86% [62.15%, 78.70%] |
| U2 | 30% | Mohéli | 12.41% [8.47%, 21.45%] |
| U2 | 30% | Anjouan | 10.33% [7.00%, 20.53%] |
| U2 | 30% | Mayotte | 4.06% [0.78%, 12.99%] |
| T1 | 30% | Grande Comore | 76.93% [71.59%, 83.56%] |

**Supplementary Table 1 (continued): Vaccine allocation to islands in the Comoros archipelago under a range of vaccine strategies.** Refer to the initial table caption for the full description.

| Strategy identifier | Percentage of livestock vaccinated annually across the archipelago | Island | Median percentage of vaccines assigned to the island [95% credible interval] |
|---|---|---|---|
| T1 | 30% | Mohéli | 10.64% [5.33%, 14.75%] |
| T1 | 30% | Anjouan | 10.59% [4.03%, 15.33%] |
| T1 | 30% | Mayotte | 1.69% [0.08%, 5.36%] |
| T2 | 30% | Grande Comore | 71.47% [66.16%, 76.94%] |
| T2 | 30% | Mohéli | 11.33% [9.26%, 17.26%] |
| T2 | 30% | Anjouan | 12.31% [9.89%, 18.67%] |
| T2 | 30% | Mayotte | 2.96% [0.96%, 9.48%] |

**Supplementary Table 2: Percentage of livestock vaccinated annually on each island the Comoros archipelago under a range of vaccine strategies**. Assuming animals were not tagged (U) or tagged (T) and given a total percentage of livestock to vaccinate annually across the archipelago, vaccines were allocated to each of the four islands according to optimal infections averted across the archipelago (1) or optimal infections averted on the worst-performing island (2). The table shows the median and 95% credible interval for the percentage of livestock vaccinated annually on each island in the archipelago for each vaccine strategy and six different vaccination rates. The percentage of livestock vaccinated annually on each island proportional to population size (P) are not shown as these are equal to the archipelago vaccination rates. Each median and 95% credible interval was calculated from 500 executions of the optimisation algorithm.

| Strategy identifier | Percentage of livestock vaccinated annually across the archipelago | Island | Median percentage of vaccines assigned to the island [95% credible interval] |
|---|---|---|---|
| U1 | 5% | Grande Comore | 0.53% [0.02%, 1.72%] |
| U1 | 5% | Mohéli | 9.28% [1.57%, 19.68%] |
| U1 | 5% | Anjouan | 14.16% [10.95%, 17.22%] |
| U1 | 5% | Mayotte | 3.69% [0.23%, 10.23%] |
| U2 | 5% | Grande Comore | 4.93% [3.88%, 6.02%] |
| U2 | 5% | Mohéli | 10.83% [6.29%, 19.34%] |
| U2 | 5% | Anjouan | 2.85% [1.27%, 5.40%] |
| U2 | 5% | Mayotte | 4.05% [0.99%, 13.71%] |
| T1 | 5% | Grande Comore | 0.34% [0.01%, 1.43%] |
| T1 | 5% | Mohéli | 16.39% [4.76%, 30.09%] |
| T1 | 5% | Anjouan | 12.40% [7.54%, 16.07%] |
| T1 | 5% | Mayotte | 2.83% [0.12%, 9.16%] |
| T2 | 5% | Grande Comore | 5.16% [4.09%, 5.99%] |
| T2 | 5% | Mohéli | 10.50% [6.62%, 17.57%] |
| T2 | 5% | Anjouan | 2.79% [1.40%, 4.74%] |
| T2 | 5% | Mayotte | 3.44% [1.10%, 11.16%] |
| U1 | 10% | Grande Comore | 6.38% [3.58%, 8.94%] |
| U1 | 10% | Mohéli | 32.04% [15.57%, 41.65%] |
| U1 | 10% | Anjouan | 13.52% [7.15%, 17.53%] |
| U1 | 10% | Mayotte | 4.03% [0.25%, 10.62%] |
| U2 | 10% | Grande Comore | 11.13% [9.20%, 13.01%] |
| U2 | 10% | Mohéli | 17.22% [9.90%, 31.57%] |
| U2 | 10% | Anjouan | 4.49% [2.23%, 8.71%] |
| U2 | 10% | Mayotte | 7.44% [1.67%, 25.10%] |
| T1 | 10% | Grande Comore | 4.20% [2.23%, 7.16%] |
| T1 | 10% | Mohéli | 39.78% [27.81%, 46.49%] |
| T1 | 10% | Anjouan | 14.82% [8.80%, 17.98%] |
| T1 | 10% | Mayotte | 4.15% [0.24%, 12.12%] |
| T2 | 10% | Grande Comore | 11.42% [10.00%, 12.81%] |

**Supplementary Table 2 (continued): Percentage of livestock vaccinated annually on each island the Comoros archipelago under a range of vaccine strategies.** Refer to the initial table caption for the full description.

| Strategy identifier | Percentage of livestock vaccinated annually across the archipelago | Island | Median percentage of livestock vaccinated annually [95% credible interval] |
|---|---|---|---|
| T2 | 10% | Mohéli | 16.41% [10.67%, 26.20%] |
| T2 | 10% | Anjouan | 4.69% [2.86%, 7.73%] |
| T2 | 10% | Mayotte | 5.72% [1.63%, 17.13%] |
| U1 | 15% | Grande Comore | 14.23% [10.98%, 17.19%] |
| U1 | 15% | Mohéli | 36.09% [19.16%, 50.64%] |
| U1 | 15% | Anjouan | 11.95% [6.27%, 18.57%] |
| U1 | 15% | Mayotte | 5.12% [0.24%, 14.18%] |
| U2 | 15% | Grande Comore | 16.90% [14.69%, 19.32%] |
| U2 | 15% | Mohéli | 25.53% [15.61%, 43.17%] |
| U2 | 15% | Anjouan | 6.49% [3.76%, 11.96%] |
| U2 | 15% | Mayotte | 11.42% [2.09%, 35.38%] |
| T1 | 15% | Grande Comore | 15.10% [12.06%, 20.06%] |
| T1 | 15% | Mohéli | 32.31% [20.29%, 43.01%] |
| T1 | 15% | Anjouan | 11.04% [0.67%, 16.48%] |
| T1 | 15% | Mayotte | 4.23% [0.23%, 11.59%] |
| T2 | 15% | Grande Comore | 17.50% [15.87%, 19.29%] |
| T2 | 15% | Mohéli | 22.09% [15.87%, 34.54%] |
| T2 | 15% | Anjouan | 6.85% [4.92%, 10.34%] |
| T2 | 15% | Mayotte | 9.42% [2.34%, 29.18%] |
| U1 | 20% | Grande Comore | 20.49% [17.31%, 24.72%] |
| U1 | 20% | Mohéli | 38.31% [20.36%, 56.00%] |
| U1 | 20% | Anjouan | 15.59% [7.55%, 20.62%] |
| U1 | 20% | Mayotte | 5.77% [0.39%, 16.51%] |
| U2 | 20% | Grande Comore | 22.89% [19.77%, 25.86%] |
| U2 | 20% | Mohéli | 30.53% [19.78%, 52.26%] |
| U2 | 20% | Anjouan | 8.17% [5.29%, 16.12%] |
| U2 | 20% | Mayotte | 16.16% [2.32%, 53.58%] |
| T1 | 20% | Grande Comore | 26.07% [21.37%, 30.67%] |
| T1 | 20% | Mohéli | 31.18% [4.26%, 42.71%] |
| T1 | 20% | Anjouan | 4.55% [0.38%, 14.41%] |
| T1 | 20% | Mayotte | 4.46% [0.16%, 14.27%] |
| T2 | 20% | Grande Comore | 23.62% [21.89%, 25.64%] |
| T2 | 20% | Mohéli | 26.49% [20.46%, 42.25%] |
| T2 | 20% | Anjouan | 9.40% [6.87%, 14.74%] |

**Supplementary Table 2 (continued): Percentage of livestock vaccinated annually on each island the Comoros archipelago under a range of vaccine strategies.** Refer to the initial table caption for the full description.

| Strategy identifier | Percentage of livestock vaccinated annually across the archipelago | Island | Median percentage of livestock vaccinated annually [95% credible interval] |
|---|---|---|---|
| T2 | 20% | Mayotte | 11.42% [2.72%, 36.60%] |
| U1 | 25% | Grande Comore | 28.41% [24.23%, 32.41%] |
| U1 | 25% | Mohéli | 40.17% [18.45%, 63.40%] |
| U1 | 25% | Anjouan | 15.38% [9.04%, 21.09%] |
| U1 | 25% | Mayotte | 7.48% [0.67%, 22.05%] |
| U2 | 25% | Grande Comore | 28.96% [25.39%, 32.71%] |
| U2 | 25% | Mohéli | 33.96% [23.76%, 66.72%] |
| U2 | 25% | Anjouan | 11.13% [7.11%, 22.49%] |
| U2 | 25% | Mayotte | 18.40% [2.47%, 56.91%] |
| T1 | 25% | Grande Comore | 33.97% [29.17%, 38.63%] |
| T1 | 25% | Mohéli | 32.53% [3.73%, 45.19%] |
| T1 | 25% | Anjouan | 5.12% [0.21%, 14.65%] |
| T1 | 25% | Mayotte | 5.97% [0.45%, 19.46%] |
| T2 | 25% | Grande Comore | 29.46% [27.00%, 31.78%] |
| T2 | 25% | Mohéli | 35.21% [26.63%, 54.42%] |
| T2 | 25% | Anjouan | 11.59% [8.82%, 17.79%] |
| T2 | 25% | Mayotte | 13.92% [4.22%, 45.14%] |
| U1 | 30% | Grande Comore | 35.54% [30.96%, 40.77%] |
| U1 | 30% | Mohéli | 47.87% [15.71%, 71.60%] |
| U1 | 30% | Anjouan | 15.43% [8.65%, 21.67%] |
| U1 | 30% | Mayotte | 8.39% [0.47%, 29.02%] |
| U2 | 30% | Grande Comore | 35.05% [30.74%, 38.93%] |
| U2 | 30% | Mohéli | 43.22% [29.49%, 74.68%] |
| U2 | 30% | Anjouan | 12.24% [8.29%, 24.34%] |
| U2 | 30% | Mayotte | 22.47% [4.29%, 71.89%] |
| T1 | 30% | Grande Comore | 38.05% [35.41%, 41.33%] |
| T1 | 30% | Mohéli | 37.04% [18.56%, 51.35%] |
| T1 | 30% | Anjouan | 12.56% [4.78%, 18.17%] |
| T1 | 30% | Mayotte | 9.38% [0.42%, 29.65%] |
| T2 | 30% | Grande Comore | 35.35% [32.72%, 38.06%] |
| T2 | 30% | Mohéli | 39.46% [32.24%, 60.10%] |
| T2 | 30% | Anjouan | 14.59% [11.72%, 22.13%] |
| T2 | 30% | Mayotte | 16.37% [5.31%, 52.46%] |

**Supplementary Table 3: Total infections on each island in the Comoros archipelago without vaccination.** The mathematical model was fitted in a Bayesian framework to seroprevalence surveys from July 2004 until June 2015 (refer to Tennant et al. [1]), yielding a posterior distribution of model parameters. For each sample from this posterior distribution, the model was simulated forward for another 35 years (equivalent to until June 2050) in the absence of any vaccination and the number of infections on each island was calculated. The table shows the median and 95% prediction interval of the total number of infections over 35 years from the model. The median and 95% prediction intervals were calculated using 25,000 samples from the posterior distribution of fitted model parameters.

| Island / archipelago | Median number of infections without any vaccination [95% prediction interval] |
|---|---|
| Grande Comore | 1260356 [1149211, 1346690] |
| Mohéli | 163823 [151393, 175044] |
| Anjouan | 212473 [185217, 233006] |
| Mayotte | 20600 [13078, 29304] |
| Comoros archipelago | 1655932 [1538312, 1749449] |

**Supplementary Table 4: Infections averted across the Comoros archipelago under a range of vaccine strategies.** The mathematical model was simulated forward from the end of the fitting period (June 2015) for 35 years (equivalent to until June 2050) under six different vaccination rates, assuming animals were tagged (`T`) or untagged (`U`), and for three different methods of allocating vaccines between islands: proportional to population size (`P`), optimised in terms of infections averted across the archipelago (`1`) or optimised in terms of infections averted on the worst-performing island (`2`). The table shows the median and 95% prediction interval of the model predicted number infections and percentage of infections averted across the archipelago due to vaccination over the 35 year period for each vaccine strategy. The median and 95% prediction intervals were calculated using 25,000 samples from the distribution of vaccine allocation parameters and posterior of fitted model parameters.

| Strategy identifier | Percentage of livestock vaccinated annually across the archipelago | Median number of infections post-vaccine introduction [95% prediction interval] | Median percentage of infections averted [95% prediction interval] |
|---|---|---|---|
| UP | 5% | 1458068 [1350398, 1544754] | 12.04% [5.94%, 17.55%] |
| U1 | 5% | 1413381 [1308928, 1490439] | 14.82% [8.54%, 20.84%] |
| U2 | 5% | 1451597 [1342787, 1532248] | 12.47% [6.31%, 18.20%] |
| TP | 5% | 1448689 [1336517, 1533834] | 12.60% [6.36%, 18.21%] |
| T1 | 5% | 1381623 [1280147, 1458159] | 16.66% [10.57%, 22.53%] |
| T2 | 5% | 1437097 [1327326, 1519605] | 13.34% [7.12%, 19.09%] |
| UP | 10% | 1255611 [1151742, 1358401] | 24.21% [18.00%, 29.48%] |
| U1 | 10% | 1225275 [1142274, 1320680] | 26.02% [19.80%, 30.37%] |
| U2 | 10% | 1253868 [1157752, 1354478] | 24.32% [17.87%, 29.27%] |
| TP | 10% | 1206201 [1103031, 1305536] | 27.24% [21.07%, 32.36%] |
| T1 | 10% | 1145566 [1056572, 1235806] | 30.80% [24.88%, 35.81%] |
| T2 | 10% | 1192331 [1101001, 1292715] | 28.02% [21.68%, 32.69%] |
| UP | 15% | 1069201 [982056, 1165669] | 35.50% [29.42%, 39.99%] |
| U1 | 15% | 1047443 [967539, 1139447] | 36.83% [30.99%, 40.81%] |
| U2 | 15% | 1068605 [981486, 1168306] | 35.55% [29.27%, 39.95%] |
| TP | 15% | 970902 [889455, 1069595] | 41.28% [35.34%, 45.67%] |
| T1 | 15% | 928340 [850401, 1017426] | 43.92% [38.59%, 48.08%] |
| T2 | 15% | 945963 [862637, 1047709] | 42.77% [36.78%, 47.25%] |
| UP | 20% | 918721 [838738, 1009414] | 44.48% [38.94%, 48.86%] |
| U1 | 20% | 883250 [806351, 971522] | 46.64% [41.37%, 50.74%] |
| U2 | 20% | 910792 [823454, 1014240] | 44.99% [38.85%, 49.66%] |
| TP | 20% | 776351 [705664, 860921] | 52.98% [48.30%, 56.87%] |
| T1 | 20% | 677158 [588650, 775252] | 59.08% [53.54%, 63.67%] |
| T2 | 20% | 707044 [611542, 819922] | 57.30% [50.79%, 62.36%] |
| UP | 25% | 792607 [720883, 874601] | 52.14% [47.18%, 56.01%] |
| U1 | 25% | 735237 [653229, 822990] | 55.61% [50.57%, 59.85%] |
| U2 | 25% | 761706 [668704, 867184] | 54.02% [47.76%, 58.94%] |
| TP | 25% | 606866 [524723, 701344] | 63.38% [58.10%, 67.59%] |
| T1 | 25% | 391266 [308325, 488459] | 76.37% [70.92%, 80.90%] |

**Supplementary Table 4 (continued): Infections averted across the Comoros archipelago under a range of vaccine strategies.** Refer to the initial table caption for the full description.

| Strategy identifier | Percentage of livestock vaccinated annually across the archipelago | Median number of infections post-vaccine introduction [95% prediction interval] | Median percentage of infections averted [95% prediction interval] |
|---|---|---|---|
| T2 | 25% | 435179 [329293, 563404] | 73.74% [66.29%, 79.52%] |
| UP | 30% | 695571 [615363, 777367] | 57.97% [53.17%, 62.24%] |
| U1 | 30% | 599357 [510378, 694436] | 63.82% [58.47%, 68.46%] |
| U2 | 30% | 625943 [527303, 745690] | 62.22% [55.37%, 67.56%] |
| TP | 30% | 416207 [338041, 497006] | 74.83% [70.48%, 79.13%] |
| T1 | 30% | 136204 [54423, 223829] | 91.78% [86.68%, 96.61%] |
| T2 | 30% | 170394 [63170, 289646] | 89.72% [82.71%, 96.08%] |

**Supplementary Table 5: Vaccine efficiency on each island in the Comoros archipelago under a range of vaccine strategies.** Vaccine efficiency was defined as the percentage of vaccines that were administered to animals with no vaccine-induced or natural protection against infection from Rift Valley fever virus. The mathematical model was simulated forward from the end of the fitting period (June 2015) for 35 years (equivalent to until June 2050) under six different vaccination rates, assuming animals were tagged (T) or untagged (U), and for two different methods of allocating vaccines between islands: optimised in terms of infections averted across the archipelago (1) or optimised in terms of infections averted on the worst-performing island (2). The table shows the median and 95% prediction intervals of model predicted mean vaccine efficiency on each island over the 35 year period for each vaccine strategy. The median and 95% prediction intervals were calculated using 25,000 samples from the distribution of vaccine allocation parameters and posterior of fitted model parameters.

| Strategy identifier | Percentage of livestock vaccinated annually across the archipelago | Island | Median vaccine efficiency [95% credible interval] |
|---|---|---|---|
| U1 | 5% | Grande Comore | 62.18% [60.49%, 63.94%] |
| U1 | 5% | Mohéli | 62.87% [60.04%, 65.48%] |
| U1 | 5% | Anjouan | 82.74% [81.18%, 84.64%] |
| U1 | 5% | Mayotte | 91.14% [87.40%, 93.71%] |
| U2 | 5% | Grande Comore | 61.42% [59.83%, 63.26%] |
| U2 | 5% | Mohéli | 62.65% [60.01%, 65.41%] |
| U2 | 5% | Anjouan | 84.30% [82.36%, 86.07%] |
| U2 | 5% | Mayotte | 89.85% [84.30%, 92.36%] |
| T1 | 5% | Grande Comore | 62.46% [60.85%, 64.27%] |
| T1 | 5% | Mohéli | 75.41% [66.05%, 88.42%] |
| T1 | 5% | Anjouan | 95.69% [90.94%, 98.21%] |
| T1 | 5% | Mayotte | 95.63% [91.80%, 98.66%] |
| T2 | 5% | Grande Comore | 65.00% [63.24%, 67.18%] |
| T2 | 5% | Mohéli | 70.76% [66.34%, 76.67%] |
| T2 | 5% | Anjouan | 87.08% [84.64%, 89.41%] |
| T2 | 5% | Mayotte | 94.23% [91.19%, 97.22%] |
| U1 | 10% | Grande Comore | 61.44% [59.82%, 63.08%] |
| U1 | 10% | Mohéli | 60.67% [57.97%, 63.54%] |
| U1 | 10% | Anjouan | 83.39% [81.72%, 85.56%] |
| U1 | 10% | Mayotte | 91.13% [87.16%, 93.80%] |
| U2 | 10% | Grande Comore | 60.69% [59.03%, 62.74%] |
| U2 | 10% | Mohéli | 62.26% [59.57%, 64.85%] |
| U2 | 10% | Anjouan | 84.52% [82.70%, 86.16%] |
| U2 | 10% | Mayotte | 88.67% [76.34%, 92.30%] |
| T1 | 10% | Grande Comore | 65.22% [62.90%, 67.67%] |
| T1 | 10% | Mohéli | 95.33% [85.69%, 96.72%] |
| T1 | 10% | Anjouan | 98.20% [92.77%, 99.08%] |
| T1 | 10% | Mayotte | 97.20% [93.25%, 99.35%] |

**Supplementary Table 5 (continued): Vaccine efficiency on each island in the Comoros archipelago under a range of vaccine strategies.** Refer to the initial table caption for the full description.

| Strategy identifier | Percentage of livestock vaccinated annually across the archipelago | Island | Median vaccine efficiency [95% credible interval] |
|---|---|---|---|
| T2 | 10% | Grande Comore | 69.51% [67.39%, 71.78%] |
| T2 | 10% | Mohéli | 75.89% [70.18%, 84.72%] |
| T2 | 10% | Anjouan | 89.07% [86.75%, 92.05%] |
| T2 | 10% | Mayotte | 96.05% [92.56%, 98.02%] |
| U1 | 15% | Grande Comore | 61.10% [59.24%, 62.93%] |
| U1 | 15% | Mohéli | 60.29% [57.54%, 63.25%] |
| U1 | 15% | Anjouan | 83.88% [81.66%, 86.07%] |
| U1 | 15% | Mayotte | 90.80% [84.90%, 93.96%] |
| U2 | 15% | Grande Comore | 60.87% [59.09%, 62.72%] |
| U2 | 15% | Mohéli | 61.74% [58.44%, 64.41%] |
| U2 | 15% | Anjouan | 84.37% [82.64%, 86.34%] |
| U2 | 15% | Mayotte | 86.44% [70.62%, 92.60%] |
| T1 | 15% | Grande Comore | 73.07% [70.05%, 77.47%] |
| T1 | 15% | Mohéli | 90.97% [79.26%, 96.30%] |
| T1 | 15% | Anjouan | 95.49% [86.04%, 99.01%] |
| T1 | 15% | Mayotte | 96.44% [92.37%, 99.21%] |
| T2 | 15% | Grande Comore | 74.96% [72.48%, 77.43%] |
| T2 | 15% | Mohéli | 81.39% [74.87%, 92.66%] |
| T2 | 15% | Anjouan | 90.96% [88.83%, 95.24%] |
| T2 | 15% | Mayotte | 97.66% [93.76%, 98.70%] |
| U1 | 20% | Grande Comore | 60.44% [58.62%, 62.39%] |
| U1 | 20% | Mohéli | 60.24% [56.71%, 63.55%] |
| U1 | 20% | Anjouan | 83.31% [80.88%, 85.59%] |
| U1 | 20% | Mayotte | 90.84% [83.11%, 94.19%] |
| U2 | 20% | Grande Comore | 60.39% [58.61%, 61.97%] |
| U2 | 20% | Mohéli | 61.17% [57.36%, 63.76%] |
| U2 | 20% | Anjouan | 84.37% [82.43%, 86.30%] |
| U2 | 20% | Mayotte | 83.00% [61.57%, 92.70%] |
| T1 | 20% | Grande Comore | 82.82% [77.60%, 87.71%] |
| T1 | 20% | Mohéli | 89.35% [67.24%, 96.42%] |
| T1 | 20% | Anjouan | 89.65% [85.21%, 98.77%] |
| T1 | 20% | Mayotte | 95.80% [91.70%, 99.16%] |
| T2 | 20% | Grande Comore | 80.21% [77.39%, 83.51%] |
| T2 | 20% | Mohéli | 85.31% [79.48%, 96.36%] |

**Supplementary Table 5 (continued): Vaccine efficiency on each island in the Comoros archipelago under a range of vaccine strategies.** Refer to the initial table caption for the full description.

| Strategy identifier | Percentage of livestock vaccinated annually across the archipelago | Island | Median vaccine efficiency [95% credible interval] |
|---|---|---|---|
| T2 | 20% | Anjouan | 94.33% [90.72%, 98.63%] |
| T2 | 20% | Mayotte | 98.52% [95.06%, 99.25%] |
| U1 | 25% | Grande Comore | 59.84% [58.00%, 61.56%] |
| U1 | 25% | Mohéli | 59.92% [55.76%, 63.84%] |
| U1 | 25% | Anjouan | 83.54% [80.47%, 85.68%] |
| U1 | 25% | Mayotte | 90.03% [79.13%, 93.92%] |
| U2 | 25% | Grande Comore | 59.75% [57.96%, 61.46%] |
| U2 | 25% | Mohéli | 60.62% [55.06%, 63.46%] |
| U2 | 25% | Anjouan | 84.21% [79.76%, 86.20%] |
| U2 | 25% | Mayotte | 81.70% [60.33%, 92.72%] |
| T1 | 25% | Grande Comore | 91.05% [85.65%, 95.49%] |
| T1 | 25% | Mohéli | 91.91% [67.18%, 96.86%] |
| T1 | 25% | Anjouan | 89.93% [85.29%, 99.01%] |
| T1 | 25% | Mayotte | 96.55% [92.18%, 99.25%] |
| T2 | 25% | Grande Comore | 86.58% [83.02%, 89.93%] |
| T2 | 25% | Mohéli | 93.93% [84.58%, 97.14%] |
| T2 | 25% | Anjouan | 96.92% [93.08%, 99.15%] |
| T2 | 25% | Mayotte | 99.03% [96.90%, 99.46%] |
| U1 | 30% | Grande Comore | 59.20% [57.03%, 60.94%] |
| U1 | 30% | Mohéli | 59.02% [53.81%, 63.78%] |
| U1 | 30% | Anjouan | 83.73% [80.27%, 85.93%] |
| U1 | 30% | Mayotte | 89.41% [74.64%, 93.96%] |
| U2 | 30% | Grande Comore | 59.24% [57.15%, 61.01%] |
| U2 | 30% | Mohéli | 59.40% [52.94%, 62.25%] |
| U2 | 30% | Anjouan | 84.22% [78.52%, 86.13%] |
| U2 | 30% | Mayotte | 78.75% [54.89%, 92.05%] |
| T1 | 30% | Grande Comore | 95.65% [92.78%, 97.65%] |
| T1 | 30% | Mohéli | 96.10% [79.41%, 97.62%] |
| T1 | 30% | Anjouan | 97.72% [89.69%, 99.23%] |
| T1 | 30% | Mayotte | 98.83% [93.66%, 99.50%] |
| T2 | 30% | Grande Comore | 93.16% [89.67%, 96.50%] |
| T2 | 30% | Mohéli | 96.68% [91.01%, 97.63%] |
| T2 | 30% | Anjouan | 99.02% [95.95%, 99.26%] |
| T2 | 30% | Mayotte | 99.37% [98.50%, 99.55%] |

**Supplementary Table 6: Total infections on the four islands in the Comoros archipelago under a range of vaccine strategies.** The mathematical model was simulated forward from the end of the fitting period (June 2015) for 35 years (equivalent to until June 2050) under six different vaccination rates, assuming animals were tagged (`T`) or untagged (`U`), and for three different methods of allocating vaccines between islands: proportional to population size (`P`), optimised in terms of infections averted across the archipelago (`1`) or optimised in terms of infections averted on the worst-performing island (`2`). The table shows the median and 95% prediction interval of the model predicted number infections on each island in the archipelago over the 35 year period for each vaccine strategy. The median and 95% prediction intervals were calculated using 25,000 samples from the distribution of vaccine allocation parameters and posterior of fitted model parameters.

| Strategy identifier | Percentage of livestock vaccinated annually across the archipelago | Island | Median number of infections post-vaccine introduction [95% prediction interval] |
|---|---|---|---|
| UP | 5% | Grande Comore | 1150103 [1052533, 1221514] |
| UP | 5% | Mohéli | 148612 [134102, 162090] |
| UP | 5% | Anjouan | 150233 [126610, 174618] |
| UP | 5% | Mayotte | 9454 [5563, 14710] |
| U1 | 5% | Grande Comore | 1218127 [1118332, 1289254] |
| U1 | 5% | Mohéli | 135470 [101531, 162827] |
| U1 | 5% | Anjouan | 49351 [22603, 89854] |
| U1 | 5% | Mayotte | 9166 [1841, 19721] |
| U2 | 5% | Grande Comore | 1144055 [1048062, 1215179] |
| U2 | 5% | Mohéli | 130558 [101734, 151762] |
| U2 | 5% | Anjouan | 166309 [132552, 199059] |
| U2 | 5% | Mayotte | 10858 [3050, 20492] |
| TP | 5% | Grande Comore | 1145327 [1047565, 1216591] |
| TP | 5% | Mohéli | 147616 [133196, 161372] |
| TP | 5% | Anjouan | 147575 [123337, 171052] |
| TP | 5% | Mayotte | 8629 [5267, 13450] |
| T1 | 5% | Grande Comore | 1220176 [1122230, 1288605] |
| T1 | 5% | Mohéli | 103915 [40416, 151579] |
| T1 | 5% | Anjouan | 46125 [13186, 113801] |
| T1 | 5% | Mayotte | 9924 [1444, 19589] |
| T2 | 5% | Grande Comore | 1133664 [1037352, 1204799] |
| T2 | 5% | Mohéli | 127293 [97156, 149451] |
| T2 | 5% | Anjouan | 165220 [134084, 197033] |
| T2 | 5% | Mayotte | 11421 [3129, 20315] |
| UP | 10% | Grande Comore | 1033730 [942669, 1122266] |
| UP | 10% | Mohéli | 129717 [114326, 145911] |
| UP | 10% | Anjouan | 90000 [62901, 116610] |
| UP | 10% | Mayotte | 3351 [2074, 5638] |
| U1 | 10% | Grande Comore | 1086732 [989248, 1193121] |

**Supplementary Table 6 (continued): Total infections on the four islands in the Comoros archipelago under a range of vaccine strategies.** Refer to the initial table caption for the full description.

| Strategy identifier | Percentage of livestock vaccinated annually across the archipelago | Island | Median number of infections post-vaccine introduction [95% prediction interval] |
|---|---|---|---|
| U1 | 10% | Mohéli | 73135 [43558, 116437] |
| U1 | 10% | Anjouan | 48387 [11974, 122611] |
| U1 | 10% | Mayotte | 8109 [1402, 18986] |
| U2 | 10% | Grande Comore | 999158 [909963, 1090691] |
| U2 | 10% | Mohéli | 108358 [70137, 134826] |
| U2 | 10% | Anjouan | 142761 [96656, 176487] |
| U2 | 10% | Mayotte | 6437 [1383, 17111] |
| TP | 10% | Grande Comore | 1003687 [914390, 1093892] |
| TP | 10% | Mohéli | 125376 [110543, 141406] |
| TP | 10% | Anjouan | 74076 [48546, 102390] |
| TP | 10% | Mayotte | 2086 [1357, 3311] |
| T1 | 10% | Grande Comore | 1105325 [1004996, 1205713] |
| T1 | 10% | Mohéli | 7183 [1400, 53571] |
| T1 | 10% | Anjouan | 12789 [1200, 86435] |
| T1 | 10% | Mayotte | 5998 [158, 16825] |
| T2 | 10% | Grande Comore | 950164 [868135, 1041358] |
| T2 | 10% | Mohéli | 100472 [56699, 128805] |
| T2 | 10% | Anjouan | 137626 [96108, 167850] |
| T2 | 10% | Mayotte | 7012 [1321, 16484] |
| UP | 15% | Grande Comore | 919869 [839648, 1011645] |
| UP | 15% | Mohéli | 114480 [99870, 128702] |
| UP | 15% | Anjouan | 33558 [16287, 54633] |
| UP | 15% | Mayotte | 911 [567, 1394] |
| U1 | 15% | Grande Comore | 918037 [817123, 1030928] |
| U1 | 15% | Mohéli | 60279 [26659, 105776] |
| U1 | 15% | Anjouan | 57402 [6804, 126875] |
| U1 | 15% | Mayotte | 6321 [969, 17919] |
| U2 | 15% | Grande Comore | 863927 [782492, 959593] |
| U2 | 15% | Mohéli | 84110 [42402, 117508] |
| U2 | 15% | Anjouan | 122052 [54525, 154339] |
| U2 | 15% | Mayotte | 3130 [877, 14159] |
| TP | 15% | Grande Comore | 852108 [776674, 941511] |
| TP | 15% | Mohéli | 105230 [89931, 120374] |
| TP | 15% | Anjouan | 13905 [4916, 27640] |

**Supplementary Table 6 (continued): Total infections on the four islands in the Comoros archipelago under a range of vaccine strategies.** Refer to the initial table caption for the full description.

| Strategy identifier | Percentage of livestock vaccinated annually across the archipelago | Island | Median number of infections post-vaccine introduction [95% prediction interval] |
|---|---|---|---|
| TP | 15% | Mayotte | 190 [112, 322] |
| T1 | 15% | Grande Comore | 829940 [674605, 958892] |
| T1 | 15% | Mohéli | 28339 [1841, 83354] |
| T1 | 15% | Anjouan | 49618 [2253, 180132] |
| T1 | 15% | Mayotte | 7064 [434, 18148] |
| T2 | 15% | Grande Comore | 766779 [681542, 860087] |
| T2 | 15% | Mohéli | 73751 [18993, 105844] |
| T2 | 15% | Anjouan | 109146 [54738, 139964] |
| T2 | 15% | Mayotte | 2833 [485, 13226] |
| UP | 20% | Grande Comore | 811922 [735996, 896969] |
| UP | 20% | Mohéli | 98345 [84138, 114924] |
| UP | 20% | Anjouan | 7234 [2427, 17154] |
| UP | 20% | Mayotte | 390 [192, 744] |
| U1 | 20% | Grande Comore | 794033 [688411, 897123] |
| U1 | 20% | Mohéli | 55484 [17444, 102030] |
| U1 | 20% | Anjouan | 21264 [2347, 105485] |
| U1 | 20% | Mayotte | 4387 [428, 16250] |
| U2 | 20% | Grande Comore | 749181 [659377, 846964] |
| U2 | 20% | Mohéli | 71573 [22509, 104504] |
| U2 | 20% | Anjouan | 95135 [16331, 136949] |
| U2 | 20% | Mayotte | 1567 [496, 12620] |
| TP | 20% | Grande Comore | 692648 [624987, 772410] |
| TP | 20% | Mohéli | 81143 [68390, 96778] |
| TP | 20% | Anjouan | 2409 [709, 6194] |
| TP | 20% | Mayotte | 31 [17, 51] |
| T1 | 20% | Grande Comore | 510223 [346642, 682795] |
| T1 | 20% | Mohéli | 35041 [1780, 142889] |
| T1 | 20% | Anjouan | 128576 [5797, 188706] |
| T1 | 20% | Mayotte | 7830 [500, 19485] |
| T2 | 20% | Grande Comore | 594641 [489218, 692434] |
| T2 | 20% | Mohéli | 55878 [1882, 84173] |
| T2 | 20% | Anjouan | 65053 [7517, 113959] |
| T2 | 20% | Mayotte | 1145 [149, 10196] |
| UP | 25% | Grande Comore | 706834 [636690, 784463] |

Supplementary Table 6 (continued): **Total infections on the four islands in the Comoros archipelago under a range of vaccine strategies.** Refer to the initial table caption for the full description.

| Strategy identifier | Percentage of livestock vaccinated annually across the archipelago | Island | Median number of infections post-vaccine introduction [95% prediction interval] |
|---|---|---|---|
| UP | 25% | Mohéli | 82507 [70126, 98785] |
| UP | 25% | Anjouan | 2713 [830, 7373] |
| UP | 25% | Mayotte | 317 [129, 683] |
| U1 | 25% | Grande Comore | 651703 [548789, 757498] |
| U1 | 25% | Mohéli | 50823 [9693, 104265] |
| U1 | 25% | Anjouan | 19445 [1564, 90359] |
| U1 | 25% | Mayotte | 2259 [307, 14887] |
| U2 | 25% | Grande Comore | 644623 [545232, 744789] |
| U2 | 25% | Mohéli | 63871 [7031, 92128] |
| U2 | 25% | Anjouan | 61487 [2804, 117978] |
| U2 | 25% | Mayotte | 916 [298, 10497] |
| TP | 25% | Grande Comore | 546168 [464674, 635391] |
| TP | 25% | Mohéli | 60552 [45296, 74120] |
| TP | 25% | Anjouan | 508 [195, 1036] |
| TP | 25% | Mayotte | 10 [4, 21] |
| T1 | 25% | Grande Comore | 228801 [83135, 415630] |
| T1 | 25% | Mohéli | 24840 [495, 145726] |
| T1 | 25% | Anjouan | 124437 [2005, 186957] |
| T1 | 25% | Mayotte | 5963 [284, 18755] |
| T2 | 25% | Grande Comore | 384442 [267262, 506427] |
| T2 | 25% | Mohéli | 13344 [189, 57891] |
| T2 | 25% | Anjouan | 29843 [510, 84734] |
| T2 | 25% | Mayotte | 320 [20, 6329] |
| UP | 30% | Grande Comore | 623676 [545882, 700413] |
| UP | 30% | Mohéli | 71180 [57605, 83301] |
| UP | 30% | Anjouan | 1026 [395, 2344] |
| UP | 30% | Mayotte | 295 [108, 657] |
| U1 | 30% | Grande Comore | 529379 [404535, 638797] |
| U1 | 30% | Mohéli | 31837 [2572, 110895] |
| U1 | 30% | Anjouan | 18723 [809, 94484] |
| U1 | 30% | Mayotte | 1505 [244, 15309] |
| U2 | 30% | Grande Comore | 542493 [429754, 647547] |
| U2 | 30% | Mohéli | 41890 [2201, 77567] |
| U2 | 30% | Anjouan | 49886 [1038, 97667] |

**Supplementary Table 6 (continued): Total infections on the four islands in the Comoros archipelago under a range of vaccine strategies.** Refer to the initial table caption for the full description.

| Strategy identifier | Percentage of livestock vaccinated annually across the archipelago | Island | Median number of infections post-vaccine introduction [95% prediction interval] |
|---|---|---|---|
| U2 | 30% | Mayotte | 652 [213, 6988] |
| TP | 30% | Grande Comore | 380089 [304295, 459925] |
| TP | 30% | Mohéli | 35634 [19993, 51107] |
| TP | 30% | Anjouan | 125 [46, 256] |
| TP | 30% | Mayotte | 5 [2, 12] |
| T1 | 30% | Grande Comore | 79327 [9672, 186439] |
| T1 | 30% | Mohéli | 3762 [73, 87317] |
| T1 | 30% | Anjouan | 19977 [105, 127490] |
| T1 | 30% | Mayotte | 658 [16, 14650] |
| T2 | 30% | Grande Comore | 158687 [47842, 276142] |
| T2 | 30% | Mohéli | 656 [59, 29831] |
| T2 | 30% | Anjouan | 998 [44, 40396] |
| T2 | 30% | Mayotte | 44 [3, 2372] |

**Supplementary Table 7: Infections averted on each island in the Comoros archipelago under a range of vaccine strategies.** The mathematical model was simulated forward from the end of the fitting period (June 2015) for 35 years (equivalent to until June 2050) under six different vaccination rates, assuming animals were tagged (`T`) or untagged (`U`), and for three different methods of allocating vaccines between islands: proportional to population size (`P`), optimised in terms of infections averted across the archipelago (`1`) or optimised in terms of infections averted on the worst-performing island (`2`). The table shows the median and 95% prediction interval of the model predicted percentage of infections averted on each island in the archipelago due to vaccination over the 35 year period for each vaccine strategy. The median and 95% prediction intervals were calculated using 25,000 samples from the distribution of vaccine allocation parameters and posterior of fitted model parameters.

| Strategy identifier | Percentage of livestock vaccinated annually across the archipelago | Island | Median percentage of infections averted [95% prediction interval] |
|---|---|---|---|
| UP | 5% | Grande Comore | 8.86% [0.66%, 16.15%] |
| UP | 5% | Mohéli | 9.05% [5.07%, 14.68%] |
| UP | 5% | Anjouan | 29.17% [19.41%, 37.20%] |
| UP | 5% | Mayotte | 54.10% [46.85%, 61.31%] |
| U1 | 5% | Grande Comore | 3.58% [-5.66%, 11.81%] |
| U1 | 5% | Mohéli | 17.22% [2.04%, 37.03%] |
| U1 | 5% | Anjouan | 76.66% [59.10%, 88.69%] |
| U1 | 5% | Mayotte | 55.50% [17.84%, 89.88%] |
| U2 | 5% | Grande Comore | 9.35% [0.94%, 16.60%] |
| U2 | 5% | Mohéli | 20.03% [10.18%, 36.27%] |
| U2 | 5% | Anjouan | 21.18% [7.60%, 35.33%] |
| U2 | 5% | Mayotte | 46.06% [13.53%, 84.48%] |
| TP | 5% | Grande Comore | 9.25% [0.81%, 16.46%] |
| TP | 5% | Mohéli | 9.64% [5.68%, 15.21%] |
| TP | 5% | Anjouan | 30.65% [20.97%, 38.77%] |
| TP | 5% | Mayotte | 57.22% [50.23%, 64.14%] |
| T1 | 5% | Grande Comore | 3.34% [-5.74%, 11.49%] |
| T1 | 5% | Mohéli | 36.27% [8.70%, 74.82%] |
| T1 | 5% | Anjouan | 78.25% [47.44%, 93.45%] |
| T1 | 5% | Mayotte | 49.85% [16.00%, 92.19%] |
| T2 | 5% | Grande Comore | 10.20% [1.76%, 17.40%] |
| T2 | 5% | Mohéli | 22.00% [11.50%, 39.75%] |
| T2 | 5% | Anjouan | 21.78% [8.40%, 34.78%] |
| T2 | 5% | Mayotte | 42.13% [14.27%, 83.79%] |
| UP | 10% | Grande Comore | 18.08% [9.78%, 24.69%] |
| UP | 10% | Mohéli | 20.70% [15.10%, 26.85%] |
| UP | 10% | Anjouan | 57.40% [47.51%, 67.99%] |
| UP | 10% | Mayotte | 83.36% [79.87%, 87.17%] |
| U1 | 10% | Grande Comore | 13.70% [3.80%, 21.59%] |

**Supplementary Table 7 (continued): Infections averted on each island in the Comoros archipelago under a range of vaccine strategies.** Refer to the initial table caption for the full description.

| Strategy identifier | Percentage of livestock vaccinated annually across the archipelago | Island | Median percentage of infections averted [95% prediction interval] |
|---|---|---|---|
| U1 | 10% | Mohéli | 55.52% [30.08%, 72.34%] |
| U1 | 10% | Anjouan | 77.24% [43.27%, 94.05%] |
| U1 | 10% | Mayotte | 60.21% [20.40%, 92.34%] |
| U2 | 10% | Grande Comore | 20.79% [12.08%, 27.36%] |
| U2 | 10% | Mohéli | 33.64% [19.63%, 56.49%] |
| U2 | 10% | Anjouan | 32.57% [16.76%, 53.43%] |
| U2 | 10% | Mayotte | 68.21% [25.72%, 93.19%] |
| TP | 10% | Grande Comore | 20.41% [12.20%, 26.96%] |
| TP | 10% | Mohéli | 23.29% [17.70%, 29.39%] |
| TP | 10% | Anjouan | 65.07% [54.15%, 75.18%] |
| TP | 10% | Mayotte | 89.47% [86.70%, 92.42%] |
| T1 | 10% | Grande Comore | 12.18% [2.65%, 20.31%] |
| T1 | 10% | Mohéli | 95.62% [67.28%, 99.13%] |
| T1 | 10% | Anjouan | 94.00% [59.54%, 99.41%] |
| T1 | 10% | Mayotte | 69.74% [27.20%, 99.19%] |
| T2 | 10% | Grande Comore | 24.70% [16.20%, 30.75%] |
| T2 | 10% | Mohéli | 38.57% [23.25%, 64.83%] |
| T2 | 10% | Anjouan | 34.97% [20.43%, 53.27%] |
| T2 | 10% | Mayotte | 64.67% [27.68%, 93.48%] |
| UP | 15% | Grande Comore | 27.09% [18.81%, 32.96%] |
| UP | 15% | Mohéli | 30.05% [24.98%, 36.09%] |
| UP | 15% | Anjouan | 84.19% [75.46%, 91.58%] |
| UP | 15% | Mayotte | 95.52% [94.30%, 96.69%] |
| U1 | 15% | Grande Comore | 27.10% [17.36%, 34.89%] |
| U1 | 15% | Mohéli | 63.22% [36.34%, 83.50%] |
| U1 | 15% | Anjouan | 72.91% [41.03%, 96.65%] |
| U1 | 15% | Mayotte | 68.71% [24.39%, 94.92%] |
| U2 | 15% | Grande Comore | 31.48% [22.96%, 37.56%] |
| U2 | 15% | Mohéli | 48.55% [29.84%, 73.61%] |
| U2 | 15% | Anjouan | 42.53% [27.38%, 73.83%] |
| U2 | 15% | Mayotte | 84.44% [37.11%, 95.63%] |
| TP | 15% | Grande Comore | 32.34% [24.30%, 38.15%] |
| TP | 15% | Mohéli | 35.72% [29.76%, 42.64%] |
| TP | 15% | Anjouan | 93.45% [87.63%, 97.43%] |

**Supplementary Table 7 (continued): Infections averted on each island in the Comoros archipelago under a range of vaccine strategies.** Refer to the initial table caption for the full description.

| Strategy identifier | Percentage of livestock vaccinated annually across the archipelago | Island | Median percentage of infections averted [95% prediction interval] |
|---|---|---|---|
| TP | 15% | Mayotte | 99.04% [98.18%, 99.53%] |
| T1 | 15% | Grande Comore | 34.00% [23.42%, 46.30%] |
| T1 | 15% | Mohéli | 82.70% [49.49%, 98.86%] |
| T1 | 15% | Anjouan | 76.71% [14.83%, 98.89%] |
| T1 | 15% | Mayotte | 65.21% [19.35%, 97.64%] |
| T2 | 15% | Grande Comore | 39.06% [31.21%, 45.59%] |
| T2 | 15% | Mohéli | 54.75% [37.00%, 88.23%] |
| T2 | 15% | Anjouan | 48.31% [34.41%, 73.33%] |
| T2 | 15% | Mayotte | 85.51% [42.91%, 97.72%] |
| UP | 20% | Grande Comore | 35.48% [27.83%, 41.27%] |
| UP | 20% | Mohéli | 39.91% [33.22%, 46.10%] |
| UP | 20% | Anjouan | 96.61% [92.18%, 98.75%] |
| UP | 20% | Mayotte | 98.12% [97.34%, 98.62%] |
| U1 | 20% | Grande Comore | 36.93% [28.20%, 45.17%] |
| U1 | 20% | Mohéli | 66.40% [38.85%, 89.11%] |
| U1 | 20% | Anjouan | 89.94% [50.53%, 98.85%] |
| U1 | 20% | Mayotte | 78.27% [31.81%, 97.66%] |
| U2 | 20% | Grande Comore | 40.51% [32.14%, 47.27%] |
| U2 | 20% | Mohéli | 56.19% [37.81%, 85.93%] |
| U2 | 20% | Anjouan | 54.69% [36.14%, 92.16%] |
| U2 | 20% | Mayotte | 92.18% [44.94%, 97.51%] |
| TP | 20% | Grande Comore | 44.83% [38.47%, 50.01%] |
| TP | 20% | Mohéli | 50.44% [44.11%, 56.15%] |
| TP | 20% | Anjouan | 98.86% [97.19%, 99.64%] |
| TP | 20% | Mayotte | 99.86% [99.71%, 99.91%] |
| T1 | 20% | Grande Comore | 59.50% [45.88%, 71.99%] |
| T1 | 20% | Mohéli | 78.71% [13.13%, 98.90%] |
| T1 | 20% | Anjouan | 39.32% [10.97%, 97.23%] |
| T1 | 20% | Mayotte | 61.12% [12.37%, 97.44%] |
| T2 | 20% | Grande Comore | 52.80% [45.03%, 60.45%] |
| T2 | 20% | Mohéli | 65.73% [49.87%, 98.84%] |
| T2 | 20% | Anjouan | 68.97% [47.82%, 96.30%] |
| T2 | 20% | Mayotte | 94.05% [55.75%, 99.32%] |
| UP | 25% | Grande Comore | 43.95% [37.31%, 49.01%] |

**Supplementary Table 7 (continued): Infections averted on each island in the Comoros archipelago under a range of vaccine strategies.** Refer to the initial table caption for the full description.

| Strategy identifier | Percentage of livestock vaccinated annually across the archipelago | Island | Median percentage of infections averted [95% prediction interval] |
|---|---|---|---|
| UP | 25% | Mohéli | 49.49% [42.75%, 54.93%] |
| UP | 25% | Anjouan | 98.72% [96.71%, 99.57%] |
| UP | 25% | Mayotte | 98.47% [97.56%, 99.06%] |
| U1 | 25% | Grande Comore | 48.26% [39.65%, 55.94%] |
| U1 | 25% | Mohéli | 69.01% [37.24%, 93.97%] |
| U1 | 25% | Anjouan | 90.83% [57.99%, 99.23%] |
| U1 | 25% | Mayotte | 88.68% [36.52%, 98.23%] |
| U2 | 25% | Grande Comore | 48.86% [40.63%, 56.20%] |
| U2 | 25% | Mohéli | 60.84% [45.40%, 95.64%] |
| U2 | 25% | Anjouan | 70.83% [46.03%, 98.66%] |
| U2 | 25% | Mayotte | 95.62% [52.90%, 98.34%] |
| TP | 25% | Grande Comore | 56.73% [49.70%, 62.19%] |
| TP | 25% | Mohéli | 62.86% [57.27%, 70.92%] |
| TP | 25% | Anjouan | 99.76% [99.53%, 99.90%] |
| TP | 25% | Mayotte | 99.95% [99.92%, 99.97%] |
| T1 | 25% | Grande Comore | 81.83% [67.27%, 93.30%] |
| T1 | 25% | Mohéli | 84.82% [11.10%, 99.69%] |
| T1 | 25% | Anjouan | 41.50% [11.32%, 99.04%] |
| T1 | 25% | Mayotte | 70.00% [15.19%, 98.56%] |
| T2 | 25% | Grande Comore | 69.48% [60.14%, 78.23%] |
| T2 | 25% | Mohéli | 91.78% [65.49%, 99.88%] |
| T2 | 25% | Anjouan | 85.71% [61.23%, 99.75%] |
| T2 | 25% | Mayotte | 98.39% [72.44%, 99.90%] |
| UP | 30% | Grande Comore | 50.49% [44.08%, 55.95%] |
| UP | 30% | Mohéli | 56.49% [51.52%, 63.14%] |
| UP | 30% | Anjouan | 99.51% [98.94%, 99.80%] |
| UP | 30% | Mayotte | 98.58% [97.65%, 99.21%] |
| U1 | 30% | Grande Comore | 57.94% [49.47%, 67.33%] |
| U1 | 30% | Mohéli | 80.61% [33.04%, 98.40%] |
| U1 | 30% | Anjouan | 91.13% [56.35%, 99.60%] |
| U1 | 30% | Mayotte | 92.49% [33.48%, 98.55%] |
| U2 | 30% | Grande Comore | 56.92% [48.60%, 65.33%] |
| U2 | 30% | Mohéli | 74.30% [53.77%, 98.64%] |
| U2 | 30% | Anjouan | 76.35% [54.93%, 99.50%] |

**Supplementary Table 7 (continued): Infections averted on each island in the Comoros archipelago under a range of vaccine strategies.** Refer to the initial table caption for the full description.

| Strategy identifier | Percentage of livestock vaccinated annually across the archipelago | Island | Median percentage of infections averted [95% prediction interval] |
|---|---|---|---|
| U2 | 30% | Mayotte | 96.95% [68.80%, 98.73%] |
| TP | 30% | Grande Comore | 69.76% [63.99%, 75.25%] |
| TP | 30% | Mohéli | 78.25% [70.58%, 86.93%] |
| TP | 30% | Anjouan | 99.94% [99.88%, 99.98%] |
| TP | 30% | Mayotte | 99.98% [99.96%, 99.99%] |
| T1 | 30% | Grande Comore | 93.70% [85.37%, 99.22%] |
| T1 | 30% | Mohéli | 97.69% [46.32%, 99.95%] |
| T1 | 30% | Anjouan | 90.57% [40.24%, 99.95%] |
| T1 | 30% | Mayotte | 96.58% [34.01%, 99.92%] |
| T2 | 30% | Grande Comore | 87.40% [78.29%, 96.08%] |
| T2 | 30% | Mohéli | 99.60% [82.26%, 99.96%] |
| T2 | 30% | Anjouan | 99.53% [81.54%, 99.98%] |
| T2 | 30% | Mayotte | 99.79% [89.80%, 99.99%] |

**Supplementary Table 8: Deviation in infections averted on each island in the Comoros archipelago under a range of vaccine strategies.** The mathematical model was simulated forward from the end of the fitting period (June 2015) for 35 years (equivalent to until June 2050) under six different vaccination rates, assuming animals were tagged (`T`) or untagged (`U`), and for two different methods of allocating vaccines between islands: optimised in terms of infections averted across the archipelago (`1`) or optimised in terms of infections averted on the worst-performing island (`2`). The table shows the median and 95% prediction intervals of model predicted deviation in the percentage of infections averted on each island compared to the mean percentage of infections averted across all islands over the 35 year period for each vaccine strategy. The median and 95% prediction intervals were calculated using 25,000 samples from the distribution of vaccine allocation parameters and posterior of fitted model parameters.

| Strategy identifier | Percentage of livestock vaccinated annually across the archipelago | Island | Median deviation in percentage of infections averted [95% prediction interval] |
|---|---|---|---|
| `U1` | 5% | Grande Comore | -34.69% [-45.99%, -21.62%] |
| `U1` | 5% | Mohéli | -20.89% [-38.49%, 0.32%] |
| `U1` | 5% | Anjouan | 38.21% [20.3%, 53.65%] |
| `U1` | 5% | Mayotte | 17.07% [-12.22%, 44.88%] |
| `U2` | 5% | Grande Comore | -15.42% [-26.89%, -4.09%] |
| `U2` | 5% | Mohéli | -4% [-17.81%, 12.04%] |
| `U2` | 5% | Anjouan | -3.65% [-18.35%, 12.02%] |
| `U2` | 5% | Mayotte | 21.39% [-3.79%, 52.17%] |
| `T1` | 5% | Grande Comore | -39.22% [-50.33%, -27.57%] |
| `T1` | 5% | Mohéli | -5.87% [-35.62%, 33.03%] |
| `T1` | 5% | Anjouan | 34.75% [4.51%, 54.43%] |
| `T1` | 5% | Mayotte | 7.96% [-20.29%, 42.88%] |
| `T2` | 5% | Grande Comore | -14.32% [-26.42%, -4.19%] |
| `T2` | 5% | Mohéli | -2.03% [-16.34%, 14.61%] |
| `T2` | 5% | Anjouan | -3.02% [-17.6%, 11.36%] |
| `T2` | 5% | Mayotte | 17.72% [-4.65%, 50.62%] |
| `U1` | 10% | Grande Comore | -36.77% [-53.3%, -18.61%] |
| `U1` | 10% | Mohéli | 4.52% [-19.46%, 24.56%] |
| `U1` | 10% | Anjouan | 25.68% [-1.6%, 43.83%] |
| `U1` | 10% | Mayotte | 9.86% [-19.82%, 34.08%] |
| `U2` | 10% | Grande Comore | -18.27% [-29.94%, -4.7%] |
| `U2` | 10% | Mohéli | -4.71% [-18.51%, 16.52%] |
| `U2` | 10% | Anjouan | -6.19% [-21.8%, 15.54%] |
| `U2` | 10% | Mayotte | 29.01% [-4.41%, 50.96%] |
| `T1` | 10% | Grande Comore | -54.92% [-68.79%, -34.02%] |
| `T1` | 10% | Mohéli | 26.87% [9.85%, 42.31%] |
| `T1` | 10% | Anjouan | 24.06% [3.05%, 37.54%] |
| `T1` | 10% | Mayotte | 4.05% [-27.3%, 25.28%] |
| `T2` | 10% | Grande Comore | -16.58% [-28.13%, -4.22%] |

**Supplementary Table 8 (continued): Deviation in infections averted on each island in the Comoros archipelago under a range of vaccine strategies.** Refer to the initial table caption for the full description.

| Strategy identifier | Percentage of livestock vaccinated annually across the archipelago | Island | Median deviation in percentage of infections averted [95% prediction interval] |
|---|---|---|---|
| T2 | 10% | Mohéli | -1.65% [-18.18%, 22.64%] |
| T2 | 10% | Anjouan | -6.01% [-21.17%, 13.68%] |
| T2 | 10% | Mayotte | 23.39% [-6.61%, 48.39%] |
| U1 | 15% | Grande Comore | -30.09% [-47.19%, -10.86%] |
| U1 | 15% | Mohéli | 5.78% [-19.23%, 26.73%] |
| U1 | 15% | Anjouan | 15.95% [-9.78%, 39.06%] |
| U1 | 15% | Mayotte | 11.89% [-22.95%, 31.67%] |
| U2 | 15% | Grande Comore | -19.94% [-32.14%, -5.4%] |
| U2 | 15% | Mohéli | -2.19% [-20.6%, 22.67%] |
| U2 | 15% | Anjouan | -7.87% [-22.22%, 20.31%] |
| U2 | 15% | Mayotte | 31.48% [-5.83%, 44.69%] |
| T1 | 15% | Grande Comore | -29.31% [-51.35%, 4.17%] |
| T1 | 15% | Mohéli | 18.88% [-7.87%, 41.89%] |
| T1 | 15% | Anjouan | 11.91% [-32.68%, 34.59%] |
| T1 | 15% | Mayotte | 3.23% [-28.82%, 26.31%] |
| T2 | 15% | Grande Comore | -17.82% [-30.26%, -3.02%] |
| T2 | 15% | Mohéli | -0.78% [-18.1%, 28.46%] |
| T2 | 15% | Anjouan | -7.8% [-21.52%, 16.31%] |
| T2 | 15% | Mayotte | 26.84% [-7.36%, 41.7%] |
| U1 | 20% | Grande Comore | -29.62% [-44.98%, -7.82%] |
| U1 | 20% | Mohéli | -0.17% [-23.31%, 22.88%] |
| U1 | 20% | Anjouan | 21.57% [-5.51%, 36.6%] |
| U1 | 20% | Mayotte | 12.09% [-24.25%, 28.61%] |
| U2 | 20% | Grande Comore | -19.73% [-33.29%, -5.83%] |
| U2 | 20% | Mohéli | -3.34% [-22.52%, 24.52%] |
| U2 | 20% | Anjouan | -5.37% [-21.44%, 28.25%] |
| U2 | 20% | Mayotte | 29.51% [-9.13%, 40.24%] |
| T1 | 20% | Grande Comore | 0.07% [-34.11%, 34.5%] |
| T1 | 20% | Mohéli | 13.69% [-28.51%, 42.38%] |
| T1 | 20% | Anjouan | -14.58% [-43.93%, 27.42%] |
| T1 | 20% | Mayotte | 0.64% [-33.72%, 31.13%] |
| T2 | 20% | Grande Comore | -17.61% [-29.65%, -3.99%] |
| T2 | 20% | Mohéli | -3.62% [-19.48%, 26.1%] |
| T2 | 20% | Anjouan | -1.49% [-18.85%, 24.15%] |

**Supplementary Table 8 (continued): Deviation in infections averted on each island in the Comoros archipelago under a range of vaccine strategies.** Refer to the initial table caption for the full description.

| Strategy identifier | Percentage of livestock vaccinated annually across the archipelago | Island | Median deviation in percentage of infections averted [95% prediction interval] |
|---|---|---|---|
| T2 | 20% | Mayotte | 21.62% [-10.59%, 32.55%] |
| U1 | 25% | Grande Comore | -23.23% [-39.45%, -3.32%] |
| U1 | 25% | Mohéli | -2.76% [-27.24%, 21.46%] |
| U1 | 25% | Anjouan | 17.22% [-7.25%, 34.07%] |
| U1 | 25% | Mayotte | 15.03% [-24.9%, 27.53%] |
| U2 | 25% | Grande Comore | -20.33% [-33.42%, -6.31%] |
| U2 | 25% | Mohéli | -7.37% [-23.87%, 24.36%] |
| U2 | 25% | Anjouan | 2.19% [-18.11%, 27.37%] |
| U2 | 25% | Mayotte | 23.84% [-10.5%, 33.8%] |
| T1 | 25% | Grande Comore | 12.8% [-18.77%, 46.11%] |
| T1 | 25% | Mohéli | 11.06% [-40.61%, 36.44%] |
| T1 | 25% | Anjouan | -21.15% [-50.48%, 20.59%] |
| T1 | 25% | Mayotte | 1.59% [-39.14%, 27.95%] |
| T2 | 25% | Grande Comore | -15.22% [-26.36%, -1.66%] |
| T2 | 25% | Mohéli | 6.18% [-15.51%, 19.14%] |
| T2 | 25% | Anjouan | 0.19% [-18.26%, 15.72%] |
| T2 | 25% | Mayotte | 11.97% [-8.68%, 21.16%] |
| U1 | 30% | Grande Comore | -18.85% [-34.59%, 7.12%] |
| U1 | 30% | Mohéli | 3.14% [-34.56%, 24.33%] |
| U1 | 30% | Anjouan | 12.95% [-12.87%, 31.53%] |
| U1 | 30% | Mayotte | 12.02% [-29.38%, 26.08%] |
| U2 | 30% | Grande Comore | -19.33% [-32.07%, -5.67%] |
| U2 | 30% | Mohéli | -1.77% [-20.47%, 20.83%] |
| U2 | 30% | Anjouan | 0.98% [-17.04%, 22.39%] |
| U2 | 30% | Mayotte | 18.77% [-4.65%, 28.71%] |
| T1 | 30% | Grande Comore | 4.19% [-9.63%, 26.09%] |
| T1 | 30% | Mohéli | 3.78% [-34.31%, 23.97%] |
| T1 | 30% | Anjouan | 0.17% [-38.11%, 17.17%] |
| T1 | 30% | Mayotte | 3.2% [-43%, 15.55%] |
| T2 | 30% | Grande Comore | -7.91% [-15.33%, 1.61%] |
| T2 | 30% | Mohéli | 3.11% [-9.49%, 8.25%] |
| T2 | 30% | Anjouan | 2.99% [-10.06%, 7.77%] |
| T2 | 30% | Mayotte | 4.07% [-4.05%, 8.91%] |

**Supplementary Table 9: Mathematical notation for livestock infection model.** A mathematical model describing livestock infection with and vaccination against Rift Valley Fever virus (RVFV) across the four islands in the Comoros archipelago was developed. In order to determine how best to distribute a set number of vaccines across the archipelago, a Sequential Monte Carlo (SMC) optimisation algorithm was used, where the objective function was the number of infections averted between July 2015 and June 2050. The table below shows all mathematical notation used to describe the model.

| Notation | Description |
|---|---|
| $S^{\square}$ | Number of susceptible livestock |
| $E^{\square}$ | Number of exposed (infected, but not yet infectious) livestock |
| $I^{\square}$ | Number of infectious livestock |
| $R^{\square}$ | Number of recovered (with life-long natural immunity) livestock |
| $\square^{U}$ | Unvaccinated livestock |
| $\square^{V_1}$ | Vaccinated livestock which are not yet protected (week 1) |
| $\square^{V_2}$ | Vaccinated livestock which are not yet protected (week 2) |
| $\square^{W}$ | Vaccinated and protected livestock |
| $\square^{\text{ext}}$ | Externally introduced livestock |
| $t$ | Time (weeks) |
| $i$ | Island |
| $a$ | Age group |
| $n$ | Number of islands in the metapopulation |
| $A$ | Number of age groups |
| $A^{V}$ | Maximum age group that can be vaccinated |
| $A^{\text{move}}$ | Maximum age group that could be moved between islands |
| $A^{\text{ext}}$ | Maximum age group of livestock that are externally introduced |
| $t_V$ | Time that vaccination begins |
| $T$ | Maximum simulation time |
| $p_a$ | Initial proportion of livestock in age group $a$ |
| $\nu_{t,i}$ | Number of births at time $t$ on island $i$ |
| $\mu_a$ | Probability of dying in age group $a$ per week |
| $\delta_a$ | Probability of ageing out from age group $a$ per week |
| $m_{j,i,a}$ | Probability of livestock from age group $a$ moving from island $j$ to island $i$ per week |
| $\lambda_{t,i}$ | Probability of becoming infected on island $i$ at time $t$ |
| $\beta_{t,i}$ | Disease transmission rate per week |
| $\alpha$ | Influence of Normalised Difference Vegetation Index on the disease transmission rate |
| $\gamma_i$ | Natural log of the minimum disease transmission rate on island $i$ per week |
| $p^{\text{eff}}$ | Vaccine efficacy |
| $\tau_\omega$ | Mean duration of vaccine-induced immunity (weeks) |
| $\omega$ | Probability of vaccine-induced immunity waning per week |
| $\xi_{t,i,a}$ | Probability of individual from age group $a$ and island $i$ at time $t$ being vaccinated |
| $\psi$ | Proportion of livestock vaccinated across the metapopulation per week |
| $\rho_i$ | Proportion of vaccines allocated to island $i$ |

**Supplementary Table 10: Summary of estimated parameters used during model simulations.** Prior to simulating forward the transmission model in time, a single parameter set is sampled from the posterior distribution of the parameters of interest. These parameters were estimated by fitting to data in a Bayesian framework. The table below shows the the median and 95% credible intervals (CrI) of 10,000 posterior samples. For further details on the model fitting procedure, refer to Tennant et al. [1].

| Parameter | Description | Median | [95% CrI] |
|---|---|---|---|
| $48N_1m_{12}$ | Annual movement from Grande Comore to Mohéli | 329.33 | [266.54, 391.15] |
| $48N_1m_{13}$ | Annual movement from Grande Comore to Anjouan | 65.11 | [7.82, 137.33] |
| $48N_2m_{21}$ | Annual movement from Mohéli to Grande Comore | 418.27 | [321.79, 517.59] |
| $48N_2m_{23}$ | Annual movement from Mohéli to Anjouan | 73.44 | [18.43, 139.73] |
| $48N_3m_{31}$ | Annual movement from Anjouan to Grande Comore | 624.58 | [515.76, 735.8] |
| $48N_3m_{32}$ | Annual movement from Anjouan to Mohéli | 407.93 | [339.02, 476.64] |
| $48N_3m_{34}$ | Annual movement from Anjouan to Mayotte | 1896.24 | [1653.36, 2126.16] |
| $\epsilon_1$ | Proportion immune at time $t = 0$ in Grande Comore | 0.372 | [0.284, 0.429] |
| $\epsilon_2$ | Proportion immune at time $t = 0$ in Mohéli | 0.397 | [0.346, 0.462] |
| $\epsilon_3$ | Proportion immune at time $t = 0$ in Anjouan | 0.035 | [0.015, 0.063] |
| $\epsilon_4$ | Proportion immune at time $t = 0$ in Mayotte | 0.141 | [0.107, 0.178] |
| $\alpha$ | Seasonal transmission component scalar for all islands | 7.73 | [7.09, 8.42] |
| $\gamma_1$ | Transmission constant for Grande Comore | -0.79 | [-0.89, -0.69] |
| $\gamma_2$ | Transmission constant for Mohéli | -0.79 | [-0.91, -0.67] |
| $\gamma_3$ | Transmission constant for Anjouan | -1.06 | [-1.18, -0.96] |
| $\gamma_4$ | Transmission constant for Mayotte | -1.02 | [-1.14, -0.92] |
| $t^{\text{ext}}_{\text{(start)}}$ | Start of imports into Grande Comore (epidemiological weeks since July 2004) | 130.69 | [116.94, 151.70] |
| $t^{\text{ext}}_{\text{(duration)}}$ | Duration of imports into Grande Comore | 23.13 | [5.15, 40.68] |
| $48\iota^{\text{ext}}_1$ | Annual infectious imports into Grande Comore | 175.34 | [13.82, 414.82] |

#### Supplementary Figures

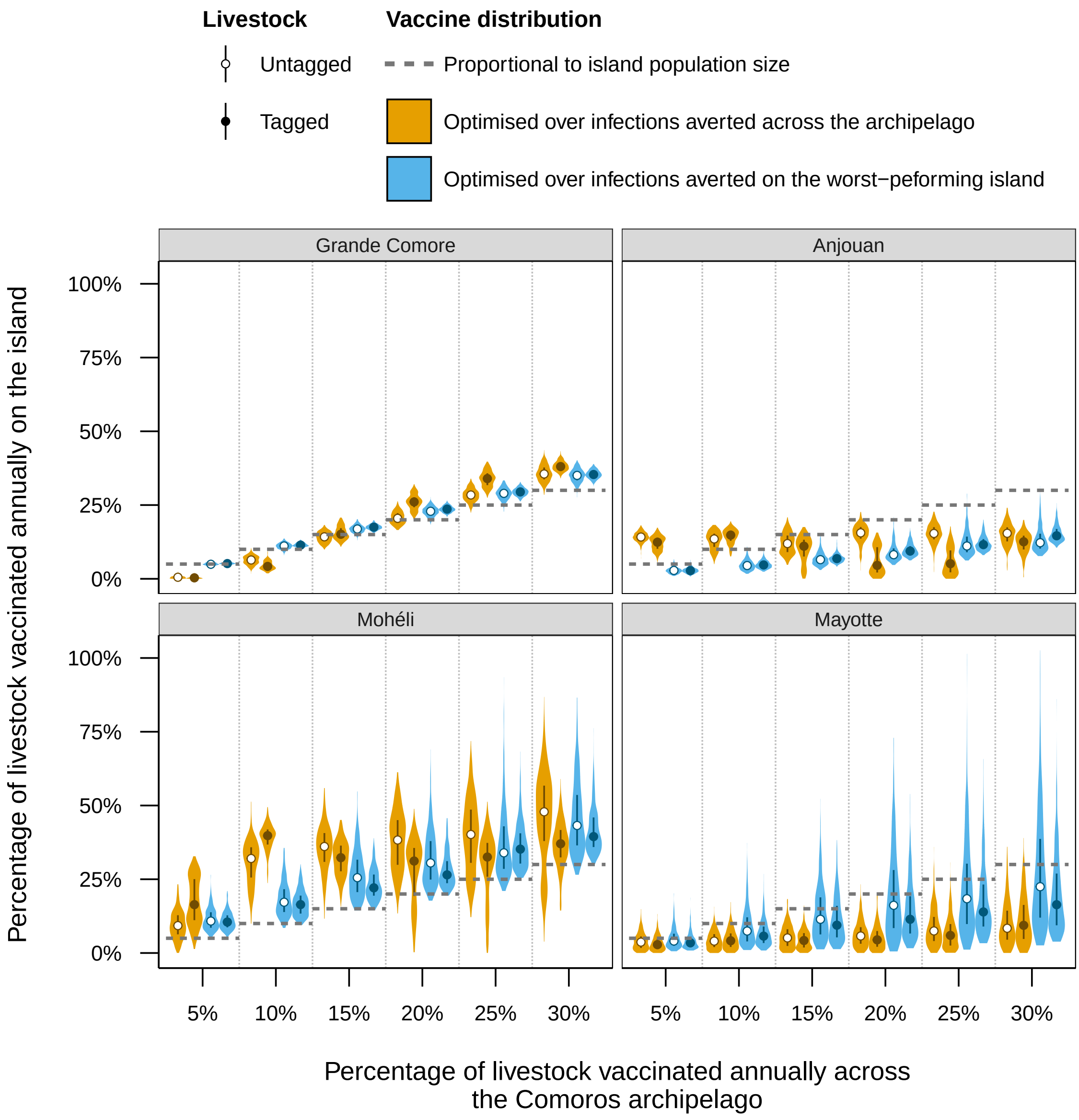


**Supplementary Figure 1: Animals vaccinated on each island in the Comoros archipelago for different vaccine strategies.** Vaccines were allocated to each of the four islands in the archipelago either proportionally to the livestock population size of each island (grey dashed line), optimally to maximise the percentage of infections averted across the archipelago (orange violins), or the percentage of infections averted on the island with the the worst performance (blue violins). For both optimal vaccine allocations, all vaccination rates and livestock tagging strategies, the median percentage of livestock vaccinated on Mohéli was greater than the overall percentage of livestock vaccinated across the archipelago, indicating overall favour towards vaccinating Mohéli. The violins show the percentage of animals vaccinated annually on each island for different annual vaccination rates, allocation methods and tagging strategies. The points and boxplots show the median and inter-quartile range for each scenario respectively. All violins shown are based on 500 executions of the optimisation algorithm. Refer to Supplementary Table 2 for detailed numerical results.

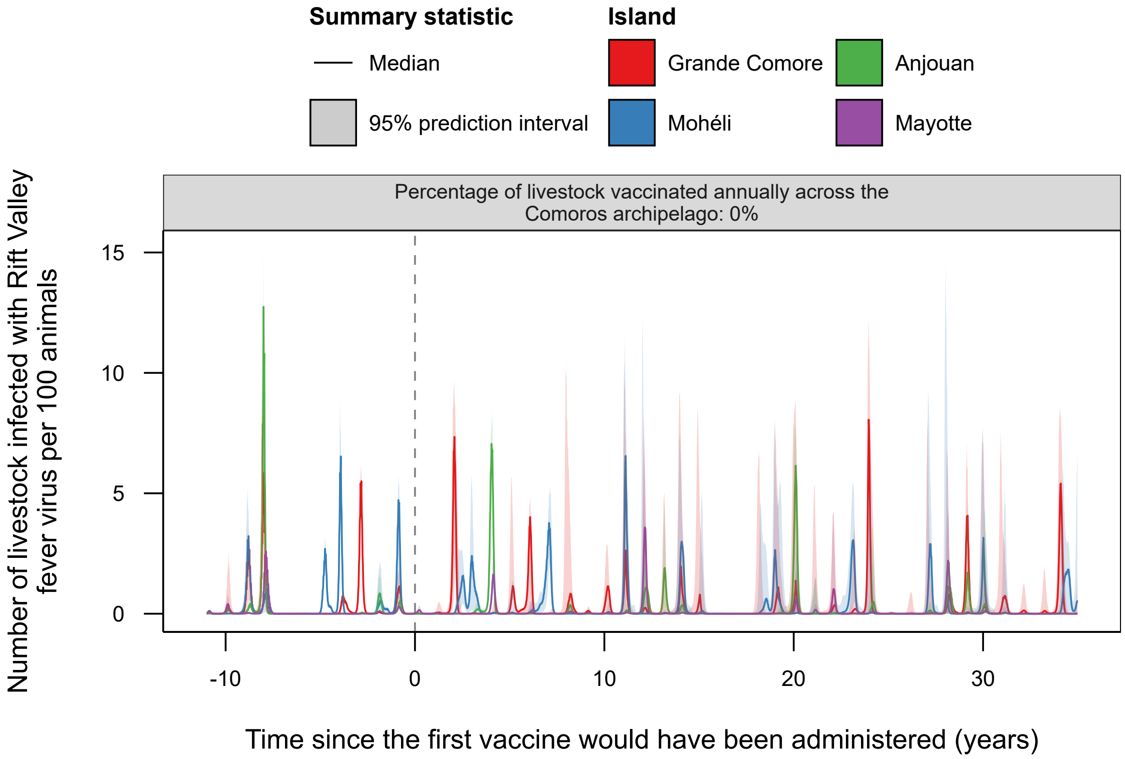


**Supplementary Figure 2: Model predicted number of infected livestock per island in the Comoros archipelago without vaccination.** The metapopulation model describing Rift Valley Fever virus (RVFV) infection in livestock on each island of the Comoros archipelago—Grande Comore (red), Mohéli (blue), Anjouan (green) and Mayotte (purple)—was fitted to serological surveys conducted between July 2004 and June 2015. Shown is the median (solid line) and 95% prediction interval (light area) of model predicted number of infections on each island for 35 years after June 2015—the time when the vaccine was administered in simulations with vaccine. This established a baseline number of infections for comparison with different vaccination strategies against RVFV in the archipelago. The median and 95% prediction intervals were calculated from 1000 model simulations.

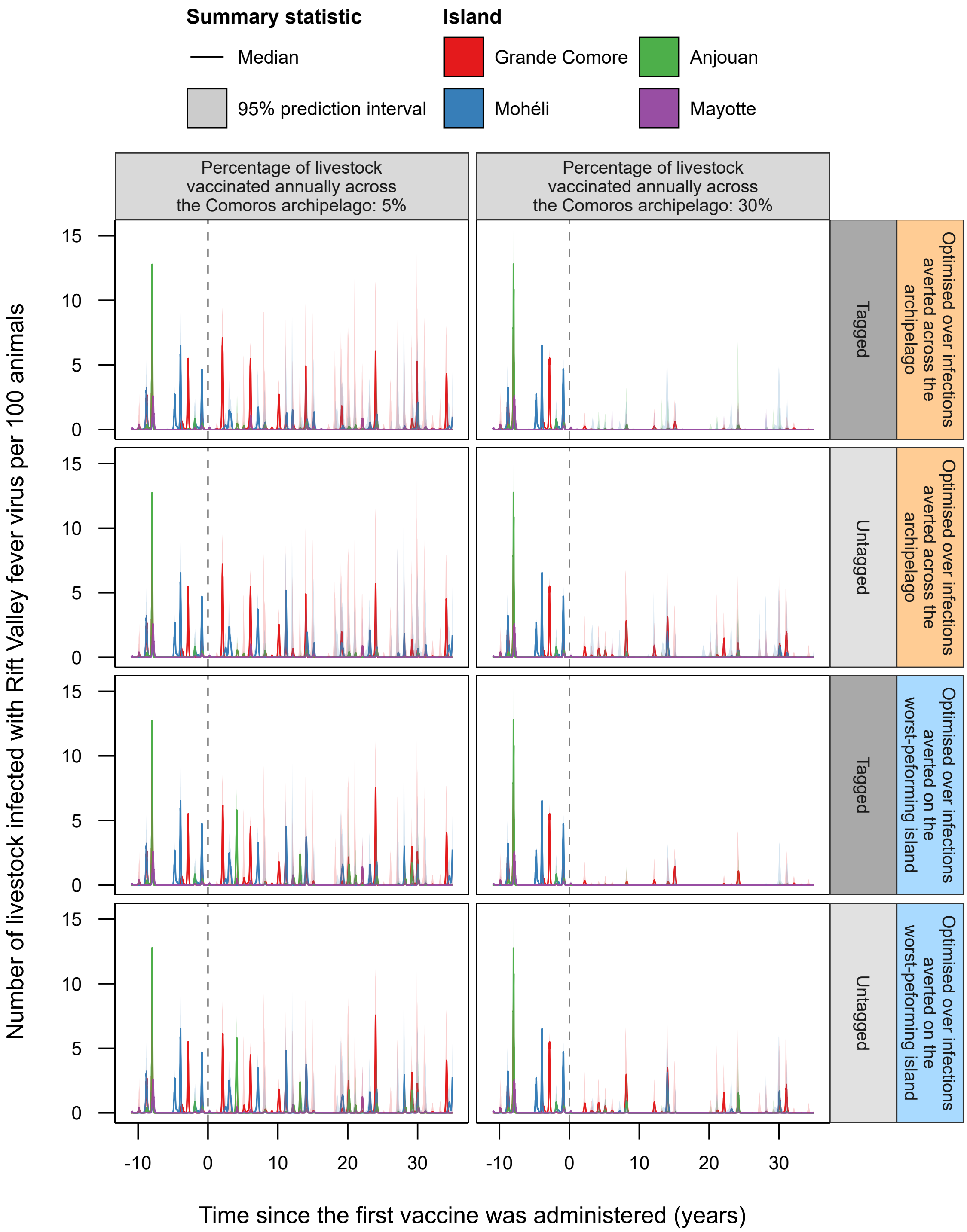


**Supplementary Figure 3: Model predicted number of infected livestock per island in the Comoros archipelago with 5% and 30% vaccination rates.** The fitted metapopulation model describing Rift Valley Fever virus (RVFV) infection in livestock on each island of the Comoros archipelago—Grande Comore (red), Mohéli (blue), Anjouan (green) and Mayotte (purple)—was simulated forward in time for 35 years from the end of the fitting period, June 2015, under a range of vaccination strategies. Shown is the median (solid line) and 95% prediction interval (light area) of model predicted number of infections on each island for 5% and 30% of livestock vaccinated annually across the archipelago, optimal vaccine allocation across islands, and two livestock tagging strategies. This was compared to the model predicted number of infections without vaccination and then used to establish the effectiveness of a given strategy. The median and 95% prediction intervals were calculated from 1000 model simulations.

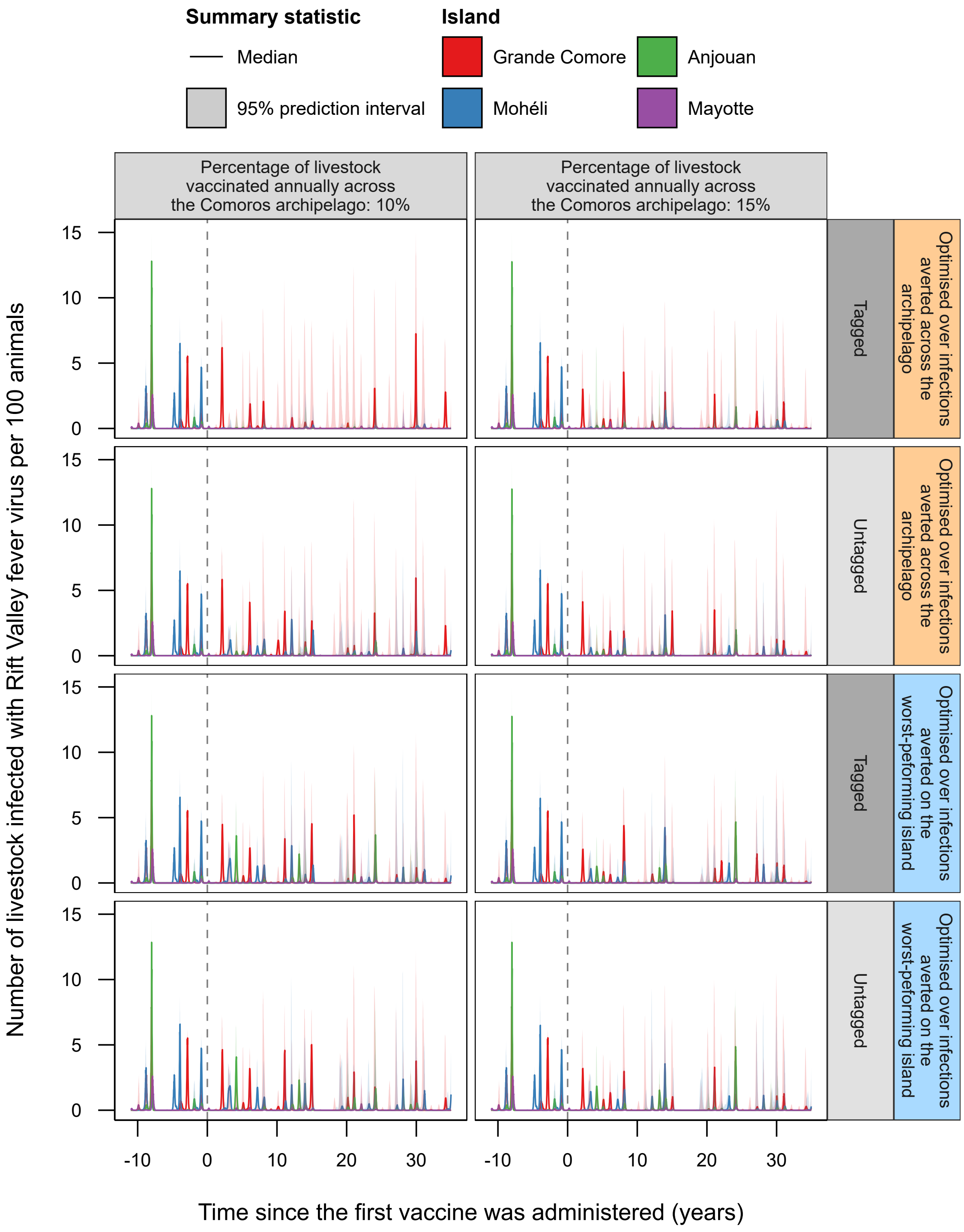


**Supplementary Figure 4: Model predicted number of infected livestock per island in the Comoros archipelago with 10% and 15% vaccination rates.** The fitted metapopulation model describing Rift Valley Fever virus (RVFV) infection in livestock on each island of the Comoros archipelago—Grande Comore (red), Mohéli (blue), Anjouan (green) and Mayotte (purple)—was simulated forward in time for 35 years from the end of the fitting period, June 2015, under a range of vaccination strategies. Shown is the median (solid line) and 95% prediction interval (light area) of model predicted number of infections on each island for 10% and 15% of livestock vaccinated annually across the archipelago, optimal vaccine allocation across islands, and two livestock tagging strategies. This was compared to the model predicted number of infections without vaccination and then used to establish the effectiveness of a given strategy. The median and 95% prediction intervals were calculated from 1000 model simulations.

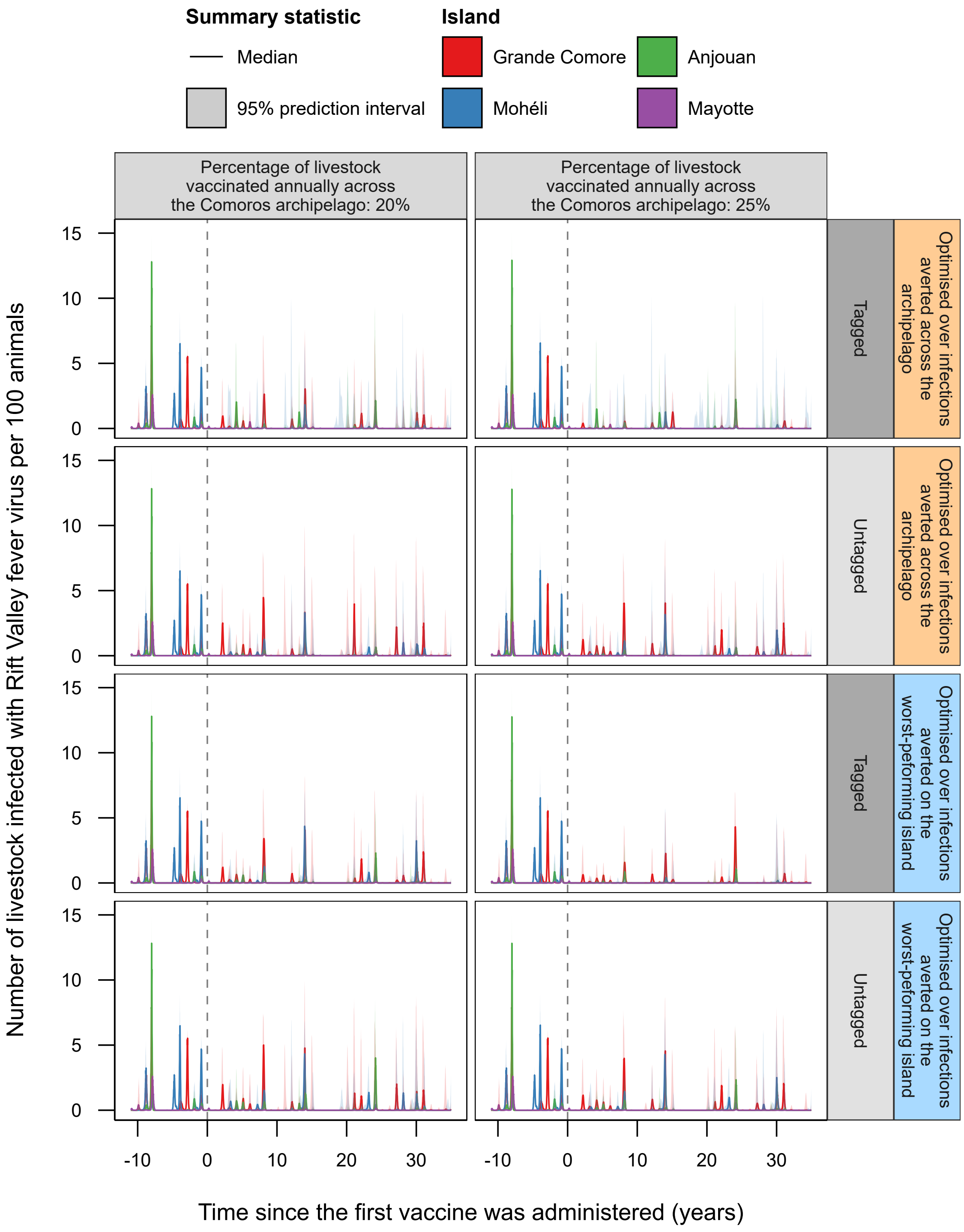


**Supplementary Figure 5: Model predicted number of infected livestock per island in the Comoros archipelago with 20% and 25% vaccination rates.** The fitted metapopulation model describing Rift Valley Fever virus (RVFV) infection in livestock on each island of the Comoros archipelago—Grande Comore (red), Mohéli (blue), Anjouan (green) and Mayotte (purple)—was simulated forward in time for 35 years from the end of the fitting period, June 2015, under a range of vaccination strategies. Shown is the median (solid line) and 95% prediction interval (light area) of model predicted number of infections on each island for 20% and 25% of livestock vaccinated annually across the archipelago, optimal vaccine allocation across islands, and two livestock tagging strategies. This was compared to the model predicted number of infections without vaccination and then used to establish the effectiveness of a given strategy. The median and 95% prediction intervals were calculated from 1000 model simulations.

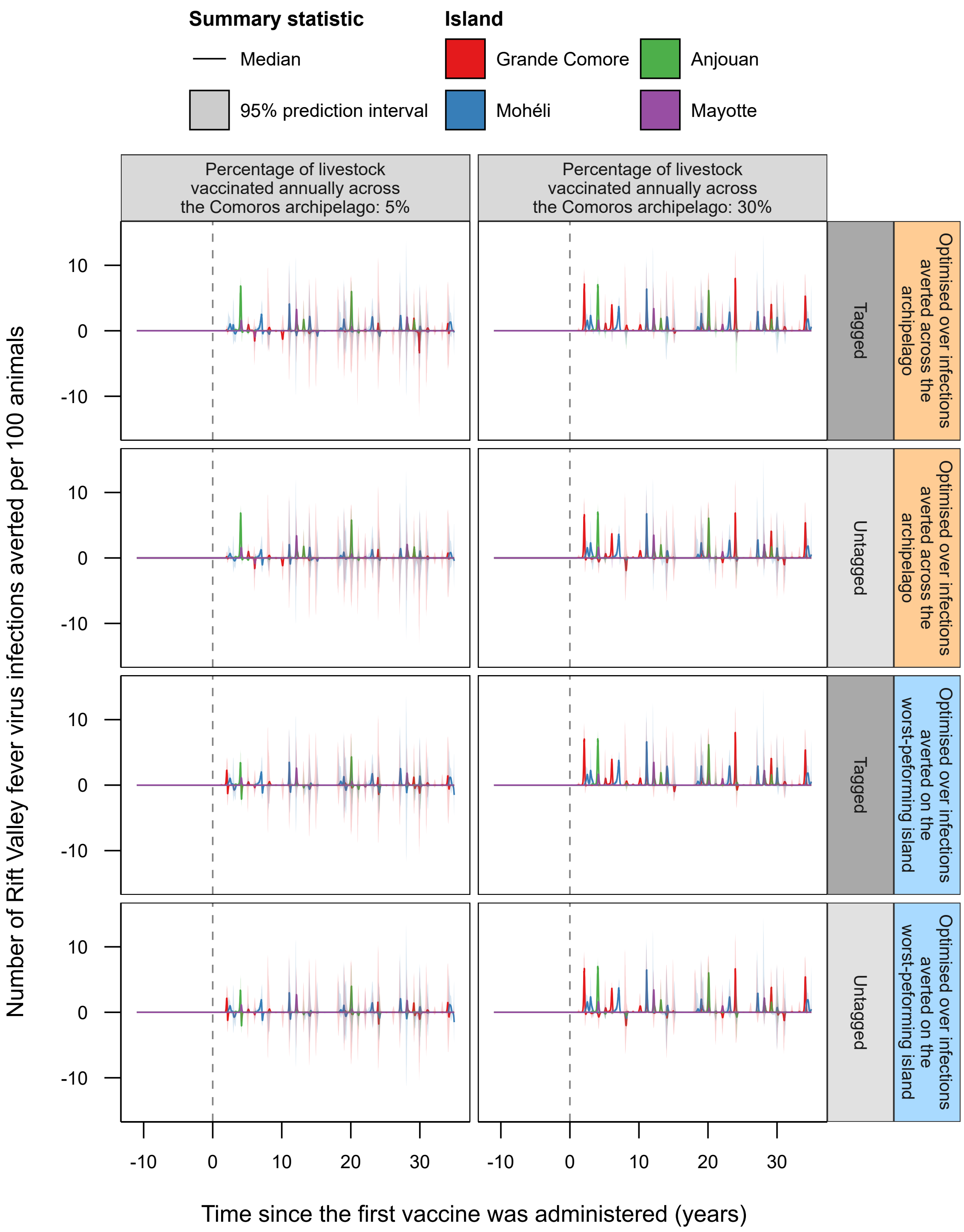


**Supplementary Figure 6: Model predicted number of infected averted livestock per island in the Comoros archipelago with 5% and 30% vaccination rates.** The fitted metapopulation model describing Rift Valley Fever virus (RVFV) infection in livestock on each island of the Comoros archipelago—Grande Comore (red), Mohéli (blue), Anjouan (green) and Mayotte (purple)—was simulated forward in time for 35 years from the end of the fitting period, June 2015, under a range of vaccination strategies. This was then compared with the model predicted number of infections without any vaccination over the same time period. Shown is the median (solid line) and 95% prediction interval (light area) of model predicted number of infections averted on each island for 5% and 30% of livestock vaccinated annually across the archipelago, optimal vaccine allocation across islands, and two livestock tagging strategies. The median and 95% prediction intervals were calculated from 1000 model simulations.

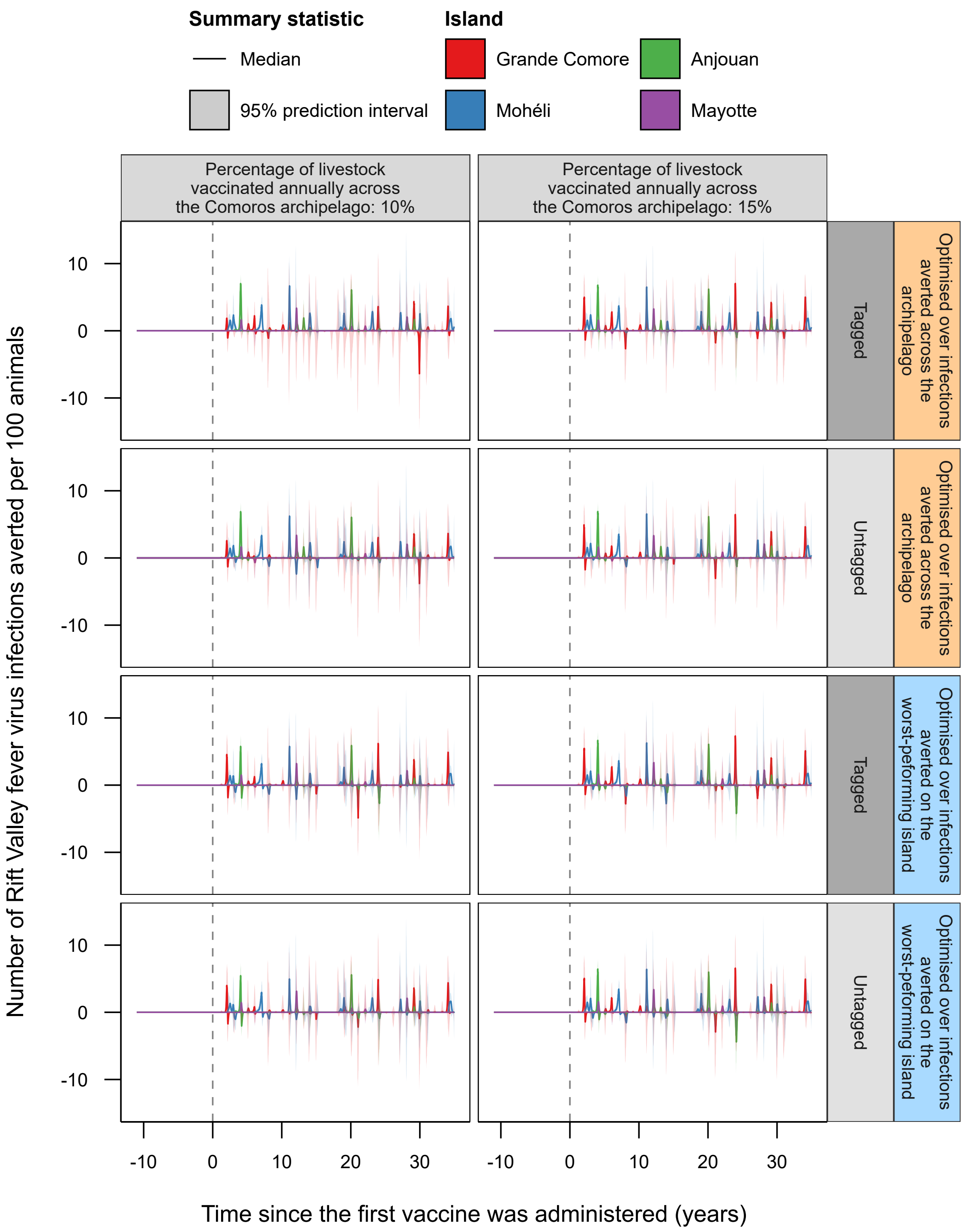


**Supplementary Figure 7: Model predicted number of infected averted livestock per island in the Comoros archipelago with 10% and 15% vaccination rates.** The fitted metapopulation model describing Rift Valley Fever virus (RVFV) infection in livestock on each island of the Comoros archipelago—Grande Comore (red), Mohéli (blue), Anjouan (green) and Mayotte (purple)—was simulated forward in time for 35 years from the end of the fitting period, June 2015, under a range of vaccination strategies. This was then compared with the model predicted number of infections without any vaccination over the same time period. Shown is the median (solid line) and 95% prediction interval (light area) of model predicted number of infections averted on each island for 10% and 15% of livestock vaccinated annually across the archipelago, optimal vaccine allocation across islands, and two livestock tagging strategies. The median and 95% prediction intervals were calculated from 1000 model simulations.

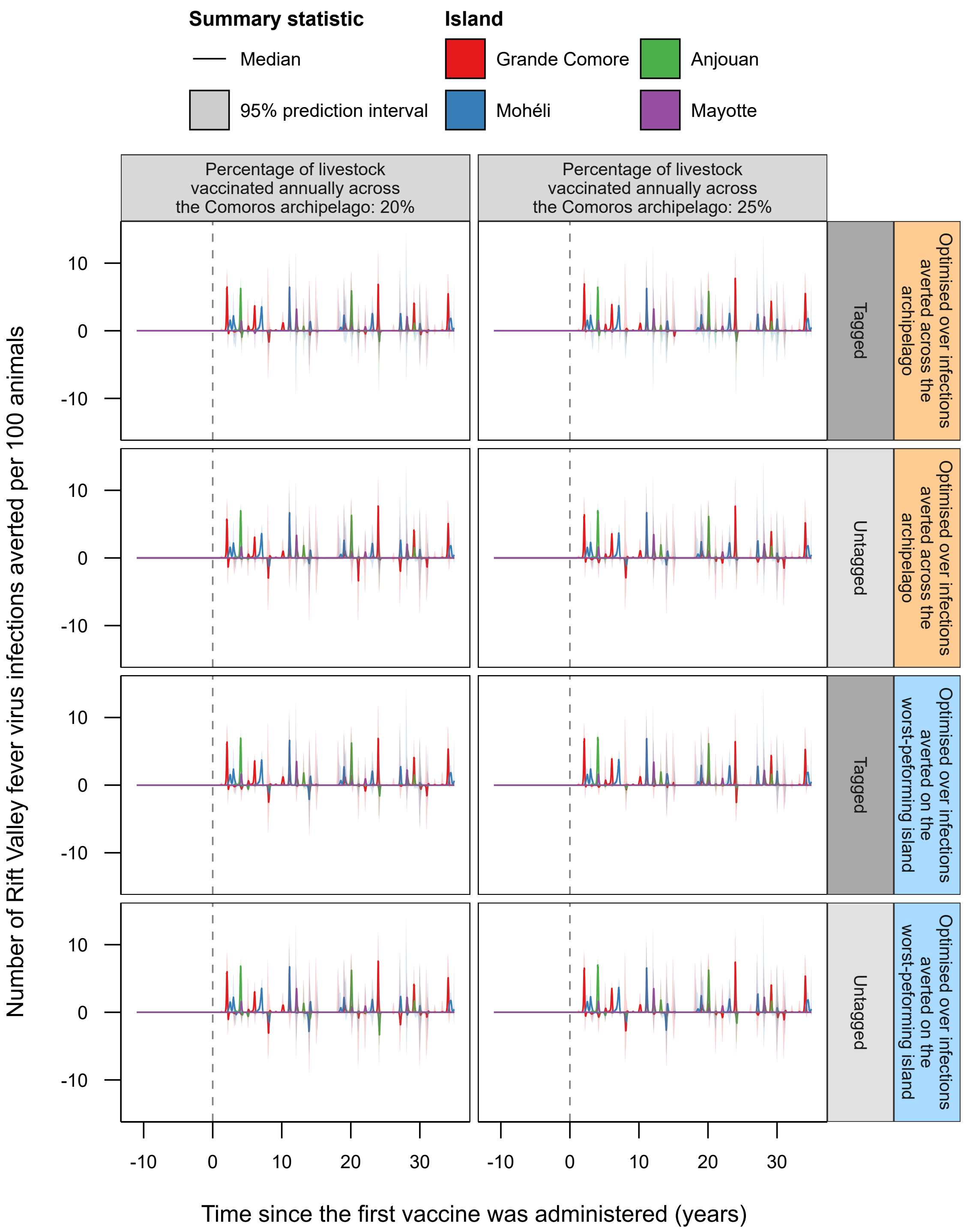


**Supplementary Figure 8: Model predicted number of infected averted livestock per island in the Comoros archipelago with 20% and 25% vaccination rates.** The fitted metapopulation model describing Rift Valley Fever virus (RVFV) infection in livestock on each island of the Comoros archipelago—Grande Comore (red), Mohéli (blue), Anjouan (green) and Mayotte (purple)—was simulated forward in time for 35 years from the end of the fitting period, June 2015, under a range of vaccination strategies. This was then compared with the model predicted number of infections without any vaccination over the same time period. Shown is the median (solid line) and 95% prediction interval (light area) of model predicted number of infections averted on each island for 20% and 25% of livestock vaccinated annually across the archipelago, optimal vaccine allocation across islands, and two livestock tagging strategies. The median and 95% prediction intervals were calculated from 1000 model simulations.

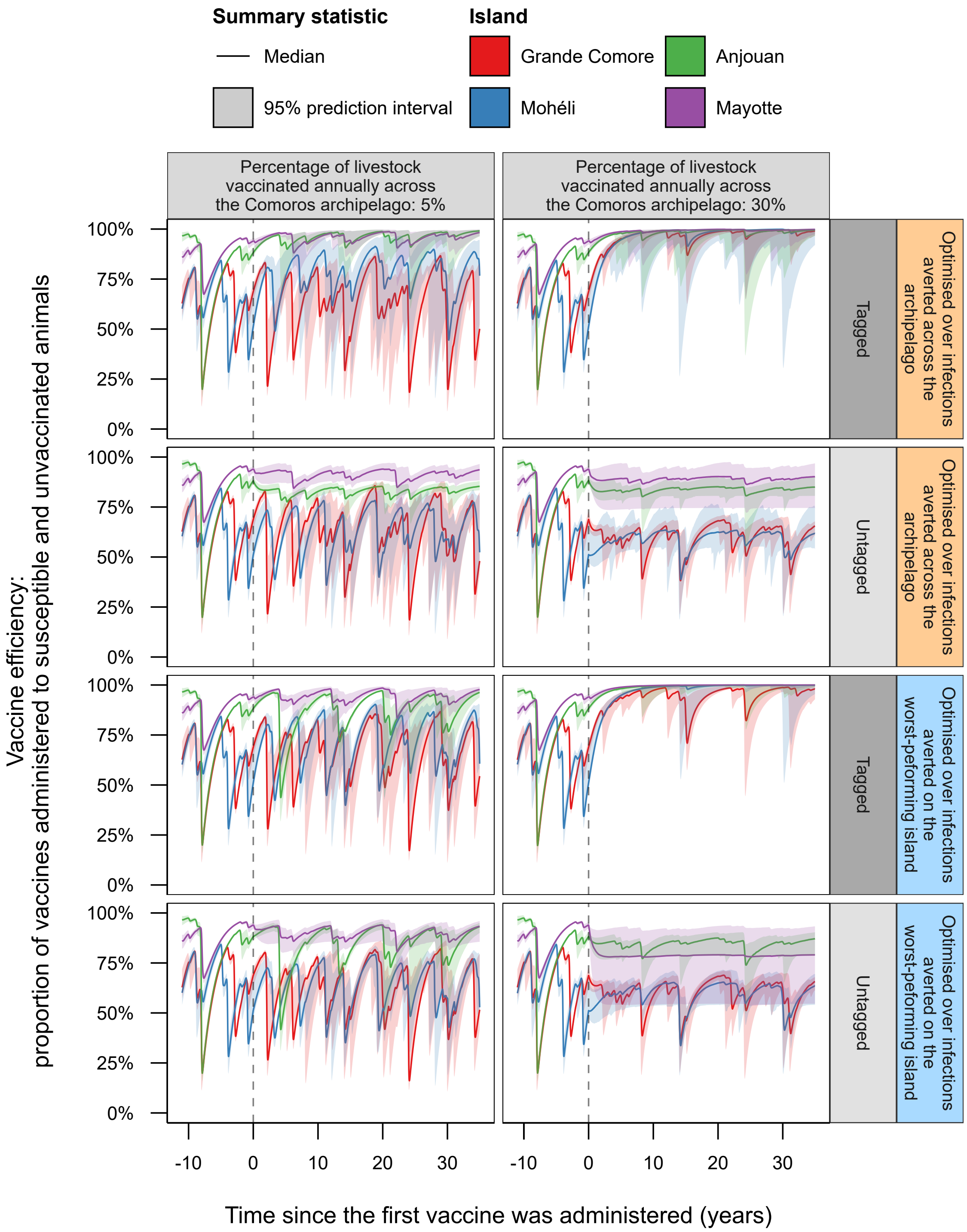


**Supplementary Figure 9: Efficiency of vaccine administration on each island in the Comoros archipelago at 5% and 30% vaccination rates.**. Shown is the weekly model predicted median (solid line) and 95% (shaded area) prediction interval of the percentage of vaccines that were administered to livestock without vaccine-induced nor natural protection against RVFV on each island in the Comoros archipelago—Grande Comore (red), Mohéli (blue), Anjouan (green) and Mayotte (purple)—when vaccinating 5% and 30% of livestock across the archipelago annually under both optimal vaccine allocations and tagging strategies. Summary metrics were generated using 1,000 model simulations.

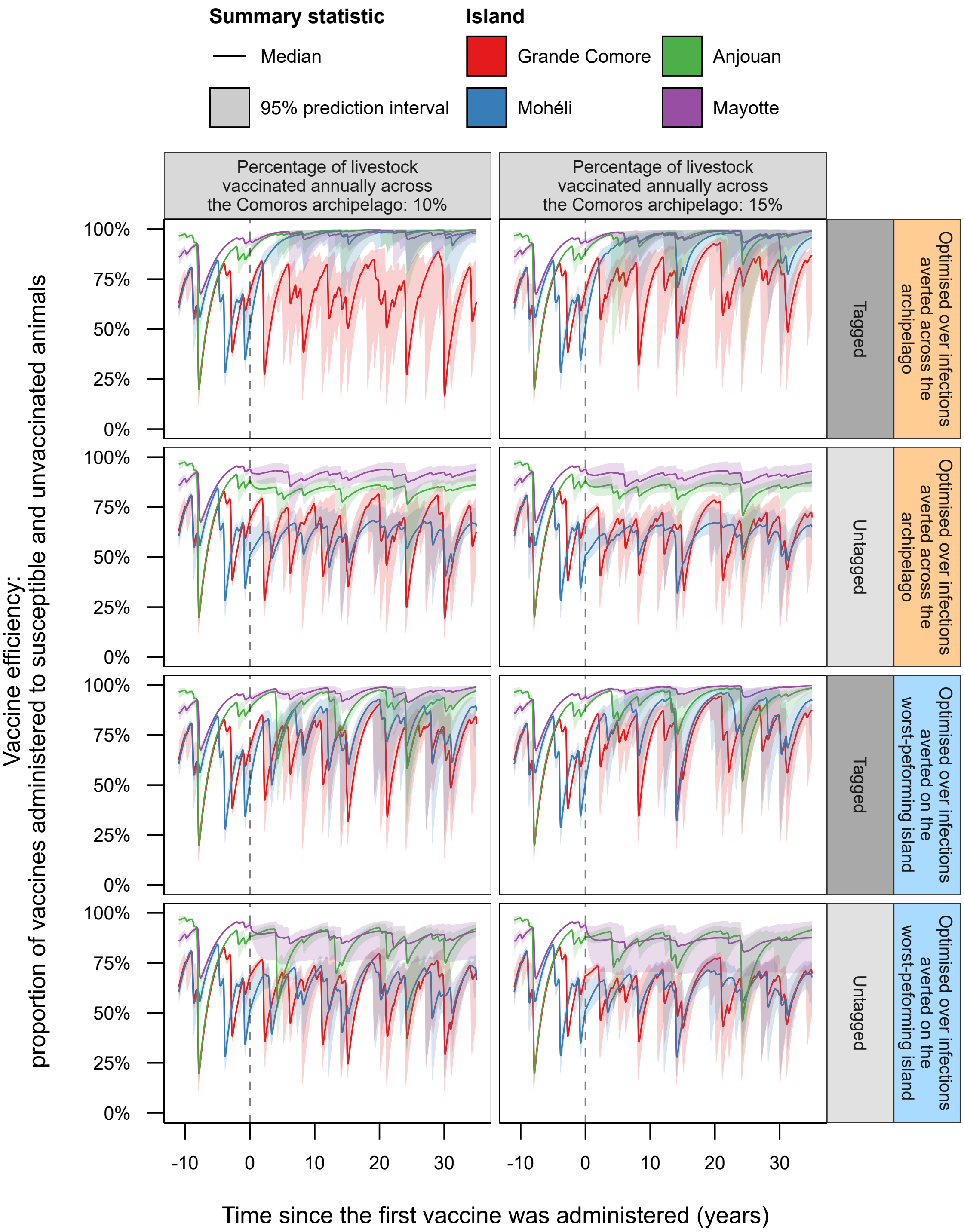


**Supplementary Figure 10: Efficiency of vaccine administration on each island in the Comoros archipelago at 10% and 15% vaccination rates.**. Shown is the weekly model predicted median (solid line) and 95% (shaded area) prediction interval of the percentage of vaccines that were administered to livestock without vaccine-induced nor natural protection against RVFV on each island in the Comoros archipelago—Grande Comore (red), Mohéli (blue), Anjouan (green) and Mayotte (purple)—when vaccinating 10% and 15% of livestock across the archipelago annually under both optimal vaccine allocations and tagging strategies. Summary metrics were generated using 1,000 model simulations.

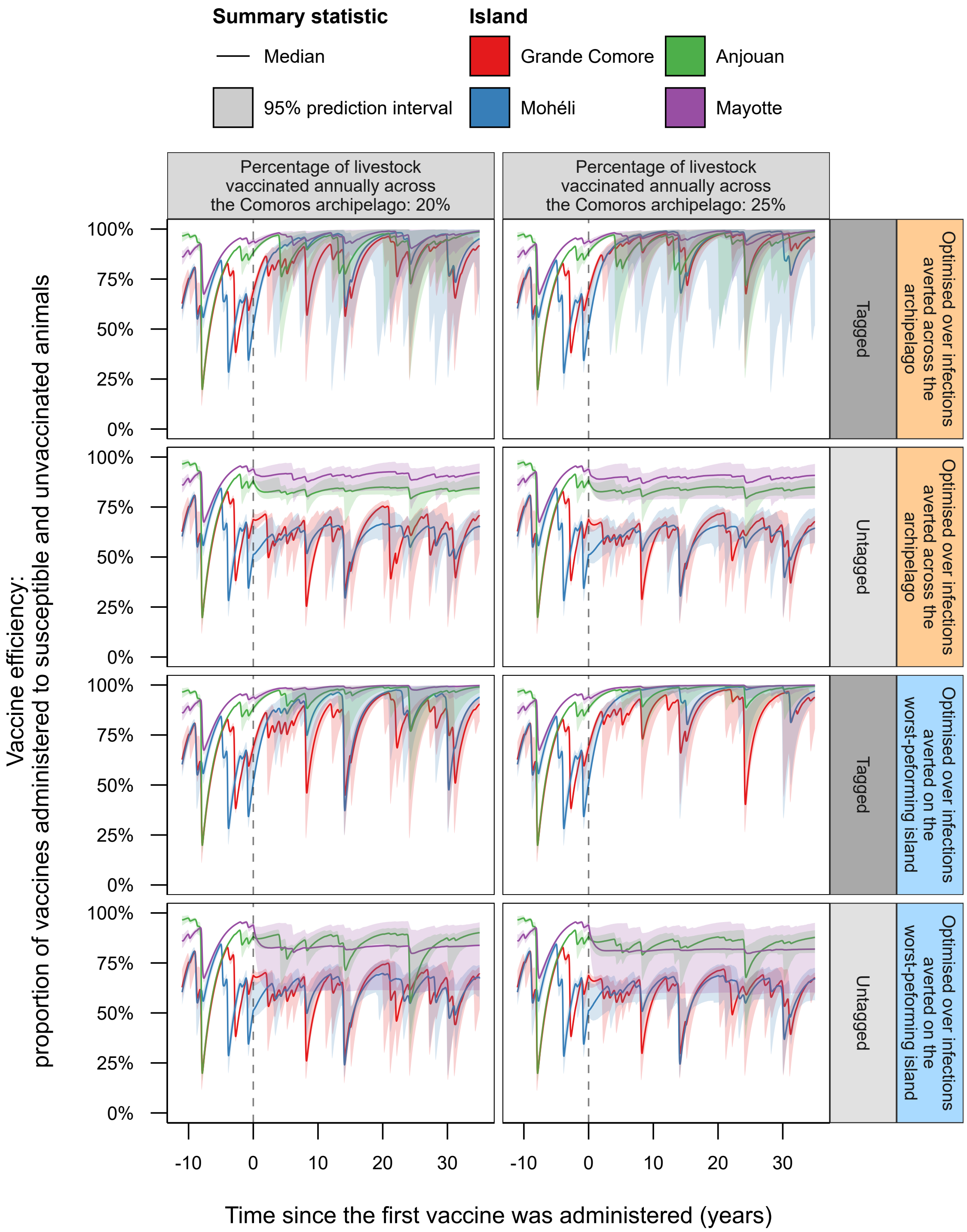


**Supplementary Figure 11: Efficiency of vaccine administration on each island in the Comoros archipelago at 20% and 25% vaccination rates.**. Shown is the weekly model predicted median (solid line) and 95% (shaded area) prediction interval of the percentage of vaccines that were administered to livestock without vaccine-induced nor natural protection against RVFV on each island in the Comoros archipelago—Grande Comore (red), Mohéli (blue), Anjouan (green) and Mayotte (purple)—when vaccinating 20% and 25% of livestock across the archipelago annually under both optimal vaccine allocations and tagging strategies. Summary metrics were generated using 1,000 model simulations.

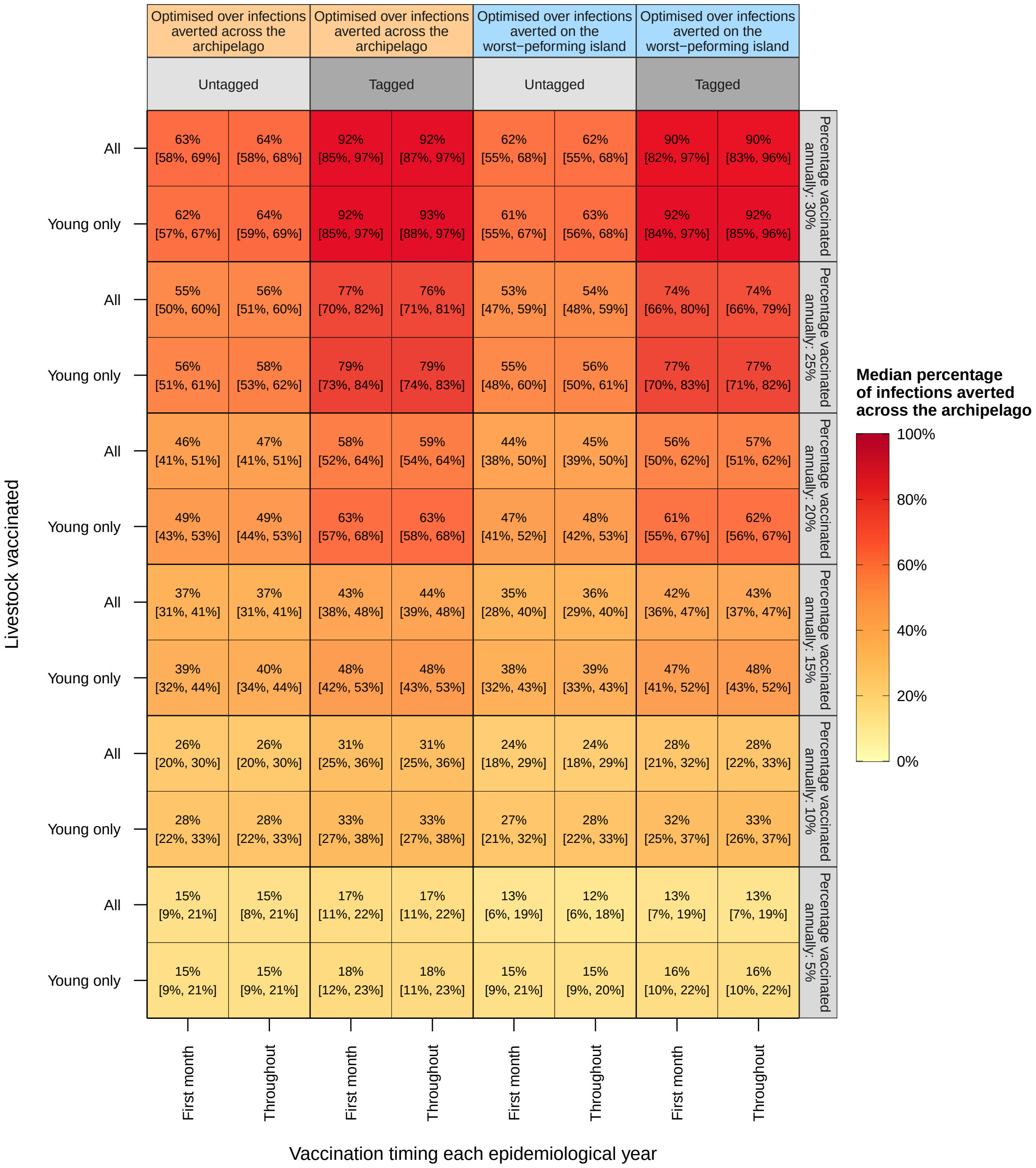


**Supplementary Figure 12: Effectiveness of targeting vaccine efforts to young livestock and only administering vaccines in the first epidemiological month of the year.** The allocation of vaccines were optimised for each vaccination rate and tagging strategy assuming that vaccines were administered across all age groups and throughout the epidemiological year. Using these optimal vaccine allocation, the effectiveness of only vaccinating the first two age groups and / or within the first epidemiological month (July) were simulated using the model. Shown is the median and 95% prediction interval of the percentage of infections averted across the Comoros archipelago for each scenario. All metrics shown were based on 25,000 model simulations.

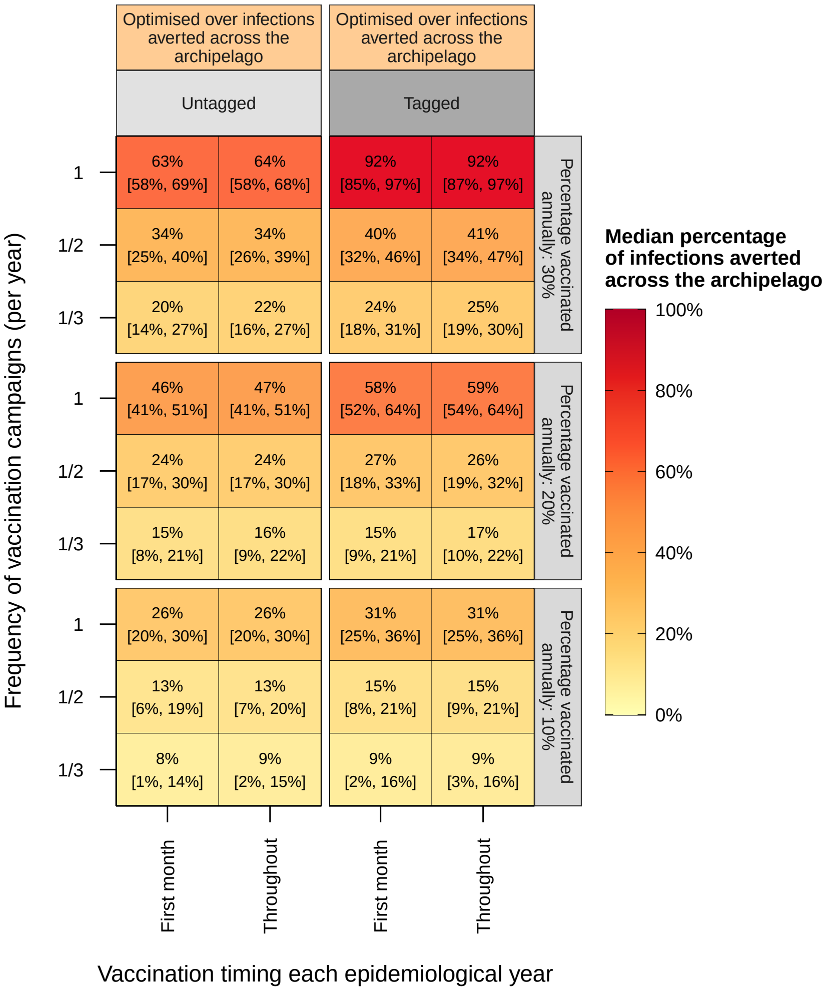


**Supplementary Figure 13: Effectiveness with altered timing and frequency of vaccine administration.** The allocation of vaccines were optimised for each vaccination rate and tagging strategy assuming that vaccines were administered across all age groups and throughout the epidemiological year. Using these optimal vaccine allocation, the effectiveness of only vaccinating every two or three years, alongside only vaccinating within the first epidemiological month (July), were simulated using the model for a reduced set of vaccination rates. Shown is the median and 95% prediction interval of the percentage of infections averted across the Comoros archipelago for each scenario. All metrics shown were based on 25,000 model simulations.

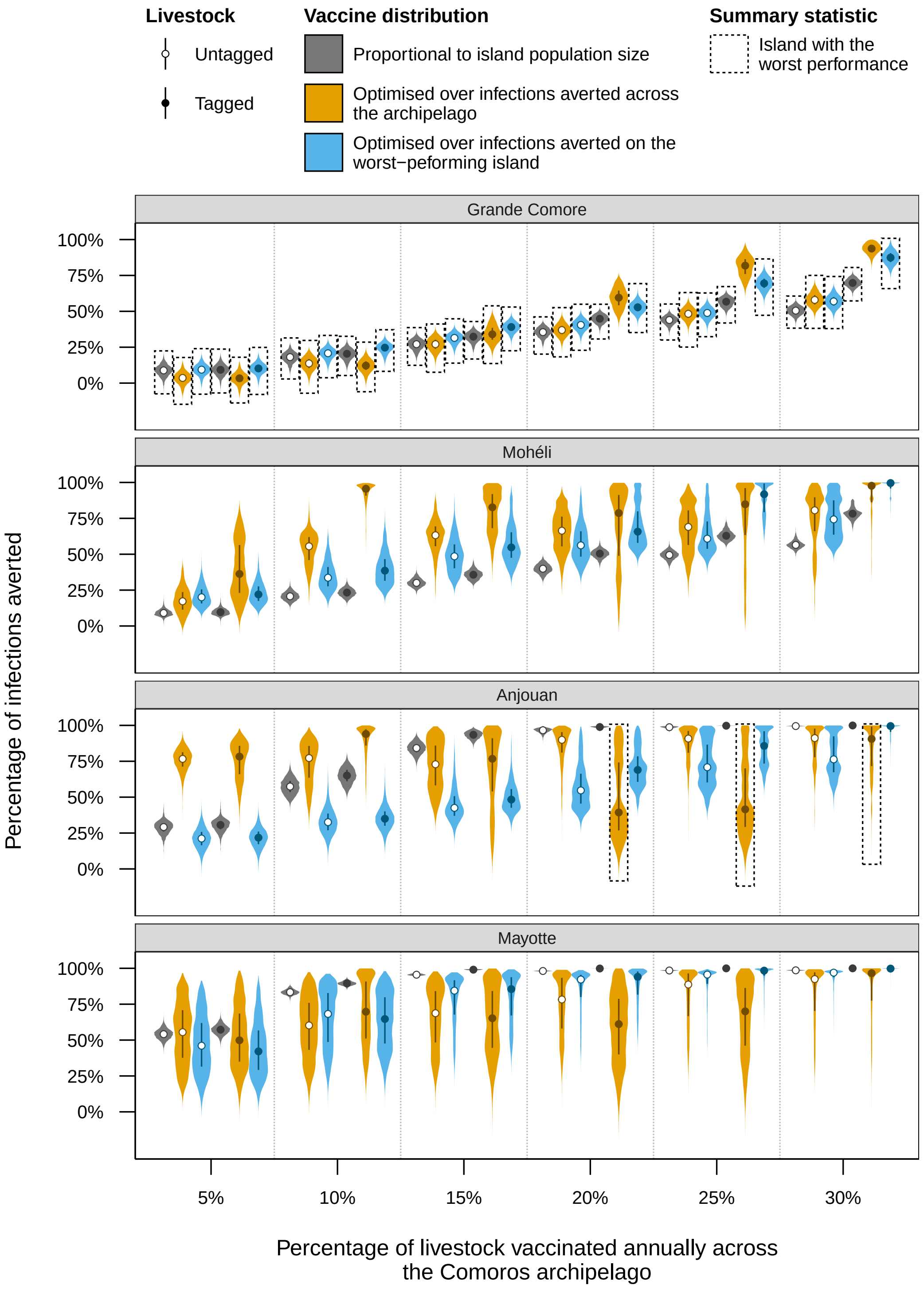


**Supplementary Figure 14: Effectiveness of different vaccine strategies against Rift Valley fever virus (RVFV) on each island in the Comoros archipelago.** For six annual vaccination rates, vaccines were allocated to islands in the Comoros archipelago either proportionally to the population size of each island (grey violins), optimally in terms of infections averted across the archipelago (orange violins) or optimmaly in terms of infections averted on the worst-performing island (blue). Livestock were also either tagged post-vaccination (black circles) or not (white circles). For the majority of vaccination rates, vaccine allocations and tagging strategies, Grande Comore averted the lowest percentage of infections on average (grey boxes), and tagging livestock resulted in a greater number of infections averted across the archipelago. The violins show the percentage of infections averted across on each island for different annual vaccination rates, allocation methods and tagging strategies. The points and boxplots show the median and inter-quartile range for each scenario respectively. All metrics shown were based on 25,000 model simulations.

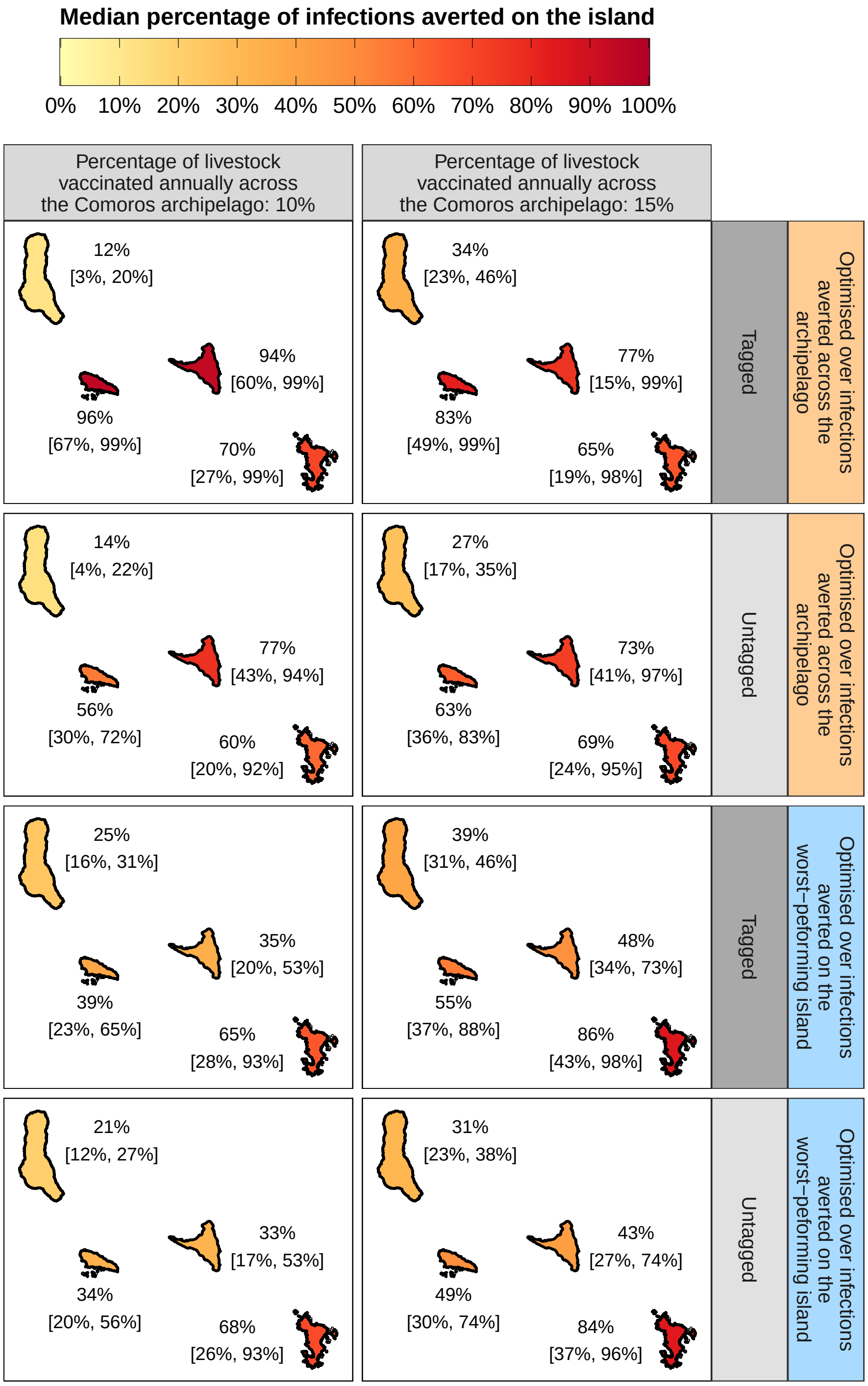


**Supplementary Figure 15: Equity in vaccine strategy effectiveness between islands in the Comoros archipelago for 10% and 15% vaccination rates.** Vaccines were allocated between islands in the Comoros archipelago either optimally in terms of total infections averted across the archipelago (orange panels) or optimally in terms of infections averted on the worst performing island (blue panels; where the worst-performing island was defined as the island with the lowest percentage of infections averted). Shown is the median and 95% prediction interval of the percentage of infections averted on each island when vaccinating 10% and 15% of livestock across the archipelago annually under each optimal vaccine allocation and tagging strategy. Summary statistics were generated using 25,000 model simulations. See Supplementary Tables 6 and 7 for detailed numerical results.

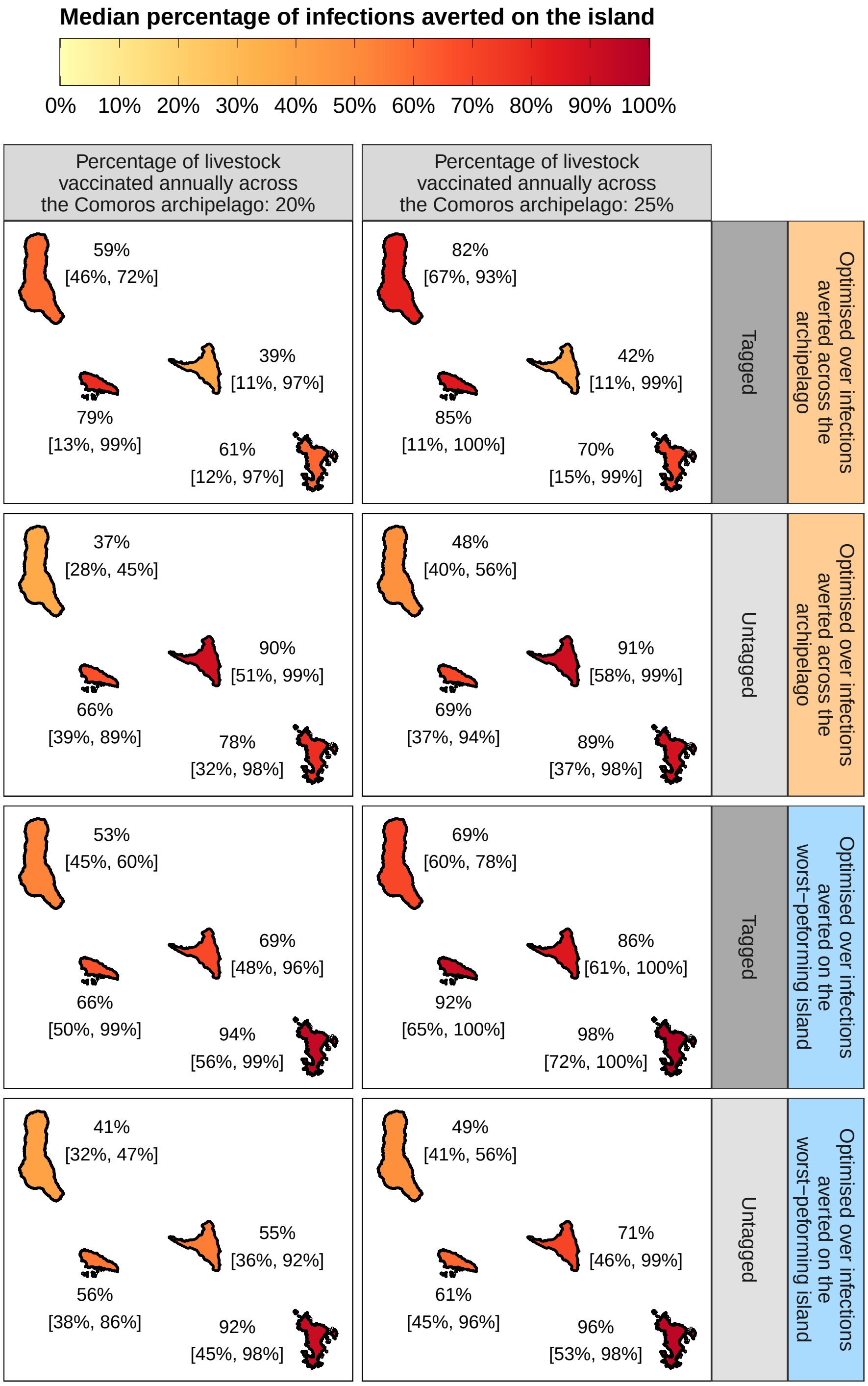


**Supplementary Figure 16: Equity in vaccine strategy effectiveness between islands in the Comoros archipelago for 20% and 25% vaccination rates.** Vaccines were allocated between islands in the Comoros archipelago either optimally in terms of total infections averted across the archipelago (orange panels) or optimally in terms of infections averted on the worst performing island (blue panels; where the worst-performing island was defined as the island with the lowest percentage of infections averted). Shown is the median and 95% prediction interval of the percentage of infections averted on each island when vaccinating 20% and 25% of livestock across the archipelago annually under each optimal vaccine allocation and tagging strategy. Summary statistics were generated using 25,000 model simulations. See Supplementary Tables 6 and 7 for detailed numerical results.

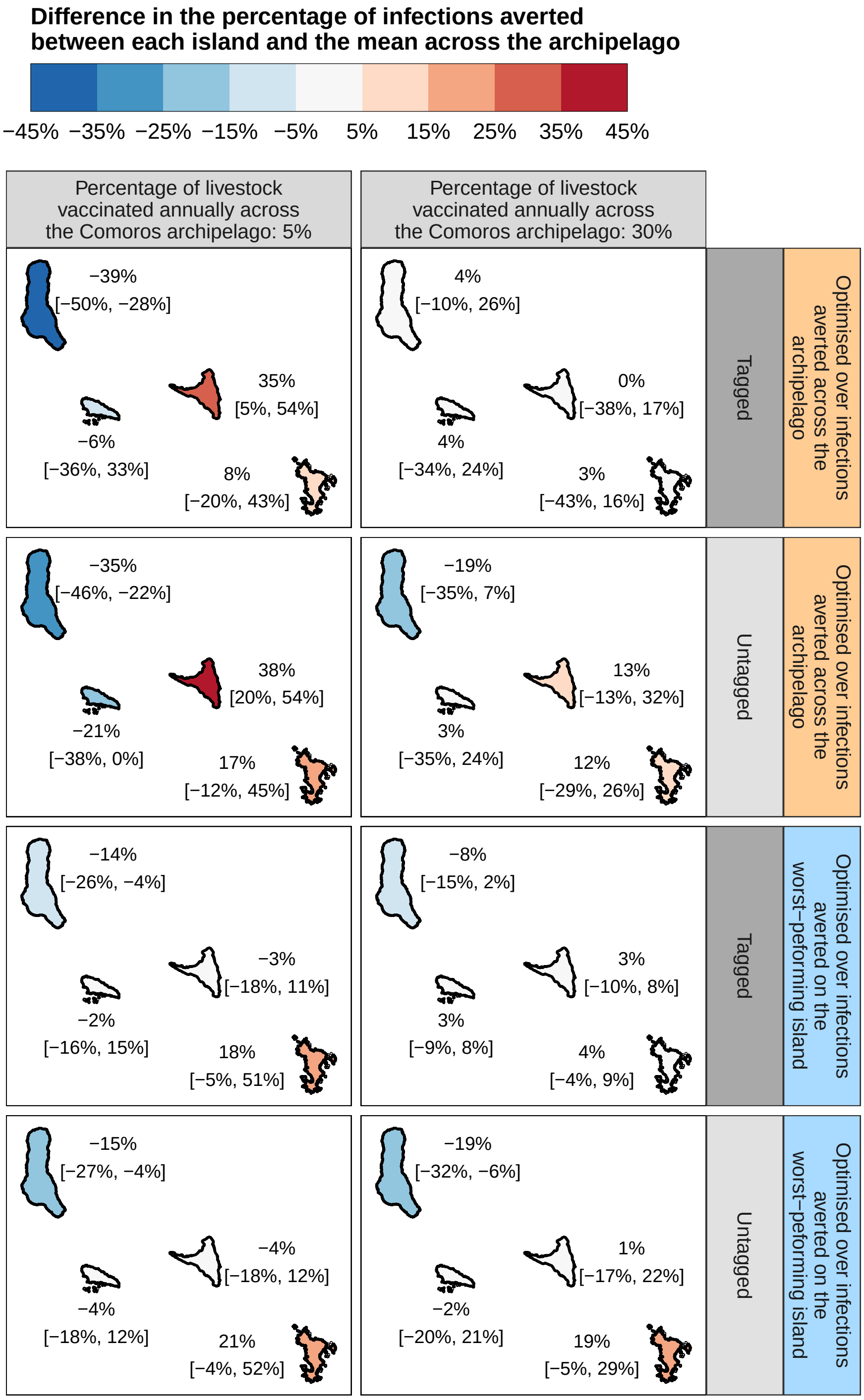


**Supplementary Figure 17: Difference in vaccine strategy effectiveness between islands and the mean across the Comoros archipelago for 5% and 30% vaccination rates.** Vaccines were allocated between islands in the Comoros archipelago either optimally in terms of total infections averted across the archipelago (orange panels) or optimally in terms of infections averted on the worst performing island (blue panels; where the worst-performing island was defined as the island with the lowest percentage of infections averted). Shown is the median and 95% prediction interval of the percentage point difference between the percentage of infections averted on each island and the mean percentage of infections averted across the archipelago, with 5% and 30% of livestock across the archipelago vaccinated annually under each optimal vaccine allocation and tagging strategy. Summary statistics were generated using 25,000 model simulations. See Supplementary Table 8 for detailed numerical results.

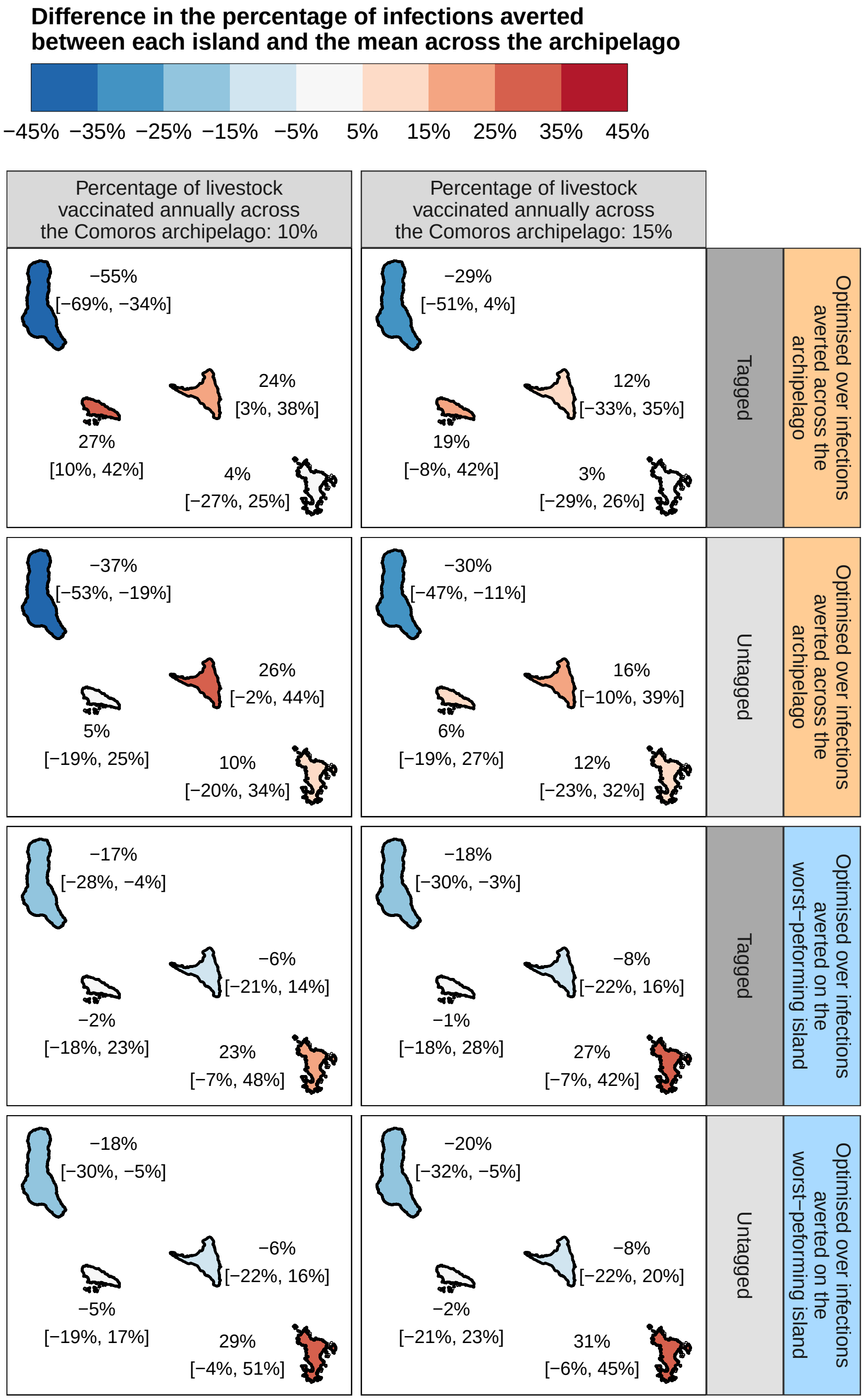


**Supplementary Figure 18: Difference in vaccine strategy effectiveness between islands and the mean across the Comoros archipelago for 10% and 15% vaccination rates.** Vaccines were allocated between islands in the Comoros archipelago either optimally in terms of total infections averted across the archipelago (orange panels) or optimally in terms of infections averted on the worst performing island (blue panels; where the worst-performing island was defined as the island with the lowest percentage of infections averted). Shown is the median and 95% prediction interval of the percentage point difference between the percentage of infections averted on each island and the mean percentage of infections averted across the archipelago, with 10% and 15% of livestock across the archipelago vaccinated annually under each optimal vaccine allocation and tagging strategy. Summary statistics were generated using 25,000 model simulations. See Supplementary Table 8 for detailed numerical results.

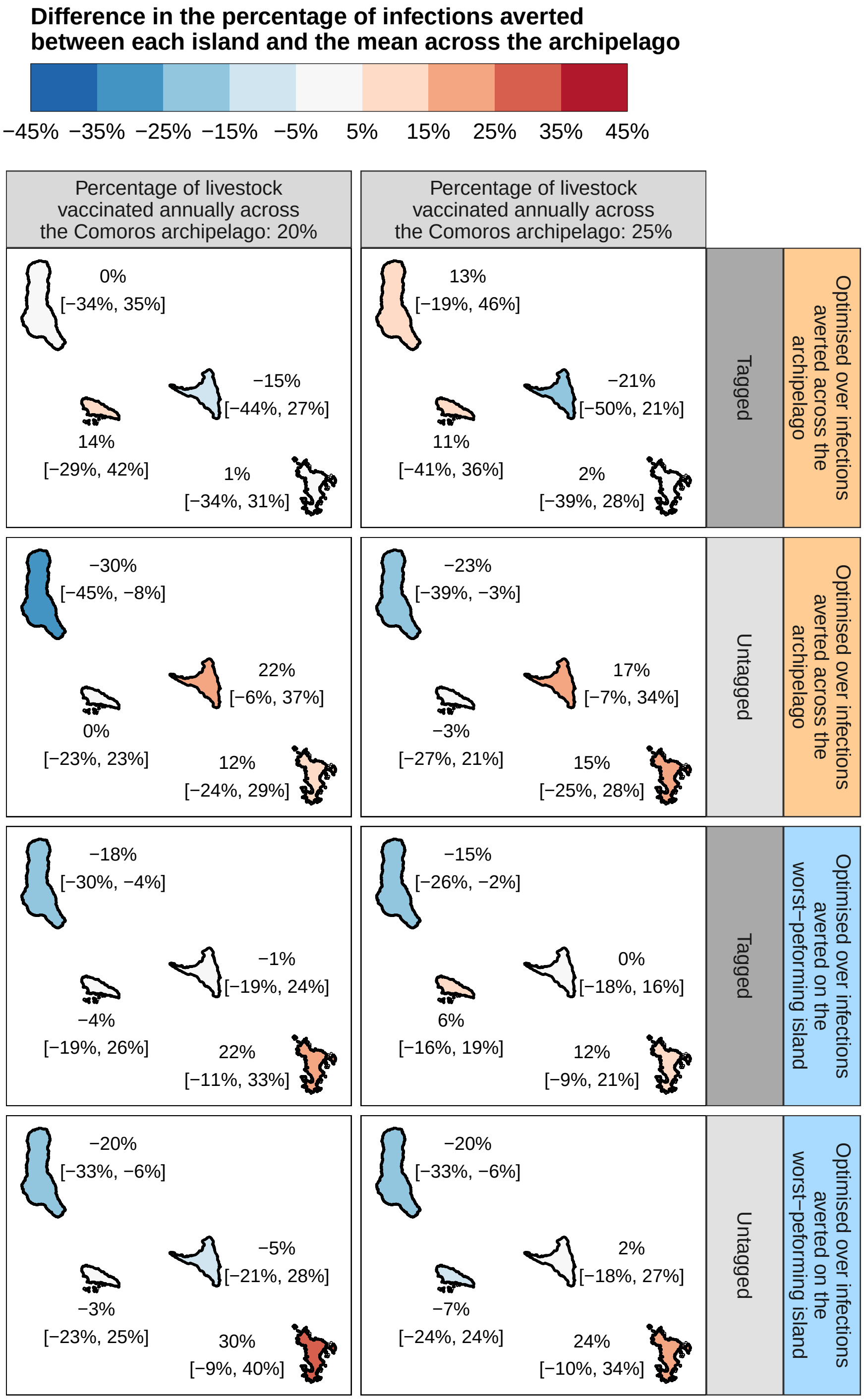


**Supplementary Figure 19: Difference in vaccine strategy effectiveness between islands and the mean across the Comoros archipelago for 20% and 25% vaccination rates.** Vaccines were allocated between islands in the Comoros archipelago either optimally in terms of total infections averted across the archipelago (orange panels) or optimally in terms of infections averted on the worst performing island (blue panels; where the worst-performing island was defined as the island with the lowest percentage of infections averted). Shown is the median and 95% prediction interval of the percentage point difference between the percentage of infections averted on each island and the mean percentage of infections averted across the archipelago, with 20% and 25% of livestock across the archipelago vaccinated annually under each optimal vaccine allocation and tagging strategy. Summary statistics were generated using 25,000 model simulations. See Supplementary Table 8 for detailed numerical results.

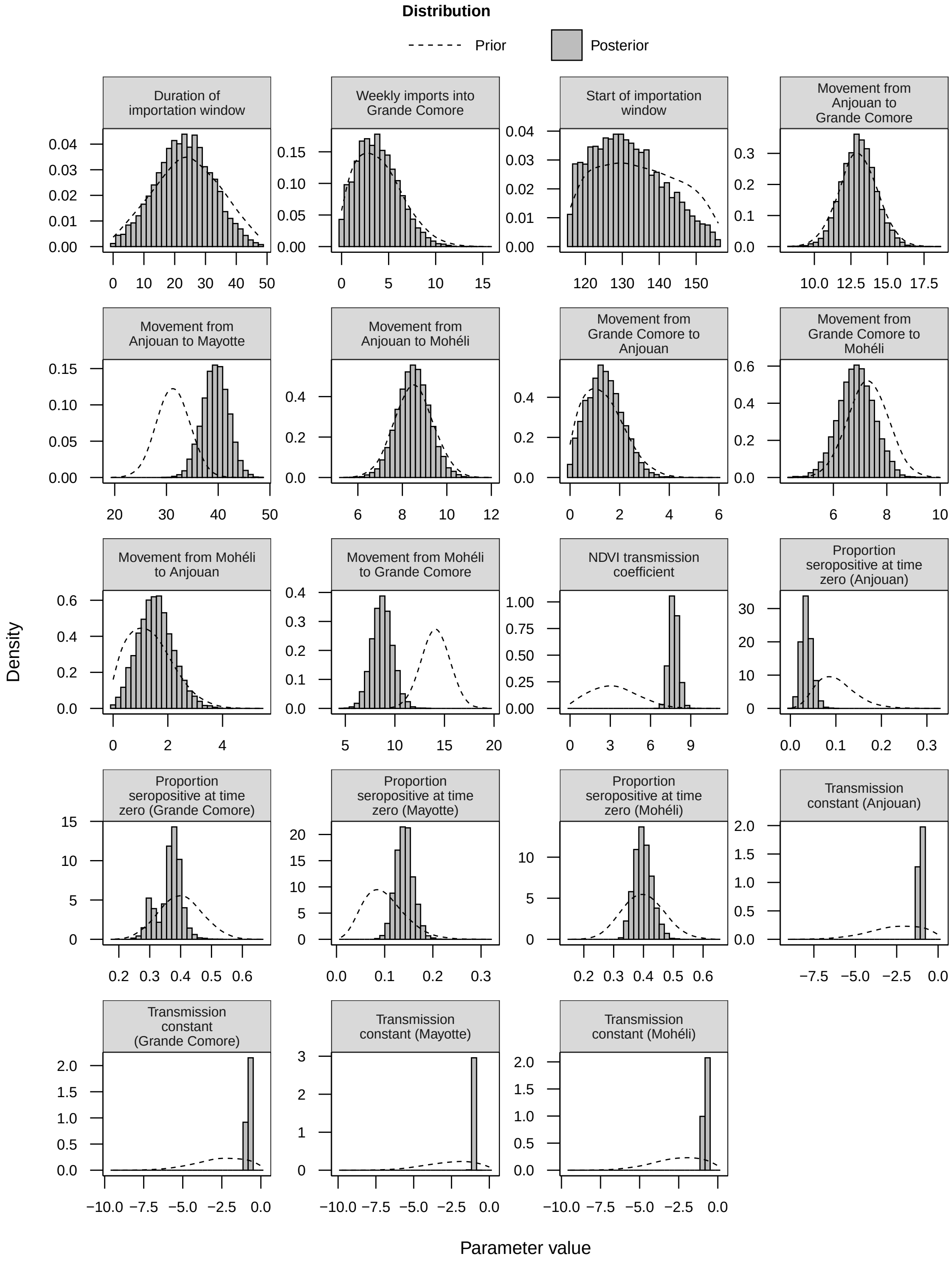


**Supplementary Figure 20: Posterior distribution of model parameters.** We fitted our transmission model—in the absence of vaccination—to age-stratified cross-sectional and longitudinal seroprevalence surveys conducted throughout the Comoros archipelago from 2004 until 2015. By fitting to these data in a Bayesian framework, we used a random-walk Metropolis-Hastings Markov Chain Monte Carlo algorithm to estimate parameters of interest within the model. The histograms show the marginal posterior distribution of each parameter of interest. The dashed lines depict the prior distribution of each parameter. The marginal posterior distributions were generated using 10,000 samples from the joint posterior distribution of parameters. For further details on the model fitting procedure, refer to Tennant et al. [1].
